# Time-resolved phenotyping at subcellular resolution reveals shared principles and key trade-offs across antimicrobial peptide activities

**DOI:** 10.1101/2025.04.10.648262

**Authors:** Alessio Fragasso, Tatjana Schlechtweg, Wei-Hsiang Lin, Annelise E. Barron, Christine Jacobs-Wagner

**Affiliations:** Sarafan ChEM-H Institute, Stanford University, Stanford, CA, USA; Department of Biology, Stanford University, Stanford, CA, USA; Howard Hughes Medical Institute, Stanford University, Stanford, CA, USA; Department of Bioengineering, Stanford University, Schools of Medicine and of Engineering, Stanford, CA, USA; Department of Microbiology and Immunology, Stanford University, School of Medicine, Stanford, CA, USA

## Abstract

Cationic antimicrobial peptides are a broad family of host defense molecules that neutralize bacteria by permeabilizing one or more membranes and/or inhibiting intracellular targets. Here, we present a time-resolved single-cell pipeline for quantifying these effects in *Escherichia coli*. Applying this pipeline to 18 diverse natural peptides and synthetic peptidomimetics reveals shared core activities, but with different kinetics, defining two classes with opposite trade-offs. Class I peptides cause abrupt growth arrest, predominantly coupled with inner membrane permeabilization and ribosome/DNA reorganization, conferring fast, multipronged action. However, rapid intracellular absorption by the first permeabilized cells depletes the extracellular pool, rendering them ineffective against dense populations, including biofilms. Class II peptides act more gradually, with delayed or absent inner membrane permeabilization, limiting their speed of action. However, this results in slower intracellular absorption and greater efficacy at high cell densities and against biofilms. These opposing functional trade-offs point to important immunological and therapeutic implications.

## INTRODUCTION

Antimicrobial peptides (AMPs) constitute a widespread group of host defense peptides found across all domains of life, from bacteria to plants and animals, including humans (Wang et al., 2016, 2022). These naturally occurring peptides have garnered significant interest due to their multimodal role in innate immunity, where they serve as both antimicrobial agents and immunomodulators (Hancock et al., 2016; Mookherjee et al., 2020). Furthermore, their broad-spectrum efficacy against bacteria, viruses, fungi, and parasites, coupled with a low tendency for resistance development, has made them attractive candidates for antibiotic alternatives or supplements (Mookherjee et al., 2020; Taheri-Araghi, 2024). Consequently, significant efforts have been dedicated to developing synthetic AMPs and peptidomimetics (PMs) to combat the threat of infectious diseases and the rise of antibiotic resistance.

With over 2,500 natural AMPs identified in animals alone (as of December 2024), the sheer diversity of peptide sequence is remarkable (Wang et al., 2016). They also vary in structure, encompassing α-helical forms, β-sheets, β-hairpins, and extended structures. Moreover, some AMPs undergo post-translational modifications (Wang et al., 2016), further contributing to their diversity. Notwithstanding their structural differences, many AMPs share two key physicochemical features: they are typically amphipathic and cationic at physiological pH. These properties enable AMPs to interact with bacterial cytoplasmic membranes, which are negatively charged in part due to the prevalence of anionic phospholipids. Gram-negative bacteria possess an additional outer membrane typically rich in highly anionic lipopolysaccharides (LPS), which provides another target for cationic AMPs.

The mechanisms of action for AMPs can be broadly categorized based on their interaction with the membrane (Hale & Hancock, 2007). The first, most common killing mechanism involves compromising the physical integrity of the cytoplasmic membrane, which can be achieved via various strategies (Wimley & Hristova, 2011). In Gram-negative bacteria, membrane-disruptive peptides are thought to promote their own access to the cytoplasmic membrane by creating holes in the outer membrane barrier in a process called “self-promoted uptake” (Hancock, 1997). However, subsequent permeabilization of the cytoplasmic membrane is not necessarily the endpoint. Evidence suggests that this step can facilitate the influx of peptides into the cytoplasm, where they can interact with anionic targets such as DNA, RNAs, and ribosomes (Cardoso et al., 2019; Luo & Song, 2021), inducing aggregation or cytoplasmic phase separation (Boeynaems et al., 2023; Chongsiriwatana et al., 2017; Sneideris et al., 2023; Zhu et al., 2019). The second, less common mechanism involves inhibiting intracellular targets without permeabilizing the membranes (Hale & Hancock, 2007; Scocchi et al., 2011). These membrane-inactive AMPs are typically characterized by a high proline content. They cross the cell envelope, often with the aid of cellular transporters, to inhibit essential cytoplasmic functions such as protein synthesis (Huang et al., 2024; Scocchi et al., 2011).

Together, these findings indicate that AMPs can exert diverse cellular effects. Yet, building a unified understanding of these mechanisms has remained challenging for several reasons. First, outside notable exceptions (Choi et al., 2016), AMP studies generally lack the temporal, single-cell, and subcellular resolution needed to precisely order the sequence of membrane disruption and intracellular reorganization events relative to the growth arrest, making it difficult to discriminate the primary cause of antibacterial action from downstream pleiotropic effects. Second, studies are rarely performed under standardized conditions; differences in growth media, bacterial strains, and experimental approaches hinder cross-AMP comparisons and the extraction of general principles. Third, under physiological conditions, bacteria often form biofilms, which are dense communities that contribute to many chronic infections (Costerton et al., 1999; Kostakioti et al., 2013). Why the potency against biofilms varies across AMPs is not well understood.

To address these outstanding questions, we developed an integrated experimental and computational pipeline that uses time-lapse fluorescence microscopy and microfluidics. This platform allowed us to quantify, over time, the permeabilization of the outer membrane (OM) and inner membrane (IM) of *Escherichia coli*, as well as the intracellular effects caused by a diverse panel of cationic AMPs and PMs. All experiments — spanning planktonic cells at low and high densities as well as biofilms — were performed under standardized conditions to enable direct cross-compound comparisons. Our comparative analysis reveals that despite their diversity, all examined compounds share a set of core activities, suggesting that their common physicochemical properties — net cationic charge and amphipathicity — likely govern these shared actions. At the same time, marked differences in the kinetics of these activities give rise to two major classes with distinct functional trade-offs that suggest complementary immunological strategies among AMPs and provide new guiding principles for PM design.

## RESULTS

For our study, we selected fourteen natural cationic AMPs that vary widely with respect to their animal source, sequence, structure, net positive charge, and size: bactenecin-7 (Bac7), camel bactenecin (CamBac), cecropin A (CecA), CRAMP, human β-defensin 3 (HBD-3), indolicidin (Indo), LL-37, magainin-2 (Mag2), melittin (Mltt), protegrin-1 (PG-1), PR-39, tachyplesin-1 (Tac1), Tur1A, and Tur1B (Figures 1A-B and S1A). These peptides also vary in their grand average of hydropathy (GRAVY) index and in their fraction of lysine, arginine, proline, polar, and hydrophobic residues (Figures 1B and S1A). Some of the selected AMPs carry modifications such as disulfide bonds or a C-terminal NH_2_ group, further increasing the diversity of our panel (Figure 1A). For each peptide, we determined the MIC for *E. coli* (strain MG1655) grown under identical growth conditions (37°C, M9 minimal medium with glucose, casamino acids, and thiamine, or M9gluCAAT), starting with a low inoculum density of cells (∼250 cells/mL) (Figures 1A and S1B-C). For imaging, we built a simple microwell device (Figure S1D) that contains four independent wells where the growth and division of immobilized *E. coli* cells can be tracked. With this setup, AMPs can be easily added at a defined final concentration, unlike with agarose pads, which are often used for time-lapse imaging of bacterial cells. We chose this setup over popular microfluidic devices (e.g., mother machines) because it requires smaller amounts of AMP, as some of these peptides are costly or difficult to obtain (e.g., due to their chemical modifications).

**Figure 1:**
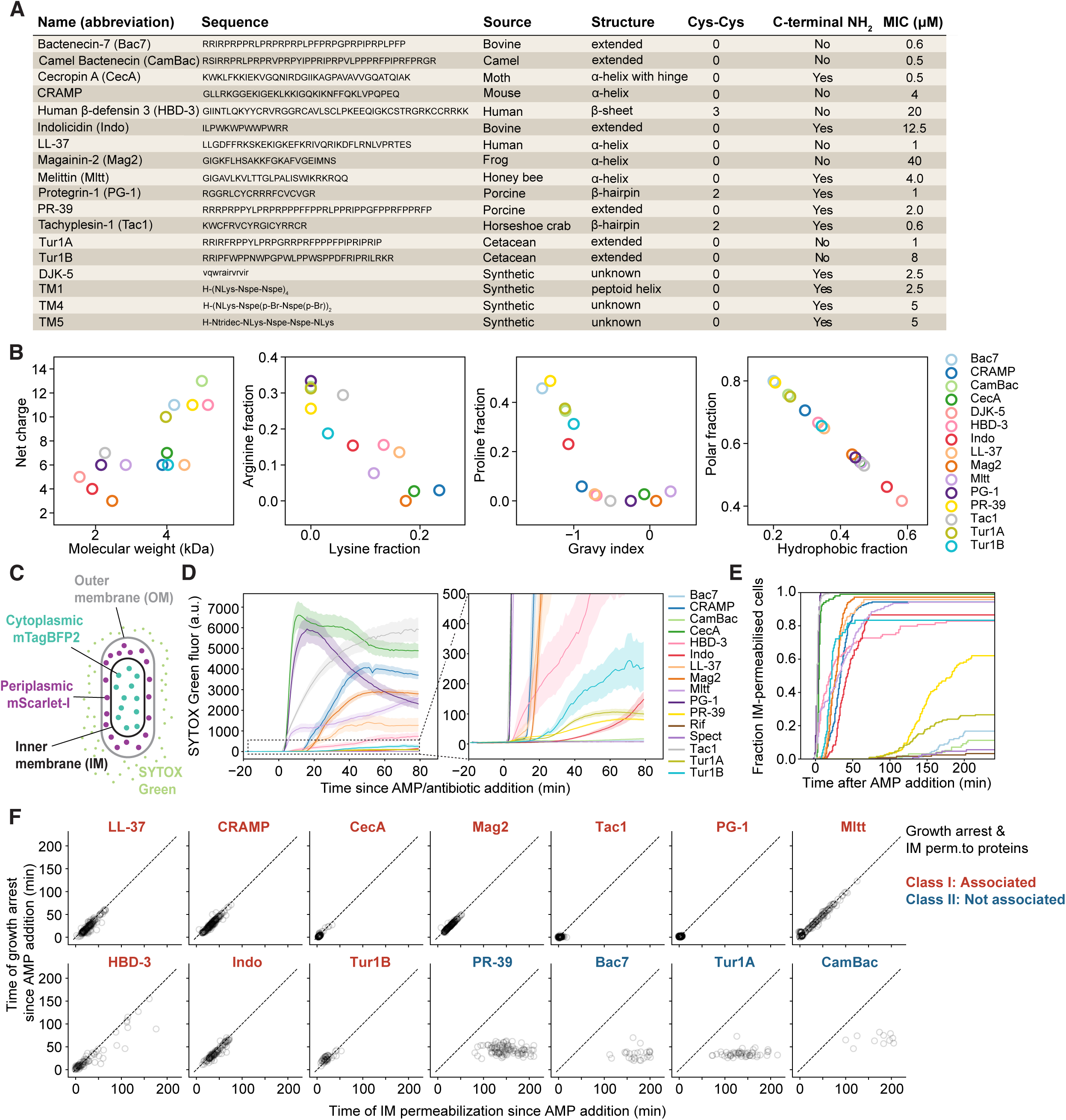
AMPs can be classified based on the phenotypes they cause at the temporal and single-cell resolution. **(A)** List of natural AMPs and synthetic mimics used in this study. Also shown are their features and minimal inhibitory concentrations (MIC) at which no growth of *E. coli* (MG1655) could be detected after incubation in M9gluCAAT at 37°C for 20 h, starting with an inoculum of ∼250 cells/mL (see Figure S1A and & for the MIC results). Some peptides have disulfide bonds (Cys-Cys) and/or an amide group (-NH_2_) at their C-terminal end, as indicated. Peptide structures were obtained from the Protein Data Bank or the AlphaFold Protein Structure Database (PDB: 5HAU, 1KJ6, 1G8C, 5NMN, 2MAG, 2MLT, 2RTV, 6FKR, 1PG1; AlphaFold: AF-P01507-F1, AF-P80054-F1) or predicted using ColabFold v1.6.1 (AlphaFold 2) (Mirdita et al., 2022). The lower-case sequence for the peptide DJK-5 indicates D-amino acids. **(B)** Plots showing how the selected AMPs vary in sequence-specific characteristics thought to be important for their antibacterial activities. Some data are difficult to see because they overlap; see Figure S1A for a complete set of numerical values. **(C)** Cell schematic of the *E. coli* strain (CJW7845) used to examine the permeabilization of the inner and outer membranes to the indicated fluorescent proteins in response to AMP exposure. **(D)** Plot showing SYTOX Green uptake over time since the addition of different AMPs at concentrations equivalent to 2X MIC. Control antibiotics Rif and Spect were present at a final concentration of 50 μg/mL (∼3X MIC) and 128 μg/mL (2X MIC), respectively. The left plot shows the full range of measured SYTOX Green values. The right plot shows a zoom-in to visualize the results for the AMPs, or antibiotics, for which SYTOX Green uptake was lower. The total number of cells was 101, 288, 187, 196, 187, 205, 159, 158, 119, 201, 179, 176, 215, 110, 121, 140 for CecA, Tac1, PG-1, Mag2, LL-37, CRAMP, Mltt, HBD-3, Indo, Tur1B, PR-39, Bac7, Tur1A, CamBac, Rif, and Spect, respectively. **(E)** Plot showing the cumulative distribution of inner membrane (IM) permeabilization times for different AMPs and control antibiotics as a function of time since AMP or antibiotic addition, using the same set of treated cells as in (D). The total number of IM-permeabilized cells examined was 99, 273, 163, 176, 160, 183, 147, 82, 103, 45, 98, 26, 51, 11, 4, and 6 for CecA, Tac1, PG-1, Mag2, LL-37, CRAMP, Mltt, HBD-3, Indo, Tur1B, PR-39, Bac7, Tur1A, CamBac, Rif, and Spect, respectively**. (F)** Scatterplots showing the time of growth arrest since AMP addition vs. that of IM permeabilization since AMP addition across the AMPs. The total number of cells for each AMP is the same as in (E). The dashed line indicates *y = x*. Opacity of each data point was set to 15% to aid visualization of the data density. By definition, AMPs were classified based on whether the growth arrest that they cause is predominantly associated with IM permeabilization to proteins (class I, red) or not (class II, blue).

AMPs are often functionally classified based on their membrane-permeabilizing activity. This activity is commonly assessed using membrane-impermeant dyes that bind to nucleic acids such as SYTOX Green (Roth et al., 1997). Unlike the selected AMPs, SYTOX Green is small enough to diffuse into the periplasmic space through OM porins. Therefore, its uptake by Gram-negative bacteria such as *E. coli* is generally assumed to examine the integrity of the IM (Roth et al., 1997). Our first aim was to determine whether the growth arrest caused by the evaluated AMPs is associated with the permeabilization of the OM, IM, or both. To monitor the integrity of the two membranes independently, we constructed an *E. coli* strain (CJW7845) that carries two free fluorescent proteins: the red fluorescent protein mScarlet-I in the periplasmic space (i.e., between the IM and OM) and the blue fluorescent protein mTagBFP2 in the cytoplasm (Figure 1C). The mScarlet-I signal was frequently enriched at one or both cell poles due to a larger periplasmic space at these locations (Figure S1E).

Time-lapse wide-field microscopy was initiated before the addition of an AMP to ensure that cells grew normally in its absence. Each AMP was then added at a concentration equivalent to 2X MIC. This concentration was chosen to investigate the primary effects of the peptide on the cells, while also ensuring robust growth inhibition across the entire cell population and enabling a standardized comparison across the different AMPs tested. SYTOX Green was also included given the widespread use of fluorogenic dyes in membrane permeabilization assays. To analyse the data, we developed a computational pipeline to extract quantitative information (cell area, intracellular fluorescence intensities, etc.) for each cell over time to assess the start of growth inhibition, the growth arrest (Figure S2A), and the permeabilization of each cell membrane (see Methods). This is illustrated in Video 1, which shows a representative example of a cell treated with CecA, where permeabilization of both the OM and IM is indicated by the loss of the periplasmic and cytoplasmic fluorescent proteins.

### The tested AMPs fall into two phenotypic classes based on the timing of growth arrest relative to IM permeabilization to proteins

To begin, we analysed SYTOX Green uptake in cells treated with the different AMPs. To our surprise, the intracellular SYTOX Green signal differed widely across the fourteen AMPs investigated (Figure 1D), making it difficult to select a consistent threshold for defining IM permeabilization. In contrast, monitoring the loss or change in signal from the fluorescent proteins mScarlet-I and mTagBFP2 gave a reliable and robust method to monitor the permeabilization of each membrane to proteins over time (see Methods and Video 1 for an example).

Under our experimental conditions, all tested AMPs displayed some IM-permeabilizing activity, albeit to varying degrees (Figure 1E, see Methods for details on IM permeabilization detection). Even the proline-rich peptides PR-39, Bac7, CamBac, and Tur1A, which are typically regarded as nonmembrane-disruptive near MIC concentrations (Boman et al., 1993; Mardirossian et al., 2018; Podda et al., 2006; Scocchi et al., 2011), exhibited some degree of IM permeabilization (Figure 1E). Although their activity was considerably lower than that of other AMPs, all four proline-rich peptides showed higher permeabilization rates than the control antibiotics, rifampicin (Rif) and spectinomycin (Spect), which inhibit transcription and translation, respectively (Figure 1E).

The distinction among AMPs became clearer when we compared the timing of growth arrest and IM permeabilization to the fluorescent protein markers (Figures 1F and S2B). This resulted in two major classes. For class I (LL-37, CRAMP, CecA, Mag2, Tac1, PG-1, Mltt, HBD-3, Indo, and Tur1B), IM permeabilization to proteins was *predominantly* (i.e., in most cells) associated with the growth arrest (Figure 1F). For some of these peptides, we observed considerable cell-to-cell variability in the timings of IM permeabilization and growth arrest, occurring earlier in some cells and later in others (Figure 1F). However, regardless of this variability, both events remained closely associated in time. This is indicated by the data points falling close to the *y* = *x* diagonal in the plots (dashed line, Figure 1F), resulting in near-zero residuals from the diagonal (Figure S2B). The observed variability in event timings across cells highlights the importance of using a time-resolved method with single-cell resolution. For class II AMPs (PR-39, Bac7, CamBac, and Tur1A), when IM permeabilization did occur, it happened after the growth arrest (Figure 1F), resulting in high residuals from the diagonal (Figure S2B). This observation indicates that IM permeabilization to proteins was not the underlying cause of growth inhibition for these class II peptides. Furthermore, unlike the abrupt growth arrest characteristic of class I AMPs, class II AMPs induced a more gradual growth inhibition, with a substantial delay between the initiation of growth slowing and complete arrest (Figure S2C).

### AMPs can be further categorized based on their outer membrane effect

The class I phenotype could be further subdivided into two subtypes based on the relative timing of OM and IM permeabilization (Figure 2A): type A, in which OM permeabilization to fluorescent proteins predominantly preceded or closely followed IM permeabilization (< 10 min), and type B, in which the OM permeabilization either did not occur or did occur but considerably later (> 10 min) than IM permeabilization. The type A phenotype was prevalent for cells exposed to CecA, PG-1, Tac1, LL-37, CRAMP, or Mag2. It was characterized by the high rate of OM permeabilization among cells that became IM-permeabilized (Figure 2B), along with the strong temporal concurrence between these two permeabilization events (Figure 2C). This is illustrated in Video 1 and its corresponding fluorescence kymographs (Figure 2D) showing how a representative CecA-treated cell abruptly stopped growing and shrunk in size at approximately the same time as the periplasmic mScarlet-I signal disappeared, marking the OM permeabilization event. Almost immediately after, the cytoplasmic mTagBFP2 signal vanished due to IM permeabilization.

**Figure 2:**
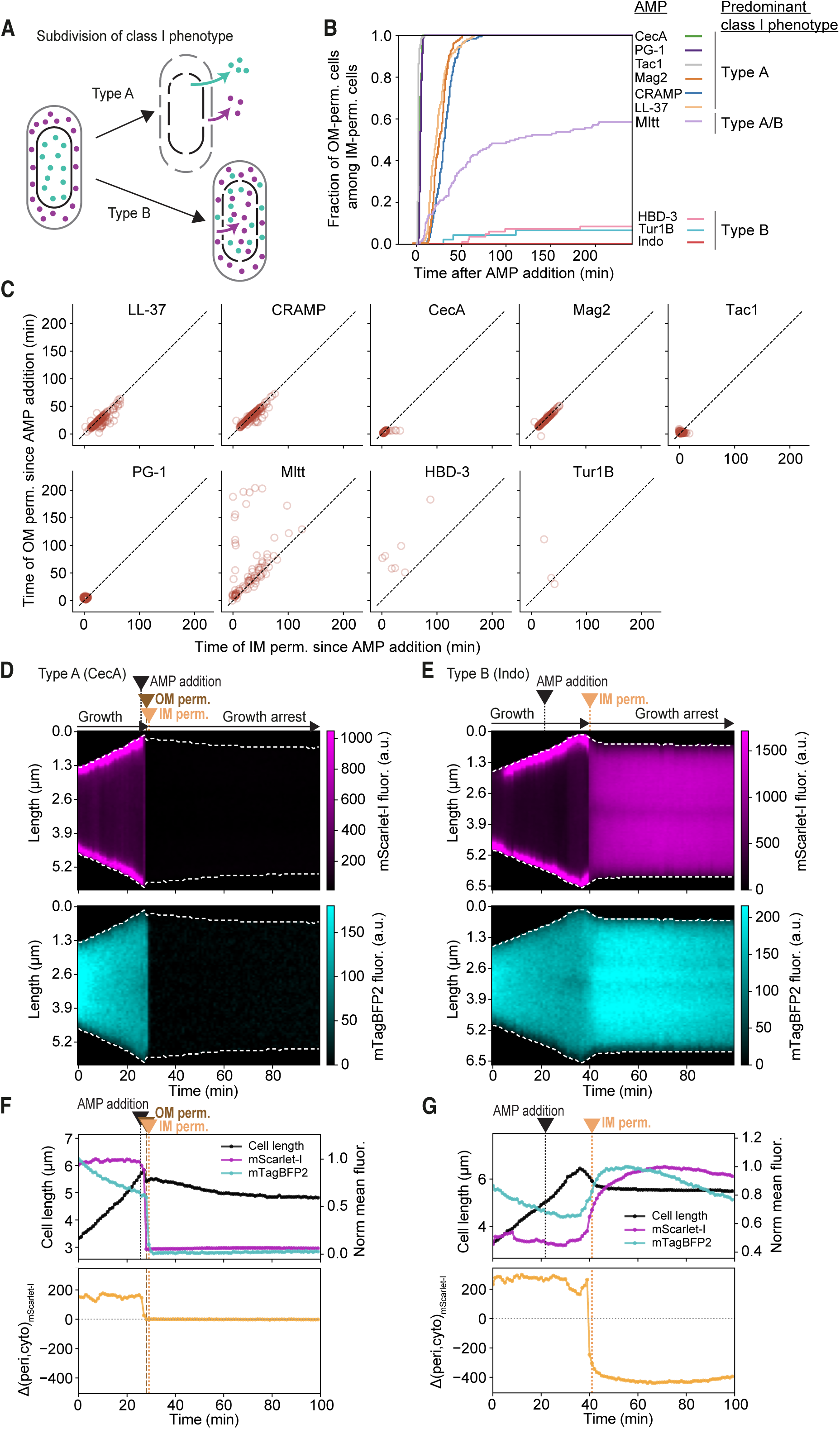
Class I AMPs can be subdivided into types A and B based on the relative timing of permeabilization of each membrane. **(A)** Schematics showing what happens to the cytoplasmic mTagBFP2 and periplasmic mScarlet-I for each class I AMP type. By definition, type A predominantly permeabilizes the outer membrane (OM) of cells simultaneously with, or before, the inner membrane (IM), resulting in the release of both mScarlet-I and mTagBFP2 into the environment. In contrast, type B primarily permeabilizes the IM well before the OM (which may or may not permeabilize during the experiment, see panel B), resulting in the mixing of mScarlet-I and mTagBFP2 within the OM boundaries. **(B)** Plot showing the cumulative distribution of OM permeabilization events for different class I AMPs used at concentrations equivalent to their 2X MICs (Figure 1A). Here, only cells for which the IM became permeabilized during the experiment were considered. They represent 99/99, 273/273, 163/163, 176/176, 160/160, 183/183, 86/147, 7/82, 3/45, and 0/103 cells for CecA, Tac1, PG-1, Mag2, LL-37, CRAMP, Mltt, HBD-3, Tur1B, and Indo, respectively. **(C)** Scatterplots showing the time of OM permeabilization since AMP addition as a function of IM permeabilization since AMP addition across the AMPs. Opacity of each data point was set to 25% to aid visualization of the data density. The dashed line indicates *y = x*. **(D**) Example kymographs of mScarlet-I and mTagBFP2 signals for a representative CJW7845 cell exposed with CecA (2X MIC) at the indicated time. The times of IM and OM permeabilizations are shown. See also Video 1. **(E)** Same as (D) but for a cell treated with Indo (2X MIC). See also Video 2. **(F)** Plots showing the quantification of the cell length and fluorescence signal (top) and the difference Δ(peri,cyto)_mScarlet-I_ between the mScarlet-I signal in the cell periphery (periplasm) relative to the cell interior (cytoplasm) over time for the cell shown in panel (D). **(G)** Same as (F) except for the cell shown in (E).

For type B, the sequence of events was less intuitive, as the OM became permeable to proteins well after the IM or not at all (Figure 2A). Video 2 and its corresponding kymographs (Figure 2E) showcase an example of this type B phenotype for a representative Indo-treated cell. Permeabilization of the IM but not OM was evident by the periplasmic mScarlet-I signal moving into the cytoplasm and mixing with mTagBFP2. This event was readily detectable, in contrast to the corresponding SYTOX Green signal, which increased only slowly (Video 2, Figure 1D), in stark contrast to the rapid dye uptake when the OM is disrupted first by a class I type A AMP (Video 1, Figure 1D). This indicates that the OM presents a significant barrier even for small dyes and that the slow dye uptake characteristic of type B AMP (Figure S2D) would have been difficult to interpret without the orthogonal readout of OM and IM integrity provided by our dual-reporter strain.

As previously demonstrated for growth-arrested cells with protein synthesis inhibitors (Balleza et al., 2018; Bevis & Glick, 2002; Hebisch et al., 2013; Megerle et al., 2008), the signal intensity of the fluorescent reporters in cells treated by a class I type B AMP increased after growth arrest due to the delayed fluorophore maturation, before subsequently decreasing as a result of photobleaching (Video 2). The transient increase in fluorescent protein signal was exacerbated by the cell shrinkage. For Indo and Tur1B, OM permeabilization was either absent or rare during the experiment (Figure 2B). For HBD-3, OM permeabilization in IM-permeabilized cells occurred at a low rate (Figure 2B), and when it happened, it often was after the IM permeabilization event (Figures 2C and S3A).

Quantitatively, treatment with class I type A peptides resulted in an abrupt drop in mScarlet-I and mTagBFP2 signals inside cells due to the permeabilization of both membranes (Figure 2F). Class I type B also caused IM permeabilization, but the cells maintained both fluorescent protein signals during all or part of the experiment (Figures 2E and S3A). To track this specific phenotype, the timing of IM permeabilization was determined by calculating the difference in the mScarlet-I signal between the cell periphery (periplasm) and the cell interior (cytoplasm). This metric Δ(peri, cyto)_mScarlet-I_ changed rapidly from a positive value to a negative one once mScarlet-I proteins entered the cytoplasm due to IM permeabilization (Figures 2G and S3A). Thus, our analysis shows that even though class I type B AMPs predominantly permeabilize the IM of *E. coli*, they do not release protein content into the environment at the time of growth arrest.

For class II peptides, the occasional late IM permeabilization events observed with class II PR-39, Bac7, Tur1A, and CamBac (Figure 1F) did not result in protein leakage: neither mScarlet-I nor mTagBFP2 signal was lost. Instead, IM permeabilization was apparent by the mixing of the two signals (Figure S3B-D), indicating that periplasmic and cytoplasmic antigens remained enclosed within the cell by the intact OM barrier. Thus, all 4 class II peptides tested in this study behave as type B with respect to membrane permeabilization.

### Growth inhibition by the class I AMPs is also associated with intracellular rearrangement of ribosomes and DNA

Next, we examined intracellular effects at the time of growth inhibition. We focused on the chromosome (which folds into a structure known as the nucleoid) and the ribosomes because these two large anionic cytoplasmic components are often proposed as intracellular targets of AMPs (Cardoso et al., 2019; Le et al., 2017), and the AMP LL-37 has been shown to perturb their intracellular organization (Zhu et al., 2019). During unperturbed growth, most ribosomes are bound to mRNAs and are engaged in translation, forming monosomes and polysomes (collectively referred to as polysomes for simplicity). Because of their large size relative to the DNA mesh (Xiang et al., 2021), polysomes are largely excluded from the nucleoid, resulting in ribosomal accumulation in DNA-depleted regions (Bakshi et al., 2012; Gray et al., 2019; P. J. Lewis et al., 2000; Linnik et al., 2024; Sanamrad et al., 2014; Xiang et al., 2021). This segregation pattern can be visualized by microscopy using fluorescent protein fusions to ribosomal and DNA-binding proteins such as RplA-GFP and HupA-mCherry, respectively (Papagiannakis et al., 2025; Xiang et al., 2021). Their anticorrelation in localization results in negative signal correlation factor (SCF) values (Gray et al., 2019).

To visualize several intracellular components at the same time, we used a strain (CJW7753) that expresses three different cytoplasmic markers (Figure 3A): RplA-GFP, HupA-mCherry, and mTagBFP2. Time-lapse imaging under the same standardized conditions as above demonstrated that the intracellular distribution of ribosomes and DNA changed abruptly when cells were exposed to class I type A AMPs. This is illustrated with CecA where the ribosome (RplA-GFP) and DNA (HupA-mCherry) signals became transiently mixed near the time of growth arrest (Video 3 and Figure 3B). This sudden mixing was followed within a few minutes by re-separation (unmixing) of the ribosome and DNA signals (Figure 3B), as quantified by a transient peak in positive SCF values in individual cell trajectories (Video 3). When all cell trajectories were aligned by the SCF peak, we observed a clear temporal pattern across CecA-treated cells (Figure 3C). The SCF peak (i.e., peak of ribosome/DNA mixing) occurred near the time of growth arrest, characterized by the abrupt cell shrinkage (Figure 3C). Re-segregation of the ribosome and DNA signals occurred concomitantly with DNA compaction, which was quantified by a reduction of the nucleocytoplasmic (NC) ratio (DNA signal length divided by the cell length) (Figure 3C). The mixing of ribosome and DNA signals indicates a reduction or disappearance of polysomes, which, unlike free ribosomes and ribosomal subunits, are too large to freely diffuse through the nucleoid (P. J. Lewis et al., 2000; Bakshi et al., 2012; Sanamrad et al., 2014; Gray et al., 2019; Xiang et al., 2021; Linnik et al., 2024). Since polysomes represent actively translating ribosomes, their loss is indicative of translation inhibition. Interestingly, the homogenization of the ribosome signal often slightly preceded the loss of cytoplasmic mTagBFP2 signal (Figure 3B), suggesting that intracellular ribosome/DNA reorganization can occur before and thus independently of membrane damage that leads to protein leakage. However, since the two events were typically separated by just a minute (our time resolution), we cannot rule out the presence of more subtle membrane damage at the time of ribosome/DNA mixing.

**Figure 3.**
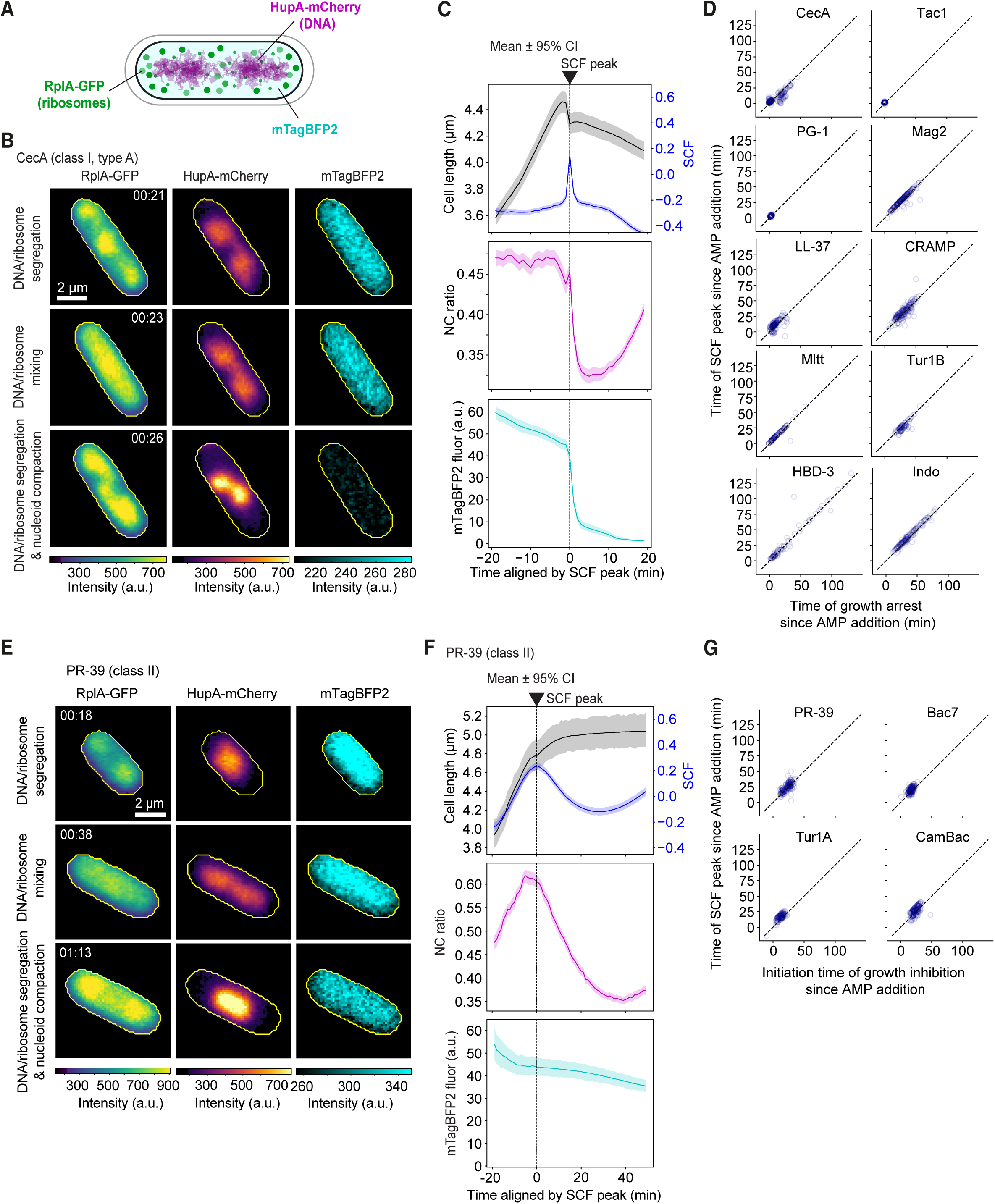
Both class I and class II AMPs feature similar intracellular reorganization of ribosomes and DNA at the time of growth inhibition. **(A)** Cell schematic of strain CJW7753 expressing mTagBFP2, RplA-GFP, and HupA-mCherry as markers for cytoplasmic proteins, ribosomes, and DNA, respectively. **(B)** Images of a representative CJW7753 cell at the indicated specific timepoints (h:min) illustrating changes in intracellular ribosome and DNA distributions. CecA (2X MIC) was added at 19 min. See Video 3 for the whole time-lapse sequence. **(C)** Plots showing the quantification of the evolution of the mean ± 95% confidence interval (CI) of the cell length, signal correlation factor (SCF), nucleocytoplasmic (NC) ratio, and mTagBFP2 fluorescence (fluor.) across 314 cells. Times were aligned based on the SCF peak in each cell (indicated by the black dashed line). **(D)** Scatterplots showing the time of SCF peak as a function of the time of growth arrest across the indicated class I AMPs (2X MIC). The total number of cells is 314, 117, 113, 205, 172, 189, 102, 78, 84, and 152 for CecA, Tac1, PG-1, Mag2, LL-37, CRAMP, Mltt, HBD-3, Tur1B, and Indo, respectively. Opacity of each data point was set to 15% to aid visualization of the data density. The dashed line indicates *y = x*. **(E)** Images of a representative CJW7753 cell at the indicated specific timepoints (h:min) illustrating changes in intracellular ribosome and DNA distributions. PR-39 (2X MIC) was added at 18 min. See Video 4 for the whole time-lapse sequence. **(F)** Plots showing the quantification of the evolution of the mean ± 95% confidence interval (CI) of the cell length, smoothened signal correlation factor (SCF), nucleocytoplasmic (NC) ratio, and mTagBFP2 fluorescence (fluor.) across 180 cells. Times were aligned based on the smoothened SCF peak at the single-cell level. The black dashed line indicates the smoothened SCF peak (time zero). **(G)** Scatterplots showing the time of SCF peak as a function of the start of growth inhibition for the indicated class II AMPs. The total number of cells is 180, 183, 131, and 168 for PR-39, Bac7, Tur1A, and CamBac (2X MIC), respectively. Opacity of each data point was set to 15% to aid visualization of the data density. The dashed line indicates *y = x*.

We also observed an abrupt and transient mixing of ribosome and DNA signals (positive SCF peak) occurring near the time of growth arrest for the nine other class I AMPs, as indicated by the data points lying close to the *y* = *x* diagonal (Figure 3D). At later timepoints, the behavior of the ribosome and DNA signal was more variable. We did not carefully characterize these late events, as the focus was on the events that occur near the time of growth inhibition.

### Similar intracellular rearrangements occur with class II AMPs but with slower kinetics

Remarkably, despite their difference in membrane activity, class II AMPs PR-39, Bac7, Tur1A, and CamBac induced a mixing between ribosome (RplA-GFP) and DNA (HupA-mCherry) signals similar to class I peptides, resulting in positive SCF values (Figure 3E and Video 4 for a PR-39-treated cell example). This mixing was followed by both ribosome/DNA re-segregation (decreasing SCF values) and DNA compaction (decreasing NC ratio). However, the kinetics of these events were considerably more gradual for these class II peptides, as shown by the broader average SCF peaks and slower decays in NC ratio (Figures 3F and S4) compared to class I peptides (e.g., Figure 3C). For all four class II AMPs, the timing of the ribosome/DNA mixing (SCF peak) coincided with the onset of growth inhibition (Figure 3G). Thus, all fourteen AMPs tested caused the same DNA/ribosome mixing event near the time of growth inhibition, but with different kinetics depending on their designated class.

### The same phenotyping classification can apply to antimicrobial peptidomimetics

Given the prospect of AMPs as antimicrobial therapeutics, significant efforts have been invested in developing synthetic AMP mimics that retain the potency of natural peptides while addressing their limitations (e.g., bioavailability, toxicity, and production costs) (Svenson et al., 2022). These peptidomimetics (PMs) are often smaller and more resistant to protease cleavage due to the incorporation of non-natural chemical modifications. To examine whether our phenotype-based classification framework applies to peptidomimetics, we selected four for testing against the strain (CJW7845) that carries the periplasmic mScarlet-I and the cytoplasmic mTagBFP2.

The first three were peptoids TM1, TM4, and TM5 (Figure 1A), which have broad-spectrum antibacterial activity (Chongsiriwatana et al., 2008, 2017). Peptoids resist protease activity because they differ from peptides by having the side chains attached to the backbone amide nitrogen instead of the α-carbon (Zuckermann et al., 1992). The fourth synthetic AMP mimic was the D-enantiomeric peptide DJK-5 (Figure 1A), known for its efficacy against biofilms (De La Fuente-Núñez et al., 2014, 2015).

We found that at 2X MIC (Figures 1A and S1B-C), all three peptoids behaved as class I AMPs, as the permeabilization of the IM was fast (Figure S5A) and coordinated with an abrupt growth arrest (Figures S2B-C and S5B). The permeabilization of the OM was comparatively less frequent (Figure S5C) and typically delayed (Figure S5D), consistent with a class I type B assignment. IM permeabilization was apparent by the internalization of periplasmic mScarlet-I, as illustrated in single-cell kymographs (Figure S5E). In contrast, at 2X MIC, DJK-5 predominantly behaved as a class II peptide. IM permeabilization occurred at a slower rate than the peptoids (Figure S5A), and in most cells, this event happened after the growth arrest (Figures S2B and S5B), which happened less abruptly as compared to the peptoids (Figure S2C). For the DJK-5-treated cells in which IM permeabilization occurred, a fraction of them also displayed OM permeabilization (Figure S5C), but usually after IM permeabilization (Figure S5D and F), indicating a class II type B phenotype. Together, these results demonstrate the applicability of our phenotyping framework for characterizing both AMPs and PMs.

### Increasing peptide concentration to 10X MIC accelerates antibacterial activity but does not alter classification

Some AMPs like Bac7, Tur1A, and Arasin-1 have been reported to be membrane-inactive at MIC concentrations, but to gain membrane-permeabilizing activity at higher concentrations (Podda et al., 2006; Mardirossian et al., 2018; Paulsen et al., 2013). To examine whether our phenotypic classification is highly sensitive to the AMP/PM concentration, we exposed cells (CJW7845) to concentrations equivalent to 10X MIC. Two compounds were excluded: HBD-3, for which the combination of a high MIC and high cost made experiments at this concentration prohibitively expensive, and TM4, for which aggregate formation and severe cell deformation observed at this concentration precluded quantitative image analysis. For the remaining 16 AMPs/PMs, we found that the higher concentration typically shortened the time to growth arrest and reduced cell-to-cell variability (Figure S6A). Conversely, a lower concentration (1X MIC), which was only tested on a subset of AMPs, often had the opposite effects (Figure S6A).

The acceleration in antibacterial activity at 10X MIC was most pronounced for class I AMPs/PMs, which were able to permeabilize the IM and arrest cell growth within minutes of exposure (Figure S6A-E). This aligns with the model in which membrane-disruptive AMPs must overcome a critical surface coverage on the bacterial membrane to trigger its disruption (Melo et al., 2009). The effect was more modest for class II peptides; even at 10X MIC concentrations, they still required tens of minutes to fully inhibit growth (Figure S6A) and when inhibition was initiated, reaching growth arrest was still gradual unlike for most class I AMPs, especially for the tested natural class II (Figure S6E). This is consistent with the proposed reliance of class II peptides such as PR-39, Bac7, Tur1A, and CamBac on specific cellular transporters (Mattiuzzo et al., 2007; Krizsan et al., 2015), an uptake mechanism that is inherently saturable.

Importantly, these peptides retained their class II affiliation, as growth arrest still preceded IM permeabilization in most cells (Figure S6C-D) despite the increased IM-disrupting activity (Figure S6B). The A and B subtype classification was also relatively robust to higher concentrations for both class I and class II AMPs/PMs, except perhaps for TM5 for which the increase in OM permeabilizing activity rendered the type A/B subtyping more ambiguous (Figure S6E-F).

### Intracellular absorption drastically differs in kinetics between class I and class II peptides

LL-37 tagged with a fluorophore has been shown to accumulate at high levels inside cells (Snoussi et al., 2018; Zhu et al., 2019). Since LL-37 is a class I AMP, membrane permeabilization may contribute to this rapid peptide uptake. If so, then there could be a considerable difference in the rate of peptide accumulation between class I and class II peptides. To test this possibility, we commissioned the synthesis of fluorescent 5-TAMRA conjugates to AMPs with high membrane-permeabilizing activity: class I type A peptides CecA, Tac1, and LL-37 (Figure 1E-F). For comparison with peptides with lower membrane-permeabilizing activity, we also obtained 5-TAMRA conjugates to class II PR-39, Bac7, and DJK-5. We found that these fluorescent AMP derivatives retained antibacterial activities, featuring slightly higher or lower MIC values than the unconjugated peptides (Figure S1F-G). To minimize potential secondary effects associated with the fluorophore tag, we treated cells with unlabeled AMPs at 2X MIC, spiked with a low amount of fluorescently labeled derivatives (50 nM final concentration).

For all three class I CecA, Tac1, and LL-37 conjugates, we observed rapid uptake and massive intracellular accumulation of fluorescence signal within just 10 min of growth inhibition (Figure 4A). This stood in striking contrast to the class II derivatives PR-39, Bac7, and DJK-5 (Figure 4A). At this 10-min timepoint, the intracellular signal from the class I conjugates was ∼400-1000 times higher than for the class II conjugates (Figure S7A), consistent with the fast membrane permeabilization by class I peptides promoting their rapid self-promoted uptake.

**Figure 4.**
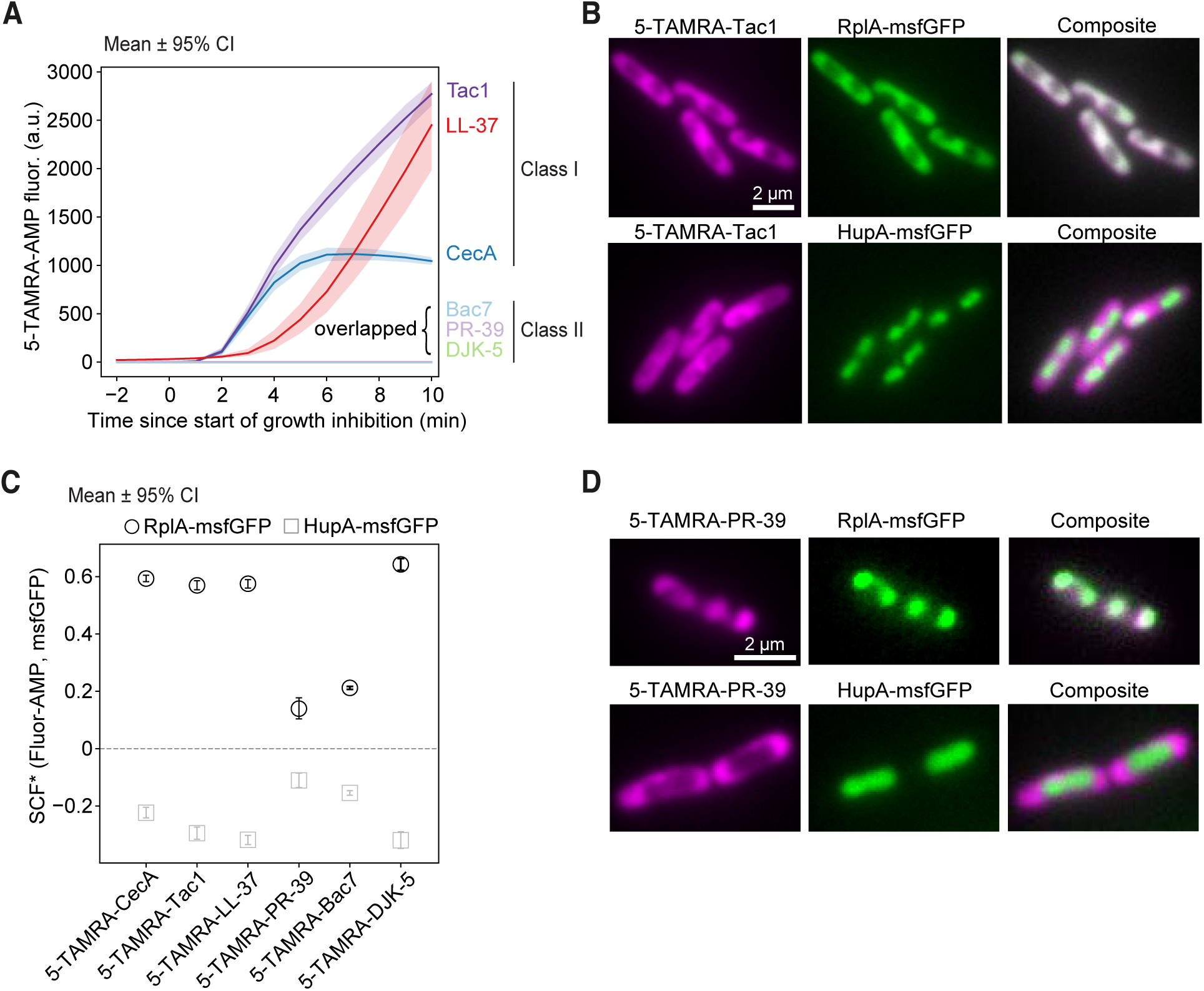
The intracellular distribution of fluorescent AMP derivatives correlates with the ribosome signal and anti-correlates with the DNA signal. **(A)** Evolution of the intracellular fluorescence intensity (mean ± 95 % CI) of 5-TAMRA-AMP conjugates, present at 50 nM together with the corresponding unlabeled AMPs (2X MIC) in MG1655 cells. Time axis of each single cell was aligned by start time of growth inhibition. The total number of averaged cells is 138, 116, 101, 125, 254, and 180 for CecA, Tac1, LL-37, PR-39, Bac7, and DJK-5, respectively. (B) *Top*, representative images of the indicated fluorescence signal in CJW7020 cells (expressing RplA-msfGFP) exposed to a mixture of Tac1 (2X MIC) and 5-TAMRA-Tac1 (60 nM, 5% of the unlabeled version) for 50 min. *Bottom*, representative images of the indicated fluorescence signal in CJW7859 cells (expressing HupA-msfGFP) exposed to a mixture of Tac1 (2X MIC) and 5-TAMRA-Tac1 (60 nM, 5% of the unlabeled version) for 20 min. See Methods for explanation about the timepoint selection. **(C)** Plot showing mean ± 95 % CI of the signal correlation factor (SCF*, which is slightly modified version of SCF, see Methods) between the fluorescently labeled AMP signal and the RplA-msfGFP (black circles) or HupA-msfGFP (grey squares) fluorescent signal. CJW7020 and CJW7859 cells (expressing RplA-msfGFP or HupA-msfGFP, respectively) were treated with a mixture of CecA (2X MIC), Tac1 (2X MIC), LL-37 (2X MIC), PR-39 (1.5X MIC), Bac7 (2X MIC), and DJK-5 (2X MIC), plus 5% (or 33% for PR-39, 100% for Bac7) of the corresponding fluorescently labeled version, which resulted in the final concentrations of 50 nM 5-TAMRA-CecA, 60 nM 5-TAMRA-Tac1, 100 nM 5-TAMRA-LL-37, 1 µM 5-TAMRA-PR-39, 1.2 µM 5-TAMRA-Bac7, and 250 nM 5-TAMRA-DJK-5, respectively. See Methods for the timepoint selection. The total number of cells for CJW7020 was 1358 for 5-TAMRA-CecA, 726 for 5-TAMRA-Tac1, 623 for 5-TAMRA-LL-37, 135 for 5-TAMRA-PR-39, 2512 for 5-TAMRA-Bac7, and 369 for 5-TAMRA-DJK-5. For CJW7859 the number of cells was 566 for 5-TAMRA-CecA, 375 for 5-TAMRA-Tac1, 480 for 5-TAMRA-LL-37, 191 for 5-TAMRA-PR-39, 1001 for 5-TAMRA-Bac7, and 176 for 5-TAMRA-DJK-5. **(D)** Same as (B) except that the selected CJW7020 cell expressing RplA-msfGFP (top) CJW7859 cell expressing HupA-msfGFP (bottom) treated with a 75:25 mixture of PR-39 (1.5X MIC) and 5-TAMRA-PR-39 (1 µM, 33% of the unlabeled version) were permeabilized, enhancing the intracellular uptake and absorption of PR-39 and its fluorescent derivative. See Methods for the timepoint selection.

The accumulation of class I AMPs inside cells implies that they bind to intracellular targets. Interestingly, when we examined the fluorescence of the AMP conjugates at late time points of treatment—when the ribosome (RplA-GFP) and the DNA (HupA-mCherry) signals are well separated through mutual exclusion—it was not homogeneously distributed inside cells. Instead, 5-TAMRA-tagged Tac1, CecA, and LL-37 (class I) were found to co-localize with the ribosome signal in RplA-msfGFP-expressing cells and to anti-correlate with the DNA signal in HupA-msfGFP-expressing cells (Figures 4B and S7B-C). At the population level, the correlations between each AMP signal and the ribosome marker (RplA-msfGFP), as well as the anti-correlation with the DNA marker (HupA-msfGFP), were quantitatively confirmed by the corresponding positive and negative correlation values, respectively (Figure 4C).

Due to their limited intracellular uptake, class II peptides showed weaker correlations with these cellular components. They did, however, colocalize with ribosomes and avoided the DNA (Figure 4C), albeit with a faint signal that required spiking in a higher concentration of the fluorescent derivative to detect (25:75 labeled-to-unlabeled ratio for PR-39 and at 50:50 for Bac7). Notably, a minority of PR-39-treated cells displayed a strong intracellular signal, likely due to a late membrane permeabilization event enabling a large peptide influx, similar to class I AMPs. In these specific cells, the peptide clearly colocalized with ribosomes (RplA-msfGFP) and was excluded from the DNA (HupA-msfGFP), mirroring the behavior of class I peptides (Figure 4D).

Thus, all tested fluorescent AMPs are excluded from the DNA and colocalize with ribosomes. This strongly suggests that ribosomes are a major intracellular target and that their high abundance creates a cellular ‘sink’, driving the massive peptide accumulation seen after membrane permeabilization.

### Class II antimicrobials are more effective than class I against dense populations

Membrane-permeabilizing activity is widely regarded as a desirable characteristic in an AMP or PM. This step not only amplifies the peptide’s own influx via self-promoted uptake but also provides access to periplasmic and intracellular targets (Hancock & Sahl, 2006), extending the range of its antimicrobial action. Furthermore, the high membrane-disruptive activity of class I AMPs results in faster antibacterial activity kinetics compared to class II AMPs (Figures 3B-C vs E-F and S2C), particularly at high concentrations (Figure S6). However, the uptake kinetics of fluorescent derivatives suggest that membrane permeabilization also promotes high peptide absorption inside cells (Figures 4A and S7A). High intracellular accumulation of LL-37 — a class I peptide — has been shown to cause a phenomenon known as the inoculum effect (Snoussi et al., 2018) where higher concentrations of an antibacterial agent are needed to kill higher-density populations.

The contrasting kinetics between class I and II AMPs inspired us to think about potential trade-offs and how they may be relevant in physiological contexts. In our experiments described above, *E. coli* cells were present at a low density, and the AMPs were delivered homogeneously. Inside a host, the delivery of AMPs can be polarized, for example, when they are secreted by the mammalian intestinal epithelium into the mucus in the gut lumen (Figure 5A). In vivo, immune cells such as neutrophils may encounter dense cell populations (Figure 5A), for example, when the infection has reached a high level of bacterial load or when the pathogen forms biofilms. To mimic both effects (polarized delivery and high cell density), we built a microfluidic device that was adapted from an earlier design (Lambert & Kussell, 2014). This device contains wide trenches where bacterial cells can grow at a high cell density (Figure 5B). A constant flow of fresh nutrients rapidly diffuses from the “feeding” channel into the trenches. Excess cells at the top of the trenches are continuously removed by the flow. AMPs are supplemented in the growth medium, creating a constant concentration in the feeding channel. Due to diffusion, their delivery in the cell-filled trenches is polarized, with cells at the top of the trenches encountering the peptides first. In this context, we predicted that class I peptides would be rapidly sequestered by the first cells they permeabilize. This sequestration or “soaking” effect would limit diffusion of the peptides through the trenches, making them less effective than class II AMPs at controlling growth across dense populations.

**Figure 5.**
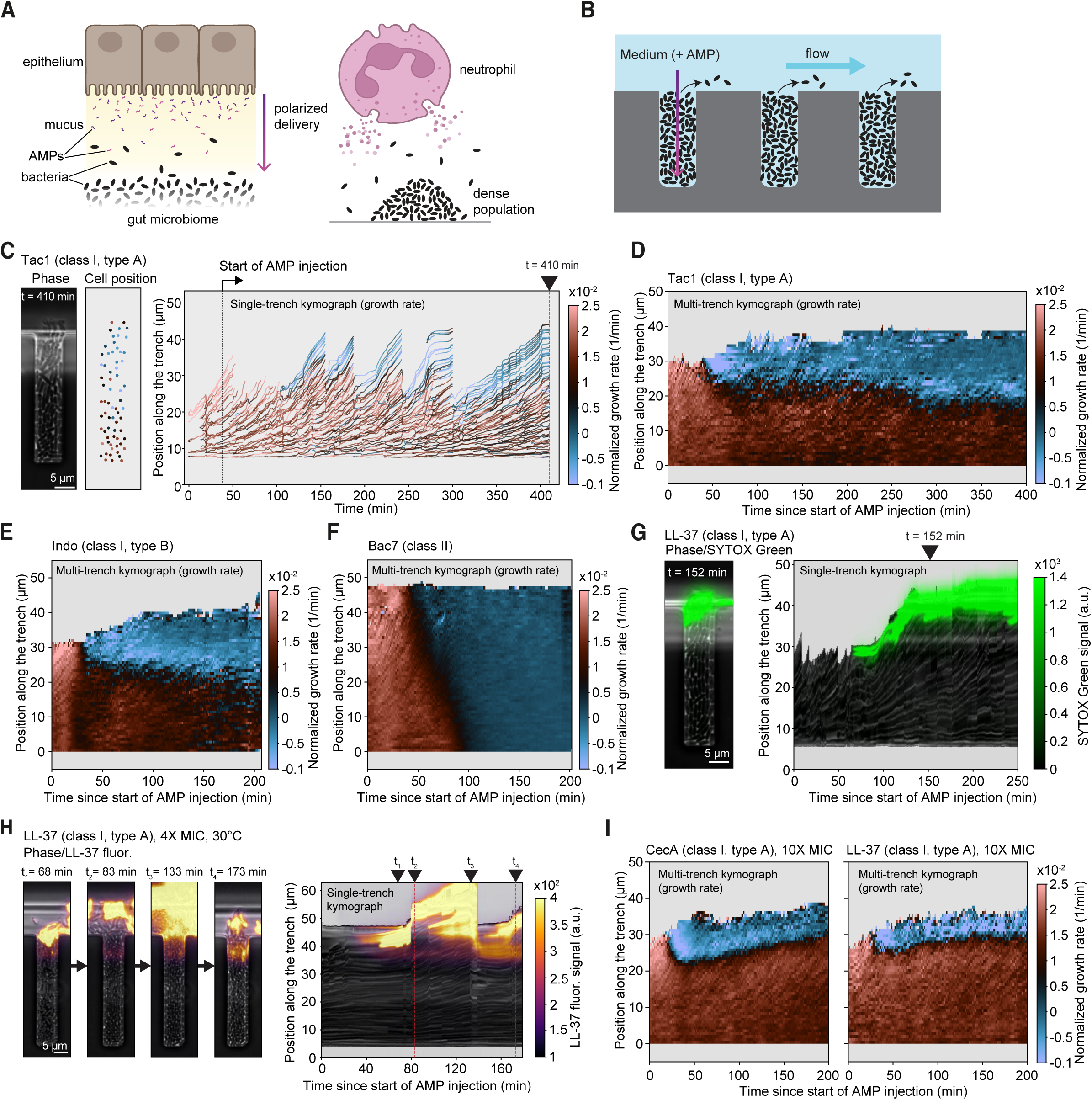
Class I AMPs are less effective against high cell densities than class II peptides. **(A)** Schematics illustrating the AMP release by a mammalian gut epithelium (left) and a neutrophil (right). Adapted from a BioRender template (BioRender.com). **(B)** Schematic of the wide-trench microfluidic device. Purple arrow indicates the polarized delivery of AMPs to emulate the in vivo scenarios in (A). **(C)** Left: Representative phase contrast image of CJW7753 cells in a wide trench at t = 410 min from the start of the acquisition and corresponding detected cell positions when Tac1 (2X MIC) is present in the medium flowed in the feeding channel. See Video 5 for the whole time-lapse sequence showing the corresponding single-cell tracking. Right: Kymograph showing cell vertical positions along the corresponding trench over time. Color coding represents single-cell normalized growth izedrate. **(D)** Ensemble kymograph computed across 23 trenches for CJW7753 cells treated with Tac1 (2X MIC), showing the spatial distribution of growth rate across trenches. Color coding represents the average normalized growth rate across trenches at a given time and vertical position. The number of cells is 7104. **(E-F)** Same as (D) but for cells exposed to Indo (E) or Bac7 (F) at 2X MIC. The number of trenches is 31 (Indo) and 27 (Bac7); the number of cells is 6195 and 4648, respectively. **(G)** Left: Representative phase-contrast image of CJW7845 cells overlaid with SYTOX Green fluorescence signal at t =152 min after the start of AMP injection. Right: Kymograph of the same single trench showing the mean projection along the trench vertical axis of the phase contrast (representing all the cells in the trench) and overlaid SYTOX Green signal (representing permeabilized cells). Videos 7 and 8 show the timelapses of CJW7845 cells treated with LL-37 and CecA, respectively, with addition of SYTOX Green (5 nM) in the flowing medium, for multiple trenches. Figure S8A and S8B show the ensemble kymographs computed from averaging 24 trenches exposed to LL-37 or CecA, respectively. **(H)** Same as (G) but for CJW7859 strain treated with 4X MIC LL-37 at 30°C, displaying the vertical distribution of LL-37 fluorescence along a single trench over time. The four selected timepoints illustrate how LL-37-absorbing cells at the top of the trench are pushed out by the growing cells below. Video 9 shows a whole time-lapse sequence for two representative cell-filled trenches. Figure S8C shows the ensemble kymograph computed from averaging across 18 trenches. **(I)** Same as (D-F) but for cells exposed to 10X MIC of CecA (left) or LL-37 (right). The number of trenches is 30 (CecA), and 16 (LL-37); the number of cells is 7730 and 3616, respectively.

Our findings support this prediction. At the 2X MIC concentration, which was effective to control low-density populations, class I AMPs primarily inhibited the growth of cells near the top of the trenches whereas cells below continued to grow unimpeded (as illustrated in Video 5 and Figure 5C for Tac1). This is consistent with the “frontline” cells rapidly sequestering the incoming peptides. This dynamic created a cyclical process, where growth from the protected lower cells (in brown) continuously pushed the arrested cells (in blue) out of the trench. This localized growth inhibition was highly reproducible across trenches and across representative class I AMPs (LL-37, CecA, CRAMP, Mag2, Mltt) and PM TM1, as shown in ensemble kymographs (Figures 5D and S8A). These results are consistent with the heterogeneous killing behavior of LL-37 (Snoussi et al., 2018). Only class I type B Indo performed slightly better by occasionally inhibiting the growth of cells located further into the trenches (Figure 5E). This is presumably because this peptide does not permeabilize the OM (Figure 2B), slowing its uptake. However, even Indo failed to stop the growth of most cells in the trenches (Figure 5E).

In striking contrast, at 2X MIC, all four class II AMPs (PR-39, Bac7, CamBac, and Tur1A) and PM DJK-5 were able to inhibit cell growth throughout the entire wide trenches, even in trenches that are slightly deeper (Figures 5F, S8B, and Video 6). We concluded that the slow uptake of class II peptides allows them to diffuse throughout the trench, resulting in growth inhibition of the entire cell population.

We also confirmed that class I AMPs, such as LL-37 and CecA (2X MIC), only permeabilized frontline cells by including SYTOX Green in the medium. Indeed, only the top-layer cells incorporated the dye (Figures 5G and Videos 7-8), which was highly reproducible across multiple trenches (Figure S8C-D). To verify that rapid intracellular absorption of class I AMPs is what prevents them from diffusing past the first cells they permeabilize, we exploited our accidental discovery that LL-37 exhibits low intrinsic fluorescence (predominantly in the red channel) at 30°C. This property allowed us to visualize peptide diffusion without adding a potentially interfering fluorophore. We found that the fluorescence of LL-37 (4X MIC) rapidly spread across the entire length of empty trenches (Figure S8F-G and Video 9), demonstrating that diffusion itself was not inherently limited. In stark contrast, LL-37 was unable to diffuse across nearby trenches filled with cells. Instead, the peptide’s diffusion was arrested at the frontline where it rapidly accumulated inside cells, forming a bright fluorescent ‘crust’ (Figures 5H, S8E, and Video 9). This crust occasionally compressed the growing cells below, which ultimately pushed the protective crust out of the trenches.

Increasing the concentration of class I peptides to 10 times the concentration needed for neutralizing low-cell-density populations was insufficient to overcome the intracellular “soaking” effect, as shown with LL-37 and CecA at 10X MIC (Figure 5I). The result was similar: only the top-layer cells were growth-inhibited, with no impact on the growth of the underlying cells.

Altogether, our results reveal opposing functional trade-offs between classes. High membrane-permeabilizing activity and self-promoted uptake confer class I AMPs/PMs remarkable speed and multipronged action. However, these same properties also drive their rapid intracellular sequestration and extracellular depletion, making them ineffective against dense populations. Conversely, class II peptides, with weaker membrane activity and slower uptake, act more slowly yet remain effective at higher cell densities.

### Class II AMPs are more effective than class I against biofilms

Bacteria frequently form biofilms in nature and during infections, and high cell density is a defining feature of biofilms. We therefore asked whether the functional trade-off between class I and class II peptides extends to this physiological context. To test this idea, we turned to *E. coli* ATCC 25922, a clinical isolate known for its propensity to form biofilms (Naves et al., 2008). We confirmed that a GFP-expressing derivative (ATCC 25922GFP) rapidly formed biofilms under our experimental conditions (M9gluCAAT, 37°C), visible as dense GFP-labeled cell clusters and mushroom-like structures, with the extracellular matrix (ECM) labeled by the fluorogenic amyloid/polysaccharide-binding dye EbbaBiolight 680 (Video 10 and Figure 6A). To monitor sustained biofilm growth in static conditions starting from a pre-formed biofilm, we first inoculated a diluted culture (OD_600_ = 0.015) into an eight-well chamber slide and allowed growth and colonization to occur on the glass surface for 3 h. Then, we gently washed the wells with pre-warmed medium to fully remove unattached cells, after which we monitored further biofilm growth by timelapse imaging. We found that upon starting image acquisition, the biofilm-associated cell population and its extracellular matrix continued to grow for approximately 4.5 to 5 h, after which the biofilm-associated cells began to disperse into the surrounding medium while leaving the ECM structure intact. Subsequent exponential growth of the released cells produced a dramatic increase in GFP signal over time (Video 10 and Figure 6A).

**Figure 6.**
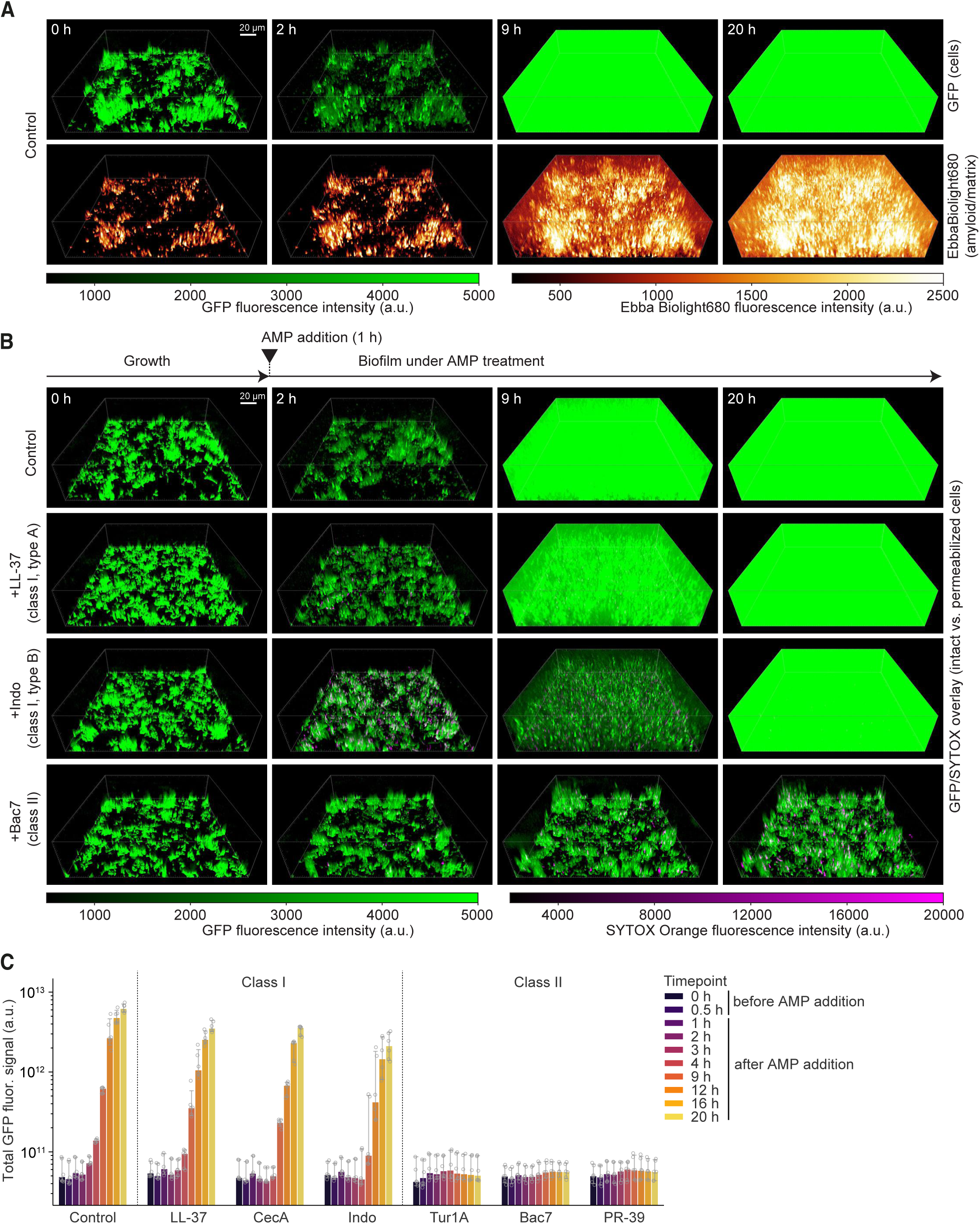
Class I AMPs are less effective against growing *E. coli* biofilms than class II peptides. **(A)** Selected timepoints of 3D side views of growing ATCC 25922GFP *E. coli* biofilms. Green signal (top) represents the GFP-expressing cells, whereas yellow/red signal (bottom) represents the matrix-staining EbbaBiolight 680 fluorogenic dye. Green signal saturation represents population overgrowth in the well upon biofilm maturation and dispersal. **(B)** Selected timepoints of biofilms treated with (or without) AMPs, showing overlaid GFP (cells) and SYTOX Orange (permeabilized cells) signals. AMPs were added at 2X MIC ∼1 h after the start of the image acquisition. **(C)** Bar plot showing the quantification of the total GFP signal, representing total cell biomass within the imaged 3D volume, over time for all the tested conditions. Fluorescence signals in (A-B) were rendered and exported from Imaris with gamma set to 2 for display only.

Next, we tested the effect of three AMPs from each class, added separately to biofilms at 2X MIC (measured for a diluted culture of ATCC 25922 cells, Figure S9A-B) after ∼1 h of growth since the start of image acquisition. SYTOX Orange was included to monitor IM permeabilization. EbbaBiolight 680 was omitted from these experiments, as this dye can bind intracellular polysaccharides once cells are permeabilized, confounding the signal. A no-AMP control experiment was performed in parallel under otherwise identical conditions. In this control, the biofilms grew normally, followed by cell dispersal and subsequent massive overgrowth, with little to no SYTOX Orange signal, confirming that IM integrity remained intact throughout (Video 11 and Figure 6B). All three class I AMPs (LL-37, CecA, and Indo) caused some IM permeabilization but failed to prevent biofilm growth or cell overgrowth (Videos 12-14 and Figures 6B and S9C). Indo, the class I type B AMP, marginally delayed overgrowth relative to class I type A LL-37 and CecA, in line with its slightly better performance in wide-trench experiments, but ultimately failed to prevent it. In contrast, all three class II AMPs (PR-39, Bac7, and Tur1A) effectively prevented biofilm growth and cell overgrowth (Videos 15-17, Figures 6B and S9C). Occasional floating cell clusters were observed landing on the glass surface during class II treatment, but they did not proliferate (Videos 15-17). Quantification of total GFP intensity confirmed that class II peptides prevented the growth of biofilms, unlike class I AMPs, which only slightly delayed it compared to the control (Figure 6C).

Collectively, these results indicate that the opposing trade-offs between AMP classes extend to biofilms. Biofilm resilience is commonly attributed to biofilm-specific gene expression programs, a protective ECM, and the presence of persister cells (K. Lewis, 2007; Kostakioti et al., 2013). Our data suggest that cell density per se is an additional, underappreciated contributor, one that operates independently of these biofilm-specific factors.

## DISCUSSION

In this study, we describe a time-resolved single-cell phenotyping method to study and classify AMPs and PMs. Temporal resolution is critical for classifying this family of antimicrobials, as diverse AMPs and mimics can produce the same cellular phenotypes but at different times relative to growth inhibition (Figures 1-4 and S5), a distinction that separates direct from secondary effects. Single-cell resolution is equally important given the cell-to-cell variability in event timing (e.g., membrane permeabilization) and the importance of probing the intracellular redistribution of molecules (e.g., mScarlet-I internalization, ribosome/DNA mixing) during AMP treatment (Figures 1-3). The combination of temporal and single-cell resolution also exposes a potential pitfall of conventional bulk assays, where membrane-impermeant dye uptake is often measured at the population level and at a fixed time point (e.g., 1 h) after AMP addition. Such measurements can be misleading: for instance, at 1 h post-treatment, SYTOX Green uptake appeared paradoxically lower for the class I type B peptide Indo than for the class II peptide PR-39 (Figure 1D). Realigning SYTOX Green uptake data for each cell relative to its time of growth inhibition resolves this discrepancy and restores a ranking consistent with the AMP class and subtype (Figure S2D).

Our comparative phenotyping analysis reveals that all examined AMPs, regardless of class or subtype designation, adhere to a set of common principles despite their remarkable diversity in sequence, structure, and characteristics (Figure 1A-B and S1A). All peptides displayed some membrane-permeabilizing activity, though at varying rates (Figure 1E). They also elicited shared intracellular effects, including a ribosome/DNA mixing event concurrent with growth inhibition (Figures 3 and S4), which mechanistic studies attribute to polysome loss and protein synthesis inhibition (Bakshi et al., 2012; Gray et al., 2019; P. J. Lewis et al., 2000; Linnik et al., 2024; Sanamrad et al., 2014; Xiang et al., 2021). In addition, all examined fluorescent AMP derivatives, irrespective of class affiliation, preferentially colocalized with ribosomes rather than the DNA (Figures 4B-D and S7B-D). This is consistent with structural studies implicating ribosomes as targets of class II AMPs such as Bac7 and Tur1A (Mardirossian et al., 2014, 2018). While our localization findings do not rule out additional intracellular targets, in vitro binding studies should be interpreted cautiously: cationic AMPs will associate with any anionic molecule in isolation. In fact, many AMPs including LL-37 and Tac1, have been shown to bind DNA in vitro (Yonezawa et al., 1992; Chongsiriwatana et al., 2017). However, fluorescent LL-37 and Tac1 derivatives were clearly excluded from DNA and instead colocalized with ribosomes (Figure 4B and S7C). Inside the cell, where highly abundant anionic components such as ribosomes compete for AMP binding, the in vivo target distribution may differ substantially from in vitro predictions. The electrostatic basis of this preference is further supported by the observation that engineering GFP to carry a high positive charge is sufficient to drive its association with ribosomes in *E. coli* (Schavemaker et al., 2017).

The distinction between the two classes of peptides lies in the kinetics of their membrane-permeabilizing activity. As noted above, the distinction cannot be simply reduced to one class being capable of permeabilizing membranes and the other not. While the frequency of IM permeabilization is generally high with class I AMPs (Figure 1E), it is not negligible with class II peptides, even at concentrations close to the MIC. Under our experimental conditions, the IM became permeabilized in the majority (> 60%) of cells treated with class II PR-39 or DJK-5 (Figures 1E and S5A). The key distinction is the timing. For class I, IM permeabilization occurs nearly simultaneously with the cell growth arrest, whereas for class II, it is generally absent or delayed relative to growth inhibition (Figures 1F, S2B, and S5B), even at high (10X MIC) peptide concentrations (Figure S6B-D).

These kinetic differences between classes give rise to major trade-offs in antibacterial activities, as illustrated in Figure S10A. At low cell density, class I AMPs/PMs offer two advantages: rapid action, particularly at high concentrations, and the ability to arrest growth through several mechanisms near simultaneously: IM permeabilization and subsequent inhibition of intracellular targets, such as ribosomes. However, this comes at the cost of rapid and massive intracellular absorption. At high cell densities, the first permeabilized cells act as “sponges” that deplete the extracellular pool of antimicrobials and protect other cells, rendering class I AMPs/PMs ineffective against dense populations. In contrast, the class II peptides inhibit growth more slowly because they rely at least in part on cellular transporters to gain access to their cytoplasmic targets (Mardirossian et al., 2018; Mattiuzzo et al., 2007). This slower uptake, however, leads to low intracellular absorption, allowing class II peptides to diffuse throughout dense populations and arrest growth even at high cell densities. These functional trade-offs likely reflect distinct host defense strategies adapted to different infection challenges, as both peptide classes can be produced by the same organism (e.g., bovine AMPs Indo and Bac7). For example, a fast-acting class I peptide may be advantageous to quickly neutralize an early infection when the bacterial burden is low, especially if the bacterial offender is motile and can quickly move away from the site of AMP release. In contrast, the low-absorption properties of a class II AMP likely confer an advantage over a class I if the infection has reached a high cell density or has developed into a biofilm. Thus, one class may be more efficacious than the other depending on the scale or type of infection.

Our findings also shed new light on the origin of biofilm resilience to immune attack. While this resilience is often attributed to the extracellular matrix and other biofilm-specific factors, our data implicate high cell density as an additional, independent contributor. The parallels between our biofilm and high-cell-density experiments (Figures 5, 6, S8, and S9) suggest that cell density alone is sufficient to confer biofilm resilience to class I AMPs. Our findings also carry implications for PM design: while high membrane-permeabilizing activity and self-promoted uptake are generally considered attractive properties for AMPs and their synthetic mimics, our results argue that in the context of biofilms, a low or delayed membrane-permeabilizing activity, as with class II peptides, is in fact preferable.

In the literature, membrane-permeabilizing AMPs are often described as “lytic”. However, Gram-negative bacteria possess two membranes, the OM and the IM, which have distinct chemical compositions. By using cytoplasmic and periplasmic fluorescent proteins with distinct spectral properties, our method demonstrates that differences in permeabilization susceptibility between the two membranes subdivide class I AMPs/PMs into two types (summarized in Figure S10B). Type A peptides permeabilize both membranes near the same time (Figure 2C-D and F), resulting in loss of both periplasmic and cytoplasmic reporters. In contrast, type B AMPs/PMs only permeabilize the IM at the time of growth arrest, keeping both periplasmic and cytoplasmic reporters within the boundaries of the intact OM (Figures 2E, 2G, S3A, S5E, and S5G). Thus, class I type B AMPs/PMs are IM-permeabilizing but not lytic, at least not at the time of growth arrest. This subtype distinction among AMPs/PMs suggests potential immunological implications, as the antibacterial activity of type A results in an immediate release of intracellular protein antigens, whereas type B does not.

It should be noted that our assay does not detect membrane permeabilization for cellular components (e.g., ions) smaller than SYTOX Green (< 600 Da). Furthermore, our classification framework is currently based on using *E. coli* as a bacterial target. Moving forward, it will be interesting to implement our approach on other bacteria, including Gram-positive bacteria, multidrug-resistant pathogens, and AMP-resistant strains. This will help refine the classification and expand our understanding of resistance mechanisms. Finally, our study highlights a few interesting sequence features in peptides with regard to their classification (Figures 1 and S10B). For example, the fact that proline-devoid, moderately hydrophobic DJK-5 falls into the same class II as the proline-rich, hydrophilic PR-39, Bac7, CamBac, and Tur1A cannot be trivially explained by sequence or structural similarity alone, suggesting that class II behavior can emerge from chemically diverse molecules. While our set of selected peptides is too small to draw conclusive relationships between sequence and function, a large-scale study integrated with machine learning models (Liu et al., 2025; Wan et al., 2024) holds the potential to bridge the gap between the sequence and function of antimicrobial peptides. Bridging this gap is essential to achieve predictive power and unlock their promise as next-generation therapeutics.

## METHODS

### Strains and growth conditions

For all experiments and *E. coli* strains, cells were grown in a medium (M9gluCAAT) with M9 salts (33.7 mM Na_2_HPO_4_, 22mM KH_2_PO_4_, 8.55 mM NaCl, 9.35 mM NH_4_Cl), 0.2% (w/v) glucose, 0.4% (w/v) casamino acids, 1 mM MgSO_4_, 0.3 mM CaCl_2_, 1 μg/mL thiamine, and trace elements (13.4 mM ethylene-diamine-tetra-acetic-acid (EDTA), 3.1 mM FeCl_3_-6H_2_O, 0.62 mM ZnCl_2_, 76 μM CuCl_2_-2H_2_O, 42 μM CoCl_2_-2H_2_O, 162 μM H_3_BO_3_, 8.1 μM MnCl_2_-4H_2_O). Prior to every experiment, cells were first inoculated into M9gluCAAT and grown to stationary phase at 37°C. For all assays except the biofilm experiments, the culture was diluted to 1/10,000 or more in fresh medium and grown at 37°C until it reached an optical density at 600 nm (OD) of 0.2-0.4. For the biofilm experiments, cells were grown in M9gluCAAT supplemented with 100 µg/mL ampicillin to maintain the GFP-expressing plasmid. The overnight stationary-phase cultures were then diluted into the same medium to an OD of 0.015. See below for details on biofilm growth.

Except for the biofilm experiments, the strains used in this study were derived from MG1655 (ATCC 47076), which was used to estimate the minimum inhibitory concentration (MIC) for all the studied peptides (or peptoids) using a microplate reader (Figure 1A). To detect the timing of outer and inner membrane permeabilization to proteins, strain CJW7845 was constructed to carry two freely diffusing fluorescent proteins: mScarlet-I in the periplasm and mTagBFP2 in the cytoplasm. Construction of CJW7845 was done by transforming pBAD-mTagBFP2 plasmid (Addgene #34632, a gift from Vladislav Verkhusha; (Subach et al., 2011)) into an *E. coli* strain expressing a periplasmic mScarlet-I reporter (Cañas-Duarte 2021), as previously described (Chung et al., 1989). Transformants were restreaked on fresh selection plates and confirmed via microscopy.

Strain CJW7753, which carries chromosomally expressed HupA-mCherry and RplA-GFP along with a plasmid-expressed cytoplasmic mTagBFP2 (Papagiannakis et al., 2025), was used to simultaneously monitor the intracellular perturbations of ribosomes and DNA over time while assessing the cell envelope integrity.

For both strains CJW7845 and CJW7753, mTagBFP2 expression was induced by supplementing the medium with 0.2% (w/v) arabinose. Ampicillin (100 μg/mL) was added to the growth medium of the liquid culture to maintain the plasmid-expressing mTagBFP2 but was not present in the microwell medium during the imaging experiment.

Strains CJW7020 (Gray et al., 2019) and CJW7859, which express RplA-msfGFP and HupA-msfGFP, respectively, were used to probe the spatial localization of ribosomes and DNA with respect to fluorescently labeled AMPs. Strain CJW7859 was constructed as follows. The monomeric superfolder GFP (msfGFP) coding sequence together with a kanamycin resistance cassette flanked with flippase recognition target (FRT) sites were amplified from a pKD13 vector (Datsenko & Wanner, 2000) encoding msfGFP without its start codon, preceded with a sequence encoding a glycine/serine-rich linker (GSGSGS) (strain CJW6871 (Gray et al., 2019)), using primers 5’-GGCATTTGTTTCTGGCAAGGCACTGAAAGACGCAGTTAAGGGCAGCGGCAGCGGCAGCTC-3’ and 5’-AAACCACCCCTTCGTTAAAACTGTTCACTGCCACGCAATCATCCGTCGACCTGCAGTTCG-3’.

Polymerase chain reaction (PCR) was performed to add homologous sequences to the 3-end region encoding the HupA C-terminus and the downstream region and to remove the *hupA* stop codon, yielding an encoded C-terminal HupA-msfGFP fusion. The amplified product was transformed by electroporation into MG1655 containing pSim6. The plasmid expresses the Red recombineering proteins (encoded by *exo*, *bet*, and *gam*) under the control of the native λ phage pL promoter and temperature-sensitive cI857 repressor, whose expression can be induced by heat-shock as previously described (Sharan et al., 2009). The pSim6 plasmid was kindly provided by the National Cancer Institute. Briefly, cells were grown at 30°C to mid-exponential phase and the synthesis of the recombineering proteins was induced for 15 min at 42°C prior to making cells electrocompetent. Mutants were selected on kanamycin, and the correct integration was verified by PCR and sequencing of the modified genomic region using primers 5’-TGTCGTACCTGGAGTCTTCC-3’ and 5’-TTCAGCAAATCCACTGGCGG-3’. The modification was moved to a clean wild-type MG1655 strain in which no recombineering system was expressed via P1 transduction as previously described (Thomason et al., 2007). Transductants were confirmed via PCR and sequencing of the modified region using the same primers as above. A single colony was grown and transformed with temperature-sensitive pCP20, expressing the site-specific Flp recombinase to excise the kanamycin cassette used for initial selection, as previously described (Cherepanov & Wackernagel, 1995). Briefly, transformants were selected on LB plates containing chloramphenicol (30 μg/mL) at 30°C and streaked on LB plates at 42°C overnight to induce Flp recombinase expression and loss of pCP20 plasmid. The loss of FRT-flanked kanamycin resistance cassette was confirmed by PCR and the lack of growth in the presence of kanamycin (50 μg/mL). The loss of pCP20 was confirmed by the lack of chloramphenicol resistance.

Strain ATCC 25922GFP (ATCC, Virginia, USA), which carries a GFP-expressing plasmid, was selected for biofilm experiments. The strain’s ability to form biofilms was first verified at 37°C using Congo Red staining on agar plate containing M9gluCAAT. Congo Red dye binds curli, a major component of the biofilm extracellular matrix. Strain ATCC 25922 (lacking the GFP-expressing plasmid) was used for MIC determination (to avoid interference from the presence of ampicillin required to maintain the plasmid) following the standard growth protocol described above.

### Sources and preparation of antimicrobial peptides/peptoids

Lyophilized powders of LL-37 (AS-61302), human β-defensin 3 (AS-60741), magainin 2 (AS-20639), cecropin A (AS-24009), indolicidin (AS-60999), protegrin-1 (AS-64819) were purchased from AnaSpec Inc. (California, USA). PR-39 (350590) was purchased from Abbiotec (California, USA). CRAMP (CRB1000262) was purchased from Biosynth Ltd (UK). Tachyplesin I (4030734) was purchased from Bachem AG (Switzerland). Camel bactenecin (71266-2), bactenecin 7 (318782), Tur1A (318733), Tur1B (319842), and DJK-5 (318921) were purchased from NovoPro Bioscience Inc. (Shanghai, China). 5-TAMRA-labeled cecropin A, tachyplesin I, LL-37, PR-39, Bac7, and DJK-5, were commissioned to NovoPro Bioscience Inc. (Shanghai, China) for custom peptide design. For all fluorescently labeled peptides, the fluorescent label was conjugated to the N-terminus. Peptide powders were resuspended in Milli-Q water to stock concentrations of 1 mM (or 0.5 mM for human β-defensin 3), aliquoted, and stored at −20°C. TM1 peptoid (first described as Peptoid 1), TM4, and TM5 were synthesized on an automated peptide synthesizer and purified by reversed-phase high-performance liquid chromatography, as previously described (Chongsiriwatana et al., 2008, 2017). Prior to each experiment, aliquots were diluted in fresh medium to the desired concentration.

### Protein sequence analysis and feature extraction

Amino acid sequences of the selected peptides (Figure 1A) were analysed using custom-written functions in Python to extract physicochemical properties (protein_analysis.py). The GRAVY index was computed using the Kyte-Dolittle scale of amino acids hydropathy values (Kyte & Doolittle, 1982). Polar and hydrophobic fractions were calculated based on the following classification: serine, glutamine, asparagine, glycine, threonine, proline, tyrosine, lysine, arginine, histidine, aspartic acid, and glutamic acid were considered polar amino acids whereas alanine, isoleucine, leucine, methionine, phenylalanine, tryptophan, cysteine, and valine were considered hydrophobic. Net charge calculation included contributions from lysine, arginine, and C-terminal amidation (when present). Molecular weight was computed starting from the sum of the molecular weights of the *n* single amino acids, subtracting *n-1* water molecule’s weights (18.015 g/mol) as a result of peptide bond formation, weight loss due to C-terminal amidation (0.98 g/mol, when present), and weight loss due to disulfide bridge formation (2.02 g/mol per cysteine pair).

### Determination of minimal inhibitory concentrations and maximum growth rates

To determine the minimal inhibitory concentration (MIC) of the selected AMPs against either MG1655 or ATCC 25922 *E. coli* strains, exponentially growing cells were diluted in fresh medium to an OD = 10^-6^ (∼500 cells/mL) and mixed 50:50 with titrated AMP concentrations (which were chosen to be over a range that included the MIC) in a 96-well plate (flat bottom with low evaporation lid, 3595, Corning, Arizona, USA). A final OD of 5 x 10^-7^ corresponds to ∼250 cells/mL. OD curves were sampled every 5 min for a total of 48 h using a microplate reader (BioTek Synergy 2, Agilent Technologies Inc., California, USA). Raw data were exported as a text file and analysed using a custom-written Python script (microplate_reader_analysis.py). To measure maximum growth rates, the log(OD) was fitted using a linear regression over a 12-point moving window (60 min), which provides the instantaneous growth rate (log(OD)/h), from which the global maximum was computed. The MIC was defined as the minimum AMP concentration for which the maximum growth rate was < 0.2 log(OD)/h within the first 20 h of incubation. Each MIC experiment was performed independently at least three times (with at least two technical replicates each), and the mode across all replicates was taken as the final MIC; in the rare cases of a tie, an additional replicate was performed, which was sufficient to resolve it.

### Fabrication of a multiwell microfluidic device for single-cell analysis

The multiwell microfluidic device (Figure S1D) was built as follows. A solution of 9:1 polydimethylsiloxane (PDMS) and curing agent was first mixed and degassed to remove air bubbles. A silicon rubber O-ring (25.8 mm inner diameter, 3.1 mm in thickness) was glued onto a glass bottom dish (40 mm diameter, 14026-20, from Ted Pella Inc., California, USA). PDMS solution (1 gr) was poured onto the glass dish within the O-ring to form a ∼1 mm thin PDMS layer, and cured for > 3 h at 64.5°C. An 8 mm biopsy punch was used to perforate the PDMS thin layer and carve out four independent ∼70-μL wells. The four wells were thoroughly rinsed using 70% (v/v) ethanol and 100% isopropanol and blow-dried using purified air. Cells were immobilized using a protocol for atomic force microscopy (Benn et al., 2019) with some modifications. Briefly, the glass surfaces were rendered hydrophilic using air plasma treatment (45 s at 60 W). Then, a Cell-Tak^TM^ (354240, Corning^®^, Arizona, USA) solution was prepared by mixing 250 μL of 0.1 M sodium bicarbonate, pH 8.0, 10 μL Cell-Tak^TM^, and 5 μL 1M NaOH (enough to functionalize four wells). This solution (50 μL) was applied to each well for 5 min, rinsed three times with Milli-Q water and blow-dried with purified air. Half a microliter of exponentially growing cells (diluted to OD_600_ ∼ 0.15) was then spotted into the center of each well for 5 min to allow for cells to be immobilized on the surface. The small 0.5-μL volume allowed us to immobilize cells only within a small region of the well, thus minimizing cell density effects that may affect the MIC. The wells were then washed three times with fresh medium to remove the cells that did not attach to the surface, after which each well was filled with growth medium (70-80 μL). To minimize water evaporation from the wells during the experiments, ∼1 mL of medium was added to the region surrounding the O-ring, which aided in maintaining a high local humidity in proximity of the wells. Experiments were carried out with the lid on.

### Fabrication of wide-trench microfluidic device

The design of the wide-trench microfluidics was adapted from the one reported by (Lambert & Kussell, 2014). As reported before (Lin & Jacobs-Wagner, 2022), the master mold was manufactured in the clean room of the School of Engineering and Applied Science at Yale University. UV photolithography (EVG 620 Mask Aligner) was applied on Si wafers with SU-8 photoresist to generate two layers of positive patterns. The lower layer contained the bacterial culture chambers (wide trenches) while the upper layer contained the large trench for the flowing culture medium. The large feeding region features a ∼200 μm width and ∼30 μm height. The wide trenches have a width of 7 μm, a depth of ∼34-38 μm, and a height of 1.29 μm.

### Wide-trench microfluidics chip preparation and experiment

A solution of 9:1 polydimethylsiloxane (PDMS) and curing agent was mixed, degassed to remove air bubbles, poured onto the silicon wafer, and cured at 64.5°C for > 3 h. Note that the 9:1 PDMS:curing agent ratio was used instead of the standard 10:1 to reduce the hydrophobicity of the PDMS and mitigate hydrophobic interactions with AMPs. Once hardened, PDMS chips were cut out from the wafer, and a 1.2 mm biopsy punch was used to create inlet and outlet holes. The current chip design features two independent channels per chip. To allow for a higher temperature during PDMS-glass bonding, the glass bottom part of the dish was used during the following steps (without the plastic support glued to it). Both glass slide and PDMS surfaces were subsequently rinsed with 70% (v/v) ethanol and isopropanol to remove dust and debris and blow-dried with purified air. The glass slide was then placed on a 95°C hot plate for further drying for ∼2-3 min. Subsequently, the PDMS chip and glass slide were plasma-treated (for 20 s at 60 W), immediately bonded, placed on a 95°C hot plate for 5 min, and baked at 75°C for ∼2 h. The bonded chip was then left to equilibrate at room temperature, and a solution of M9gluCAAT supplemented by 0.08 % Pluronic® F-108 was perfused through the channels, which served to wet the dry PDMS and glass surfaces and to render them more hydrophilic by passivating with Pluronic (which has amphiphilic properties). Pluronic (0.08%) was also supplemented into the growth medium to improve the hydrophilicity of the cell outer membranes, with the intent to reduce cell aggregates. Exponentially growing cells (3 mL) were spun at 6500 rpm for 5 min, resuspended in 50-100 μL (to increase cell density), and loaded into the channels. The cells-loaded chip was then spun at 500 g for 3 min using a three-dimensional-printed holder (design gifted by Dr. Johan Paulsson) to facilitate entry of the cells into the trenches. Before every experiment, all the tubing were sterilized and cleaned by flowing 20% (v/v) bleach, 20% (v/v) ethanol, Milli-Q, and then perfused with 0.08% Pluronic-supplemented medium. Experiments were carried out at 37°C unless indicated otherwise. Flow was achieved by a peristaltic pump (T60-S2CWX10-14-H, Langer Instrument, USA), which was set to deliver a constant flow rate of ∼35 μL/min, which was maintained for the whole duration of the experiment, including the AMP treatment.

### Biofilm growth and sample preparation

For the biofilm experiments, ATCC 25922GFP was grown overnight to stationary phase at 37°C in M9gluCAAT supplemented with 100 µg/mL ampicillin (to maintain the GFP-expressing plasmid). The overnight culture was then diluted into fresh, pre-warmed (37°C) M9gluCAAT containing 100 µg/mL ampicillin to OD=0.015, and 300 µL of the diluted culture was added to each well of an 8-well chambered cover glass (#1.5H high-performance cover glass, 0.17 mm; C8-1.5H-N; Cellvis, California, USA). Biofilms were grown statically for 3 h at 37°C. Biofilms were then washed once by gently removing the medium, adding 400 µL of fresh pre-warmed medium, removing it, and replacing it with 300 µL of fresh pre-warmed medium (without ampicillin). This step served two purposes: it removed most planktonic cells and floating cell clusters that would otherwise deposit on the biofilm during acquisition, and, by decreasing the overall cell density of the well, it prolonged the growth phase of the biofilm prior to its maturation and cell dispersal. When extracellular matrix formation was monitored, the fluorogenic optotracer EbbaBiolight 680 (Ebba Biotech, Sweden), which labels bacterial amyloids such as curli as well as certain matrix polysaccharides, was added to the medium at a 1:500 dilution.

### Microscopy

Images were acquired using a Nikon Ti2-E inverted microscope equipped with a Photometrics Prime BSI back-illuminated sCMOS camera (2048×2048 pixels, 6.5 μm pixel size), a 100X Plan Apo 1.45NA Ph3 oil objective (with type N immersion oil by Nikon), a Lumencore Spectra III LED (Light Emitting Diode) engine, a Perfect Focus System (PFS), temperature-controlled Okolab enclosure, and a polychroic mirror (FF-409/493/596-Di02 by Semrock) combined with a triple-pass emitter (FF-1-432/523/702-25 by Semrock) for mTagBFP2/GFP/mCherry. mTagBFP2, GFP (or SYTOX Green), mCherry (or mScarlet-I) were imaged using excitation at 365 nm, 488 nm, and 594 nm, exposure of 300 ms, 200 ms, and 200 ms, and a light emitting diode (LED) power of 75%, 20%, and 30%, respectively (see (Papagiannakis et al., 2025) for power measurements for the different light sources). A 5% transmission neutral density filter was also used to lower the LED power and reduce photobleaching and phototoxicity. All experiments were carried out at 37°C unless indicated otherwise.

The microscope was controlled using NIS Elements software (Nikon), and an N-Dimensional (ND) acquisition module was used to set and acquire the time-lapse images. Phase and fluorescence images were acquired every 1 min for the single-cell and wide-trench analyses or 10 min for some of the experiments with fluorescently labeled AMPs.

For experiments with PDMS microwells, cells were allowed to grow for ∼30 min in the temperature-controlled enclosure prior to imaging. A 10X concentrated AMP solution was spiked in (at 1/10 the volume of the well) and mixed well by pipetting after ∼20 min from the start of the image acquisition, without interrupting the timelapse sequence. When the uptake of SYTOX Green was monitored, SYTOX Green was included in the AMP solution and spiked in during the experiment to obtain a final concentration of SYTOX Green of 50 nM in the well. This allowed us to monitor healthy cell growth prior to AMP treatment, to rapidly set the desired AMP (or SYTOX Green) concentration from one frame to the next (unlike for flow-controlled injection systems), and to resolve the AMP-induced phenotype in real time. The current design allows for testing up to four independent conditions in parallel.

For the biofilm imaging experiments, images were acquired on the same Nikon Ti2-E microscope described above, but using a 40X air Plan Apo 0.95NA air objective. As for the other experiments, a 5% transmission neutral density filter was kept in the light path to limit photobleaching and phototoxicity. Excitation intensities and exposures differed from the single-cell experiments: GFP was imaged using 488-nm excitation at 10% power, whereas the SYTOX Orange or amyloid stain EbbaBiolight 680 was imaged using 594-nm excitation at 50% power; images from both channels were acquired with a 100 ms exposure. To capture the three-dimensional (3D) structure of the biofilm, z-stacks of 73 planes were acquired with a 0.6 µm step size (∼43 µm total range), and the PFS was used to maintain focus over the course of the time-lapse. Z-stacks were collected every 30 min for 45 frames (∼22 h total) at 37°C. The imaging enclosure was pre-warmed to 37°C and a 50 mL water bath was placed inside it to mitigate evaporation.

In 5-TAMRA-AMP enrichment experiments (Figures 4A and S7A) that used strain MG1655, unlabeled AMPs at 2X MIC were supplemented with 50 nM of the corresponding 5-TAMRA-AMP conjugate, to have an equal bulk concentration of the fluorescent label for quantifying intracellular fluorescence increase over time. In colocalization experiments that used strains CJW7020 and CJW7859 (Figure 4C), unlabeled AMPs at concentrations equivalent to 2X MIC (or 1.5X MIC for PR-39) were supplemented with 5% of the corresponding 5-TAMRA-labeled AMP. Only for PR-39 and Bac7 and only for estimating the SCF* (Figure 4C), the labeled:unlabeled concentration ratio was increased to 25:75 and 50:50, respectively, to increase the fluorescent signal and improve the calculation of the SCF*. Selected images in Figures 4B, 4D, and S7B-D were chosen at times where both signal intensities from fluorescently labeled AMPs and cytoplasmic macromolecules (DNA or ribosomes) were the highest to aid visualization of both molecules. The different timepoints for the different fluorescently labeled AMPs or macromolecules reflect the difference in uptake kinetics or stability of the label over time (for example, HupA-msfGFP signal under Tac1 treatment exhibited a fast decay as a result of cell permeabilization).

In wide-trenches microfluidics experiments, CJW7753, CJW7845, or CJW7859 cells were allowed to grow for > 1 h with running fresh medium in the temperature-controlled enclosure to reach steady-state conditions prior to imaging. When present, SYTOX Green was added with the AMPs at 5 nM.

### Image processing and analysis of PDMS-microwells single-cell data

To analyse the single-cell experiments in microwells, OmniSegger software (https://github.com/tlo-bot/omnisegger), which combines SuperSegger MATLAB-based suite (Stylianidou et al., 2016) for cell analysis with Omnipose segmentation (Cutler et al., 2022), was used to first process the microscopy images by performing drift correction, cell segmentation, and cell linking. Given that AMP-treated cells can display extreme phenotypes, such as exceedingly bright and/or heterogeneous phase contrast signal (likely a result of the widely changing intracellular density upon AMP treatment) and that the software’s pre-trained model *bact_phase_omni* (Cutler et al, 2022) tended to split cells before cell division was completed, a new Omnipose model was trained from scratch starting from *bact_phase_omni*-generated masks that were further corrected and curated using custom-made Python scripts (screening_good_bad.py and generate_training_dataset.py). Details on the Omnipose model retraining can be found in a previous publication (Thappeta et al., 2025). The final model (omnipose_phase_microwells_model) and corresponding training datasets are available at https://github.com/JacobsWagnerLab/published/tree/master/Fragasso_et_al_2025 and BioImage Archive (accession number: S-BIAD1823), respectively.

MATLAB files of the linked cell trajectories output from OmniSegger were then imported, background subtracted using background_subtraction.py, which was adapted from a previous study (Papagiannakis et al., 2025), and converted to Python data structures (import_processing_code.py), from which cell features such as cell area, nucleoid area, mean fluorescence intensity, etc., were extracted for each cell at each frame. The exported dataframes were further processed to estimate instantaneous growth rates and were run through a series of functions to perform curation based on cell areas and instantaneous growth rates over time (post_processing_curation_functions.py). Cell trajectories (i) that were too short (< 30 min), (ii) that exhibited exceedingly large or small values in normalized (by area) instantaneous growth rate (> 0.04 min^-1^ or < −0.02 min^-1^), (iii) whose first ten frames did not feature normal healthy growth (< 0.005 min^-1^), or (iv) whose last ten frames did not feature growth arrest (> 0.005 min^-1^), were removed from further analysis. In this way, only cells that exhibited healthy growth followed by growth arrest were analysed, namely those cells for which the growth arrest-associated AMP phenotype was fully resolved. Represented data points were pooled from multiple replicates.

### Estimation of the timing of membrane permeabilization events relative to the growth rate inhibition

Analysis of strain CJW7845 proceeded by extracting the timings of permeabilization of the outer and inner membranes based on the mean fluorescence intensities of mScarlet-I and/or mTagBFP2 fluorescent proteins (using function get_permeabilization_time.py), cell growth arrest, and the initiation time of growth inhibition. To obtain accurate timings underlying membrane permeabilization events, information from both the signal intensity itself as well as its derivative were combined, which enabled us to accurately resolve the relevant changes in fluorescent protein signal beyond the intrinsic high-frequency noise and low-frequency photobleaching. Inner membrane permeabilization to proteins can result in two possible phenotypes when simultaneously monitoring freely diffusing periplasmic and cytoplasmic probes, depending on which membrane (outer or inner) becomes permeabilized first. When the outer membrane is first permeabilized, mScarlet-I signal is lost first, followed by mTagBFP2 when the inner membrane becomes also permeabilized. Conversely, when the inner membrane is first permeabilized, mScarlet-I diffuses into the cytoplasm and mixes with mTagBFP2. To detect inner membrane permeabilization when the outer membrane remained intact (i.e., mScarlet-I leaks into the cytoplasm), the difference between periplasmic and cytoplasmic contributions of the mScarlet-I signal was computed, where periplasmic contribution corresponded to the mean fluorescence intensity of mScarlet-I within the 5-pixel thick outermost layer of the cell mask, while the cytoplasmic contribution was calculated by averaging pixel fluorescence intensities within a 7-pixel eroded mask. This way, the obtained difference Δ(peri, cyto)_mScarlet-I_ remains overall positive when mScarlet-I is localized in the periplasm and becomes rapidly negative when the inner membrane becomes permeabilized as mScarlet-I diffuses into the cytoplasmic region. Detection of the permeabilization event relied on both Δ(peri, cyto)_mScarlet-I_ and its normalized first derivative, which was computed by using a 5-point moving window to evaluate the local slope across the signal and dividing it by the signal itself.

When IM was permeabilized after the OM, detection of such event relied on detection of the rapid decrease of the mScarlet-I signal first (OM permeabilization), followed by the loss of the mTagBFP2 (IM permeabilization). Both signals were treated in the same way as Δ(peri, cyto)_mScarlet-I_ for detecting permeabilization events, respectively, namely by using information from both the signal itself as well as their normalized first derivative.

The time marking the initiation of growth inhibition and the growth arrest were obtained by defining a multi-scale instantaneous growth rate with weights (multi_scale_derivative.py). Specifically, this function computes the weighted average of the first derivatives of differently smoothened cell areas, where these were computed using a Savitzky-Golay filter with varying moving windows. The normalized growth rate *GR*_*norm*_(*t*) can be expressed as

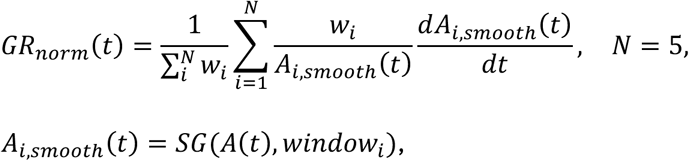

where *w*_*i*_ is the *i*^*th*^ weight, *A*_*i,smooth*_(*t*) is the *i*^*th*^ smoothened area signal, SG = Savitzky-Golay filter (Savitzky & Golay, 1964) (implemented using the Python function *scipy.signal.savgol*), *window*_*i*_ is the *i*^*th*^ moving window size for smoothing, *A*(*t*) is the original cell area signal. For the current implementation, window sizes were set to be 3, 5, 7, 9, 11, with corresponding weights 5, 8, 10, 8, 5, which were found by empirical optimization, to capture growth arrest both in the case of fast cell shrinking events (induced by class I AMPs), where growth rate changes abruptly between one frame and the next one, as well as when growth is slowly inhibited (e.g., induced by Class II AMP treatment). As displayed in Figure S2A, the start of growth inhibition is defined as the first frame where *GR*_*norm*_(*t*) ≤ 0.01 min^-1^ and remains stably below 0.012 min^-1^ for at least 30 consecutive frames. Growth arrest is defined as the first frame where *GR*_*norm*_(*t*) ≤ 0 min^-1^ and remains stably below 0.005 min^-1^ for at least 30 consecutive frames. The use of a higher threshold to evaluate stability over consecutive frames was necessary to account for noise fluctuations of the *GR*_*norm*_(*t*) signal. Scatterplots presented in Figures 1F, 2C, 3D, 3G, S5B, S5D, S6C, and S6G were obtained using these different metrics.

### Computation of the single-cell kymographs

To compute the kymographs (Figures 2D-E, S3, S5E, and S5F), a function was first used to compute the medial axis (smooth line longitudinally crossing the mid-cell) from each cell mask (get_medial_axis.py). Briefly, the medial axis was computed by first upsampling the cell mask by 20-fold in both x and y. Upon smoothing of the cell mask boundary, the Python function *skeletonize* (from *skimage.morphology*) was then used to trace the cell skeleton, which roughly corresponds to the set of pixels that are the farthest away from the cell boundary. To find the cell poles, linear regressions were fitted to the computed skeleton’s extremities, specifically to the first and last three-eighths of its length, and the intersections between the extrapolated lines and the cell boundary were identified as the cell poles. Finally, a third-order polynomial function was fit to the skeleton points and the poles to compute the medial axis.

Upon defining the medial axis, each cell pixel (x,y) coordinates could then be mapped into (l,d) medial axis coordinates (l = length along medial axis and d = minimum distance from medial axis; get_1D_proj_medial.py). Mean intensity projections along the medial axis length of the pixels with d < 4 pixels were computed by averaging together the pixels with the same l coordinate (kymograph_single_cell.py). Given that medial axis coordinates were computed with sub-pixel resolution using a 20X augmentation factor, the medial axis length was binned using equal bins with the original pixel size (0.065 μm/px). The resulting mean intensity projections along the medial axis length were then plotted over time to obtain the final kymographs.

### Analysis of intracellular perturbations upon AMP uptake

To analyse strain CJW7753, the signal correlation factor (SCF) was computed by estimating the pixel-based Spearman correlation between the RplA-GFP and HupA-mCherry signal over a 4-pixel eroded cell mask. Exclusion of the outer 4-pixel cell layer was necessary to remove the effect of cell curvature in the proximity of the cell edge, which causes both fluorescence signals to decrease in intensity (thus causing an artefactual positive correlation). To calculate the nucleocytoplasmic (NC) ratio (= ratio between nucleoid length and cell length), a mask was applied to the HupA-mCherry signal to determine the nucleoid region (nucleoid_mask.py). Then, projections of the nucleoid pixel coordinates onto the cell medial axis were computed to estimate the arc length, namely the nucleoid length, which was obtained by calculating the Euclidean distance between consecutive pairs of pixels along the nucleoid projections. The arc length of the full medial axis, namely the cell length, was estimated using the same approach (nucleoid_cell_length.py). Cell trajectories were then aligned by the SCF peak using a multi-step alignment approach (time_alignment.py). Given that SCF signal can generally have multiple local maxima during the evolution of the phenotype after growth arrest (due to, for example, progressive loss of cell biomass, which decreases de-mixing between DNA and ribosomes, thus increasing SCF), cell trajectories for class I peptides (which featured a global peak in area that is time-associated with the SCF peak) were first aligned by peak area (here the maximum value of the area), which allowed us to first roughly synchronize the trajectories by growth arrest. Then, the maximal SCF value (max SCF) within a narrow time window (± 5-10 min) from the peak area was used to align the trajectories by the peak SCF value, resulting in all cells being synchronized by the start of their intracellular perturbations (peak SCF). For class II peptides, since the area global maximum is not generally associated with cell growth arrest, the minimum NC ratio was first used to align trajectories, then the max SCF value was searched within a narrow time window (± 5-10 min) to obtain the peak SCF value. Given that for class II AMPs (PR-39, Bac7, Tur1A, and CamBac), the SCF peak is broad (tens of minutes) compared to noise fluctuations, a smoothened SCF (using a 12-point moving average window) was used for both determining the peak SCF (which improved robustness against noise) and computing the ensemble curves in Figures 3F and S5. The time alignment step resulted in a new time axis that featured a SCF peak at t = 0 and which was used to generate the ensemble plots in Figures 3C, 3F, and S4. From this, the timing of peak SCF could also be extracted, which was used in the scatterplots in Figures 3D and 3G.

### Analysis of fluorescently labeled AMP uptake and spatial localization

For Figures 4A and S7A, images of MG1655 cells were acquired every 1 min to enable accurate cell tracking (using OmniSegger) and probe the intracellular 5-TAMRA-AMP enrichment as a function of time aligned by the start of growth inhibition. The analysis steps for single-cell analysis were performed as explained in **‘**Image processing and analysis of PDMS-microwells single-cell data’ section.

In contrast, to analyse the localization of fluorescently labeled AMPs (Figure 4C) in CJW7020 cells (expressing RplA-msfGFP) or CJW7859 cells (expressing HupA-msfGFP), images were acquired every 10 min to increase the number of fields of view, and thereby the n value, per condition since the focus of this experiment was to correlate the fluorescently labeled AMPs’ spatial localization with fluorescently labeled cytoplasmic macromolecules (ribosomes and DNA) after uptake, without the need to track single cells over time. For this reason, OmniSegger was used here only to perform drift correction and cell segmentation, whereas mask labeling, cropping of individual cell regions, and feature extraction were performed using a custom-made Python script (import_processing_no_linking_code.py). Since no single-cell growth rate information was obtained, the curation of badly segmented or out-of-focus cells was performed using a different strategy. First, hard thresholds on cell area removed objects that were too large (> 2000 pixels) or too small (< 500 pixels) to be cells. Second, out-of-focus cells were removed by computing the average of the pixel intensities in the 40^th^ percentile of the pixel intensities in the peripheral region of the phase contrast signal, where the peripheral region corresponded to the outermost 1-pixel thick layer of the mask, computed from a 1-pixel eroded mask (the outer 1-pixel thick layer of the original mask was thus removed to avoid cell curvature effects). Third, cells that displayed low mean fluorescence values for either fluorescently labeled AMPs (< 100 a.u. for 5-TAMRA-labeled AMPs) or msfGFP-tagged macromolecules (< 100 a.u.) were filtered out to include mostly cells that were sufficiently enriched with fluorescently labeled AMPs. To compute the spatial correlation between fluorescently labeled AMPs and the ribosomes or DNA markers (Figure 4C), a modified version of SCF (SCF*) was defined to suppress the contributions from the region of the cell where both signals were low in intensity. Starting from a 1-pixel eroded mask, pixel-based Spearman correlation was computed between the pixel intensities in the 50^th^ percentile of both channels. The resulting SCF* appears to represent more strongly the visually observed anti-colocalization between fluorescently labeled AMPs and the DNA marker HupA-msfGFP, while providing positive correlation values when the signals are colocalized (as reported in Figure 4C for SCF* between fluorescently labeled AMPs and the ribosome marker RplA-msfGFP). The timepoints for evaluating the SCF* were chosen based on having a sufficiently high signal for both fluorescent reporters. The timepoints (relative to addition of the AMP mixture) were 10 min, 50 min, 100 min, 50 min, 30 min, 40 min, and 80 min, for 5-TAMRA-CecA, 5-TAMRA-Tac1, 5-TAMRA-LL-37, 5-TAMRA-PR-39, 5-TAMRA-Bac7, and 5-TAMRA-DJK-5, respectively, for RplA-msfGFP-expressing CJW7020, and 0 min (first frame after AMP addition), 50 min, 100 min, 50 min, 30 min, 40 min, and 80 min, for 5-TAMRA-CecA, 5-TAMRA-Tac1, 5-TAMRA-LL-37, 5-TAMRA-PR-39, 5-TAMRA-Bac7, and 5-TAMRA-DJK-5, respectively, for HupA-msfGFP-expressing strain CJW7859.

### Omnipose model training for segmentation of wide-trench data

To quantify single-cell growth rates in wide-trench microfluidics experiments, dedicated Omnipose models were trained for this imaging format using information from both phase-contrast and ribosome-fluorescence images. First, an initial training set was generated by applying the pre-trained *bact_ffuor_omni* model to the ribosome channel, manually curating the resulting masks, and pairing these masks with the corresponding phase-contrast images. This dataset was used to train a preliminary phase-contrast wide-trench model from scratch using 353 images, 4000 epochs, a learning rate of 0.1, GPU acceleration, and 224 by 224-pixel training tiles.

This preliminary model was then applied to wide-trench images from multiple AMP treatments to expand the training set across AMP-induced phenotypes. After manual curation against the ribosome signal, a final ribosome-channel model (omnipose_ribo_wide-trench_model) was generated by fine-tuning bact_fluor_omni using 592 curated images, each containing four wide trenches, with the same training parameters. Finally, an improved phase-contrast model (omnipose_phase_wide-trench_model) was trained from scratch using 868 curated images generated by applying the ribosome-channel model to a subset of wide-trench data and screening the resulting masks against phase-contrast images. Although both phase-contrast and ribosome-channel models were generated, the final ribosome-channel model was used for the single-cell segmentation in Figures 5C-F, 5I, and S8A-B and Videos 5-6, because it performed more robustly across treatment conditions and cellular phenotypes. The phase-contrast model was instead used to define the cell-containing regions for generating the SYTOX Green and LL-37 kymographs (Figures 5G-H and S8C-E). The final models and corresponding training datasets are available at https://github.com/JacobsWagnerLab/published/tree/master/Fragasso_et_al_2025 and BioImage Archive (accession number: S-BIAD1823), respectively.

### Wide-trench single-cell analysis of growth rate

To analyse wide-trench time-lapse images, drift correction was first performed using OmniSegger, followed by a custom Python script (drift_correction_wide_trenches.py) to correct for the slow residual drift that remained after the first correction step. The resulting drift-corrected timelapse data were analysed with the OmniSegger pipeline using the ribosome-fluorescence wide-trench Omnipose model described above, allowing cell segmentation and cell linking to be performed in the same general manner as for the PDMS-microwell experiments.

The MATLAB files containing linked cell trajectories were imported and pre-processed using custom Python code (import_processing_code_wide-trench.py). Cell trajectories were then curated using post_processing_curation_code_wide-trench.py to remove trajectories that were too short (< 15 min), that exhibited exceedingly large or small values in normalized-by-area instantaneous growth rate (> 0.04 min^-1^) or < −0.03 min^-1^), or that displayed large instantaneous center-of-mass displacements (> 50 px). This curation step was used to retain cells that were tracked for a sufficient number of frames to estimate meaningful growth rates, while excluding poorly linked trajectories or incorrectly segmented cells.

To compute the multi-trench kymographs for growth rate (Figures 5D-F, 5I, and S8A-B), from the curated single-cell trajectories, instantaneous growth rates were estimated and pooled across multiple trenches and/or biological replicates. Time was offset relative to the start of AMP injection, such that (t = 0) min corresponded to AMP addition. To visualize spatially resolved growth dynamics inside the wide trenches, kymographs were computed using custom Python code (compute_kymo_growth_rate_wide-trench.py). For each cell at each time point, the y-coordinate of the cell-mask center of mass was binned into 12-px (0.79 µm). To remove isolated spurious cells or poorly supported bins from the kymographs, occupied time-position bins were pruned using an iterative neighborhood filter. The resulting kymographs were plotted as the mean normalized growth rate as a function of time and position along the trench. To generate the single-trench images or videos (Figures 5C and Videos 5-6), the script plot_single_trench_growth_rate.py was used to import growth rate single-cell data from individual trenches and combine with image sequences of the corresponding trenches. Generation of mp4 videos was carried out with Python written code export_mp4_wide-trench.py.

### Kymograph generation and analysis of SYTOX Green and LL-37 fluorescence in wide trenches

Kymographs of SYTOX Green and LL-37 fluorescence in wide trenches (Figures 5G-H and S8C-E) were generated using a custom Python script (compute_kymo_fluor_wide-trench.py, plot_kymo_fluor.py) that rely on a shared functions library (quantify_fluor_kymo_functions.py). After applying drift correction (as described above), fluorescence and phase-contrast images were first background-subtracted. To identify and restrict the phase-contrast signal to cell-containing regions, dilated Omnipose-derived masks were used. Wide trenches were detected from phase-contrast images by analysing the mean projection of the phase-contrast signal onto the x-axis and using a peak detection algorithm to identify the trench vertical edges. Similarly, the top and bottom of the trench were also detected. These trench coordinates were then used to define the region of interest (ROI) for extracting fluorescence and phase-contrast signal intensities from each frame.

For each trench and time point, fluorescence and phase-contrast signals were projected onto the trench’s y-axis by averaging pixel intensities, excluding 50 px from each side of the detected trench to avoid edge artifacts. Stacking these one-dimensional profiles over time generated kymographs showing signal intensity as a function of time and position along the trench. Single-trench kymographs (Figure 5G-H) were generated by linking trenches across frames based on their x-center position and retaining tracks present for at least 20 frames. Ensemble kymographs (Figure S8C-E) were generated by averaging y-profiles across multiple trenches and/or positions at each frame. Time was aligned relative to AMP injection, with (t = 0) min corresponding to AMP addition. Fluorescence kymographs were plotted overlaid on matched phase-contrast kymographs.

To estimate the time when LL-37 fluorescence reached the bottom of the wide trench, the same single-trench tracking approach was used to quantify fluorescence intensity in a bottom ROI of three individual empty trenches using the script quantify_LL-37_fluor_empty_trench.py. Mean LL-37 fluorescence was quantified in a 40-px band at the bottom of selected tracked trenches (Figure S8F), background-corrected using a matched region outside the trench, aligned to AMP injection, and plotted in Figure S8G.

### Biofilm data processing and quantification

For the biofilm experiments, widefield fluorescence z-stacks were first deconvolved using Deconwolf ((Wernersson et al., 2024); https://deconwolf.fht.org/). Theoretical point-spread functions were generated with Deconwolf using an xy pixel size of 162.286 nm, a z-step size of 600 nm, 145 z-slices, NA = 0.95, refractive index = 1, and channel-specific emission wavelengths of 525 nm for GFP, 610 nm for SYTOX Orange, and 640 nm for EbbaBiolight 680, and are available at BioImage Archive (accession number: S-BIAD1823). Deconvolution was performed for 50 iterations with GPU acceleration. To quantify biofilm fluorescence, Deconwolf outputs were analysed using custom Python scripts (quantify_biofilm_stats.py). The scaling factors reported in the Deconwolf log files were used to rescale each deconvolved stack before quantification. For each position and time point, the first three z-planes were excluded to remove slices below or immediately adjacent to the glass surface, and the following 40 z-planes were retained for analysis. Total GFP and SYTOX Orange fluorescence signals were calculated by summing voxel intensities across the retained 3D volume and displayed in Figures 6C and S9D, respectively. For the 3D renderings shown in Figures 6A-B and S9C and Videos 10-17, deconvolved single-time-point stacks were first bundled into five-dimensional OME-TIFF, with dimensions corresponding to time, two fluorescence channels, and three spatial dimensions, using a custom Python script (export_ome-tiff_from_dw_zstack.py). During OME-TIFF generation, Deconwolf scaling factors were applied, voxel-size metadata was stored using the physical pixel sizes in x, y, and z, the first four z-planes were removed, and the central 50% of the field of view in x and y was retained. These OME-TIFF files were then converted to .ims files using ImarisFileConverter 11.0.1 (Oxford Instruments), imported into Imaris 10.2 (Oxford Instruments), rendered in 3D, and exported as TIFF image sequences with one file per frame. A separate Python script (export_3D_imaris_montage.py) was used to generate the video montages showing combined top and side views of the different fluorescence channels.

### Calculation of correlation coefficients

All Spearman’s correlations, used for SCF and SCF* calculations, were computed using the Python function *spearmanr* from *scipy.stats* library.

### Analysis software and hardware

Drift correction and cell linking using SuperSegger code (https://github.com/tlo-bot/omnisegger) were performed using MATLAB (www.mathworks.com). Repository for using the Omnipose deep neural network architecture (Cutler et al., 2022) was downloaded from https://github.com/kevinjohncutler/omnipose.git and installed in a local Miniconda environment.

Omnipose retraining, segmentation, 3D deconvolution with Deconwolf (Wernersson et al., 2024) (https://deconwolf.fht.org/), and computationally intensive tasks that involved operations on large arrays (background_subtraction.py; get_medial_axis.py) were accelerated by parallel processing on an NVIDIA A40 GPU utilizing CUDA (Compute Unified Device Architecture) to enhance performance. All custom-written code was developed using Python 3.9 (www.python.org). The following packages were employed in the analysis: *numpy* (Harris et al., 2020), *cupy* (Okuta et al., 2017), *pytorch* (Paszke et al., 2019), *pandas* (McKinney, 2010), *scipy* (Virtanen et al., 2020), *matplotlib* (Hunter, 2007), *seaborn* (Waskom, 2021), *scikit-image* (Walt et al., 2014). Interactive curation steps in the Omnipose retraining pipeline used the *keyboard* package for key-press input. For 3D rendering, OME-TIFF files were converted to .ims files using ImarisFileConverter 11.0.1 (Oxford Instruments) and rendered using Imaris 10.2 (Oxford Instruments) on a workstation at the Stanford University Cell Sciences Imaging Facility (CSIF). Final figures were assembled using Adobe Illustrator 2022. Videos 1-4 were exported as AVI files using Fiji/ImageJ (Schindelin et al., 2012) version 1.54p and converted to mp4 using HandBrake 1.9.2 (https://handbrake.fr), while for all the other videos a Python script that made use of *imageio_ffmpeg* was employed instead.

## DATA AVAILABILITY

The microscopy images generated and analysed in this study have been deposited in the BioImage Archive (BioStudies) under accession number S-BIAD1823.

## CODE AVAILABILITY

The analysis pipeline, custom-written functions, and trained Omnipose models used in this study are available in the Jacobs-Wagner laboratory GitHub repository at https://github.com/JacobsWagnerLab/published/tree/master/Fragasso_et_al_2025.

## ACKNOWLEDGEMENTS

We are grateful to Drs. Silvia Cañas-Duarte and Johan Paulsson for the gift of the periplasmic-mScarlet-I reporter construct and Dr. Johan Paulsson for also sharing the design for the 3D-printed holder used in this study. We thank Dr. Kristala Prather for the MG1655 (DE3) strain. We also thank Dr. Alexandros Papagiannakis and Kristian Sørensen for their gift of strain CJW7753 and peptoids, respectively. In addition, we are grateful to Dr. Fitnat Yildiz for sharing her expertise in biofilm biology, and to the members of the Jacobs-Wagner laboratory for fruitful discussions, support, and critical reading of the manuscript. We acknowledge the Stanford University Cell Sciences Imaging Facility (CSIF; RRID:SCR_017787) for access to the Imaris 10.2 workstation used for 3D rendering and visualization. This work was supported in part by a Director’s Pioneer Award (# 1DP1 OD029517) from The National Institute on Aging (to A.E.B.), funding from the SENS Foundation and Dr. James J. Truchard and the Truchard Foundation (to A.E.B.), and the Netherlands Organisation for Scientific Research (NWO Rubicon 2022-2 Science programme, 019.222EN.001804 to A.F.). A.E.B. acknowledges work at the Molecular Foundry supported by the Office of Science, Office of Basic Energy Sciences, of the U.S. Department of Energy under Contract No. DE-AC02-05CH11231. C.J.-W. is an investigator of the Howard Hughes Medical Institute.

## AUTHOR CONTRIBUTIONS

Conceptualization, A.F., A.E.B., and C.J.-W.; methodology, A.F., T.S., W.-H.L.; software, A.F.; formal analysis, A.F.; investigation, A.F.; resources, A.F., A.E.B., W.-H.L., and C.J.-W.; data curation, A.F.; writing – original draft, A.F. and C.J.-W.; writing – review and editing, A.F., T.S., A.E.B., C.J.-W.; visualization, A.F. and C.J.-W.; supervision, C.J.-W.; project administration, C.J.-W.; funding acquisition, A.F., A.E.B., and C.J.-W.

## SUPPLEMENTARY FIGURE LEGENDS

**Figure S1:**
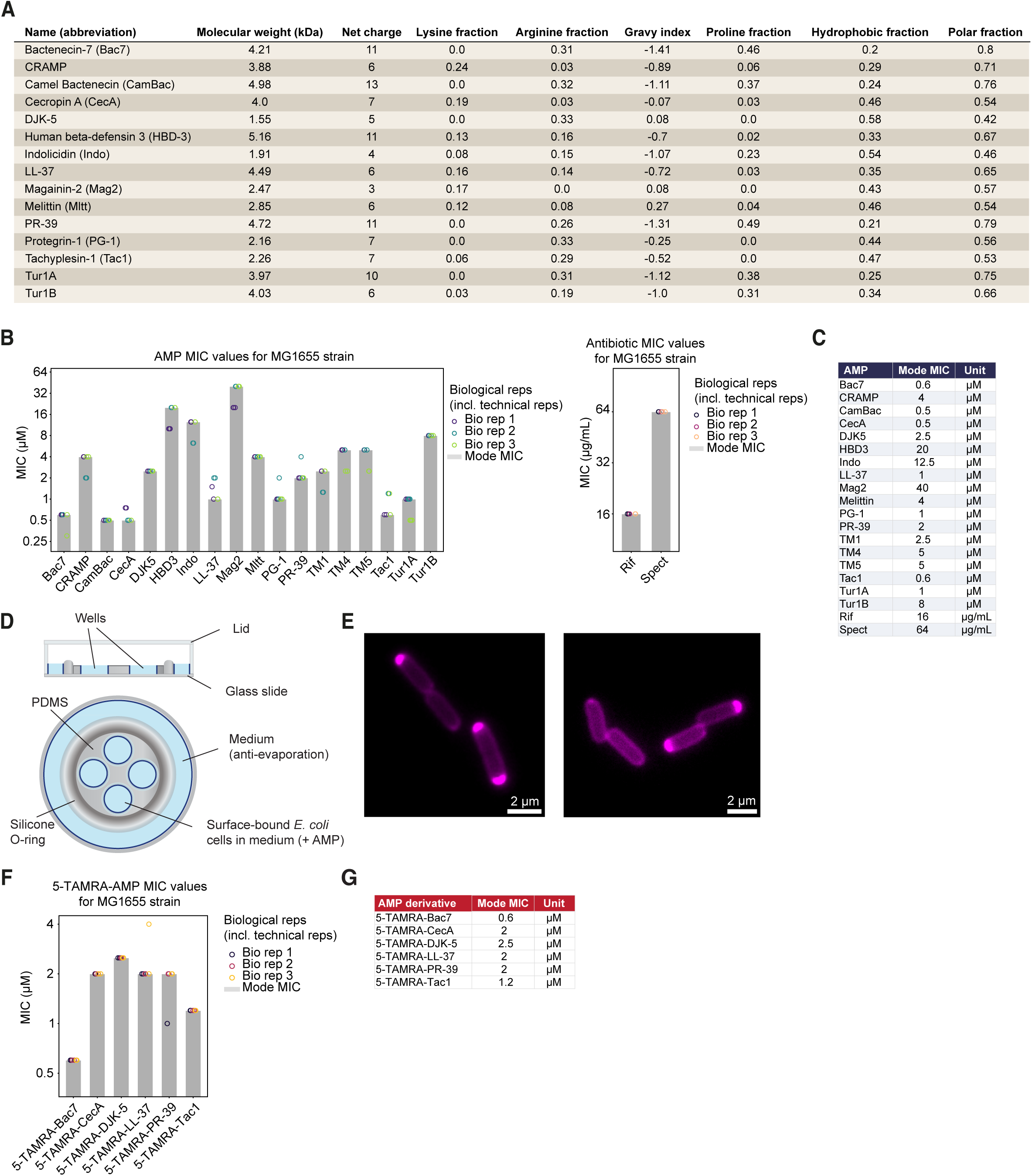
Features relevant to the experimental approach. **(A)** Data underlying scatterplots shown in Figure 1B for each AMP or PM (excluding peptoids TM1, TM4, TM5) listed in Figure 1A. **(B)** Bar plots showing the mode minimum inhibitory concentration (MIC) across biological replicates for each AMP. peptidomimetic, or antibiotic used in this study, measured in MG1655 *E. coli* cells grown in M9gluCAAT at 37°C for 20 h starting from an inoculum of ∼250 cells/mL. Grey bars indicate the mode MIC across three biological replicates (circles), featuring two technical replicates each. **(C)** Table summarizing the mode MIC values and units for all AMPs, peptidomimetics, and antibiotics shown in (B). **(D)** Schematic of the multi-well device used to image cells. The device contains four independent wells in which the growth and division of surface-immobilized *E. coli* cells can be tracked and AMPs can be added at defined final concentrations. **(E)** Representative fluorescence images of the mScarlet-I signal in CJW7845 cells, illustrating the common enrichment of the periplasmic fluorescent protein at cell poles. Scale bar, 2 µm. **(F)** Same as (A) but for the indicated 5-TAMRA–labeled AMP derivatives used to monitor intracellular accumulation. **(G)** Table summarizing the mode MIC values for the labeled AMP derivatives shown in (F).

**Figure S2:**
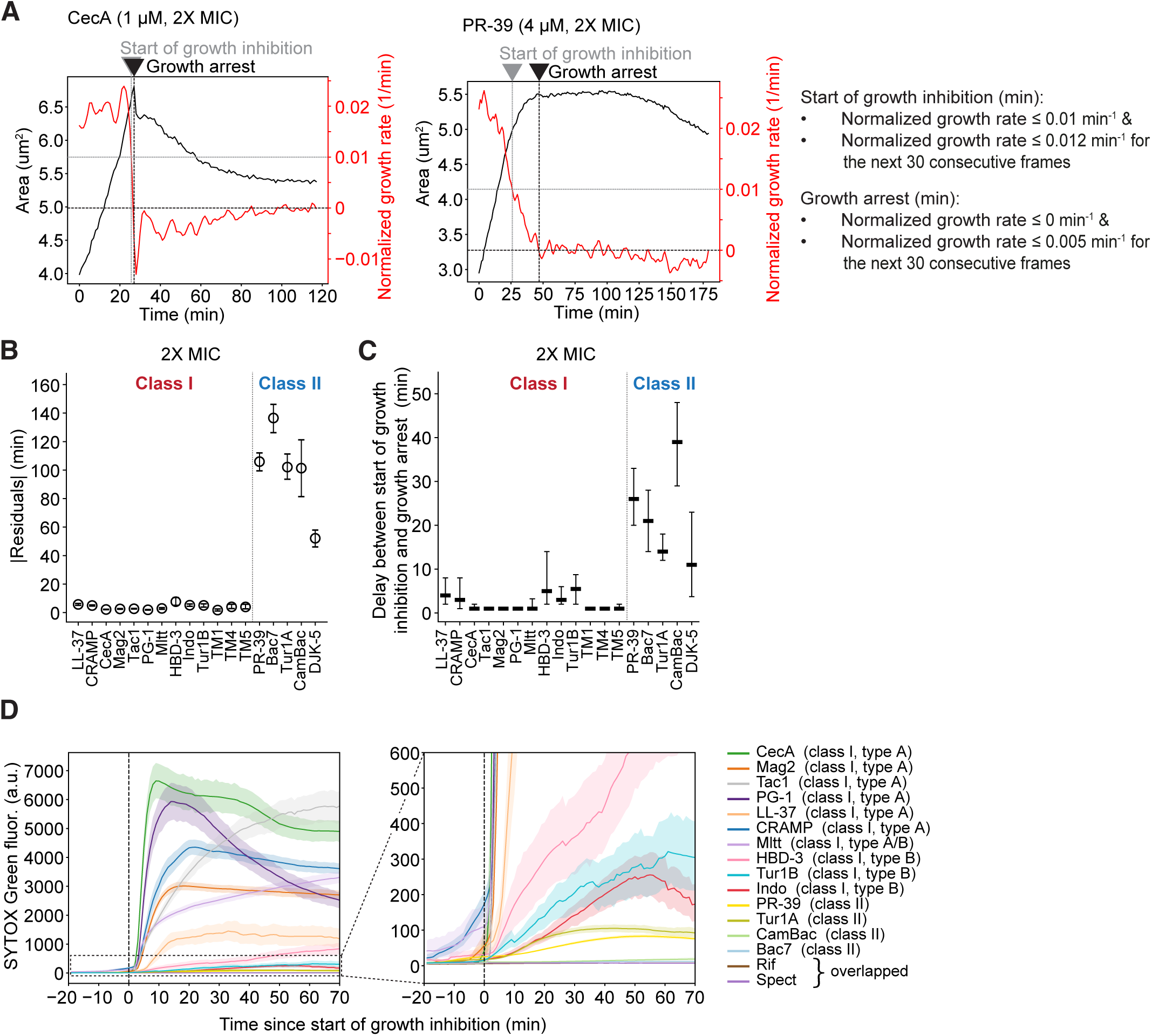
Determination of growth metrics and SYTOX Green uptake since start of growth inhibition. **(A)** Example trajectories of cell area (black) and normalized growth rate (red, as defined in Methods) for a cell treated with CecA (left) or PR-39 (right). Horizontal grey dotted and black dashed lines indicate hard thresholds used for detection of start of growth inhibition or growth arrest, respectively. Vertical grey dotted and black dashed lines indicate the identified times of start of growth inhibition and growth arrest, respectively, using the definitions written on the right. **(B)** Absolute residuals (mean ± 95% confidence interval (CI)) of times of growth arrest since AMP addition vs. times of IM permeabilization since AMP addition from the diagonal (*y = x*), for the indicated AMPs at 2X MIC (same dataset used in Figure 1F). **(C)** Time from the start of growth inhibition to growth arrest (points are medians; error bars represent interquartile range) for the indicated AMPs at 2X MIC (same dataset used in Figure 1F). **(D)** Plot showing SYTOX Green uptake *vs* time relative to initiation time of growth inhibition for different AMPs at concentrations equivalent to their 2X MICs (see Figure 1A). The left plot shows the full range of measured SYTOX Green values. The right plot shows a zoom-in to visualize the AMPs for which SYTOX Green uptake was lower. The total number of cells was 101, 288, 187, 196, 187, 205, 159, 158, 119, 201, 179, 176, 215, 110, 121, 140 for CecA, Tac1, PG-1, Mag2, LL-37, CRAMP, Mltt, HBD-3, Indo, Tur1B, PR-39, Bac7, Tur1A, CamBac, Rif, and Spect, respectively.

**Figure S3:**
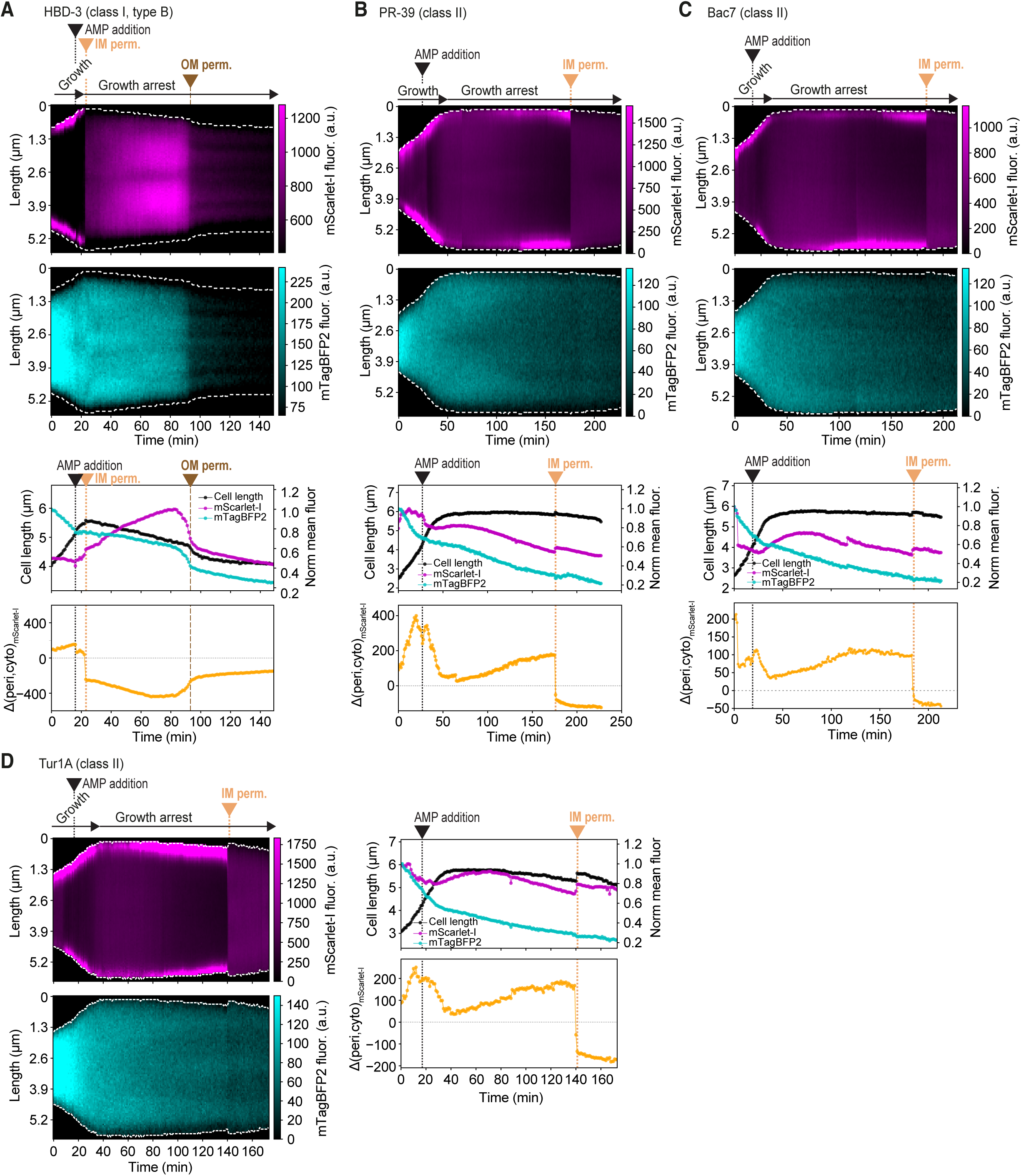
Example kymographs of mScarlet-I and mTagBFP2 signals for CJW7845 cells upon exposure to HBD-3, PR-3G, Bac7, and Tur1A. **(A)** *Top*: Example kymographs of mScarlet-I and mTagBFP2 signals for a representative CJW7845 cell exposed with HBD-3 (2X MIC) at the indicated time. The times of IM and OM permeabilizations are shown. *Bottom*: Plots showing the quantification of the cell length and fluorescence signal (top) and the difference Δ(peri,cyto)_mScarlet-I_ between the mScarlet-I signal in the cell periphery (periplasm) relative to the cell interior (cytoplasm) over time for cell shown above. **(B)** Same as (A) but for PR-39 (2X MIC). **(C)** Same as (A) but for Bac7 (2X MIC). **(D)** Same as (A) but for Tur1A (2X MIC).

**Figure S4:**
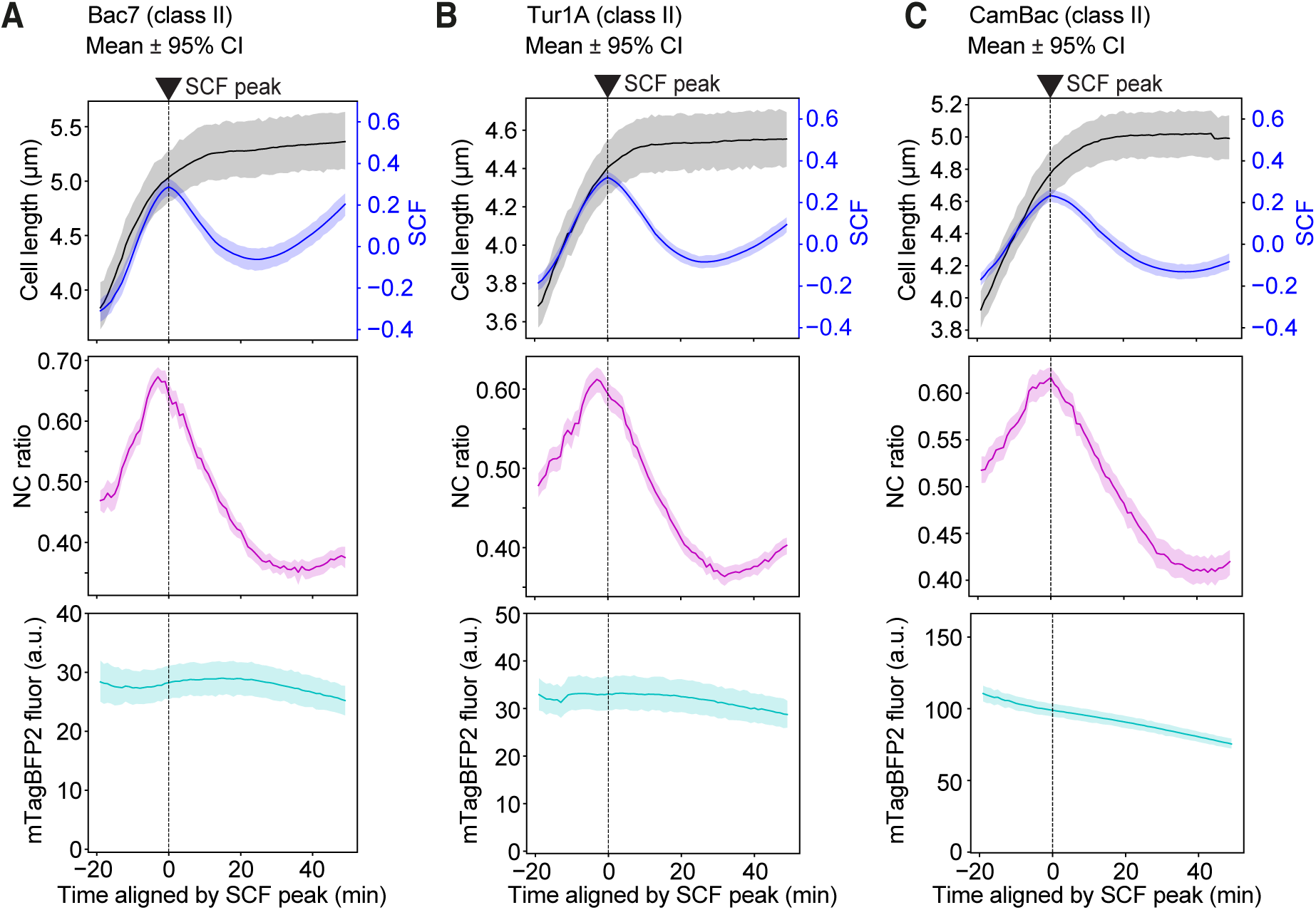
Quantification of the intracellular DNA and ribosome reorganization upon Bac7, Tur1A, and CamBac treatment. **(A)** Plots showing the quantification of the evolution of the mean ± 95% confidence interval (CI) of the cell length, signal correlation factor (SCF), nucleocytoplasmic (NC) ratio, and mTagBFP2 fluorescence (fluor.) across 183 cells exposed to Bac7 at 2X MIC. Times were aligned based on the SCF peak at the single-cell level. The black dashed line indicates the SCF peak (time zero). **(B)** Same as (A) but for Tur1A. The number of cells is 131. **(C)** Same as (A) and (B) but for CamBac. The number of cells is 168.

**Figure S5.**
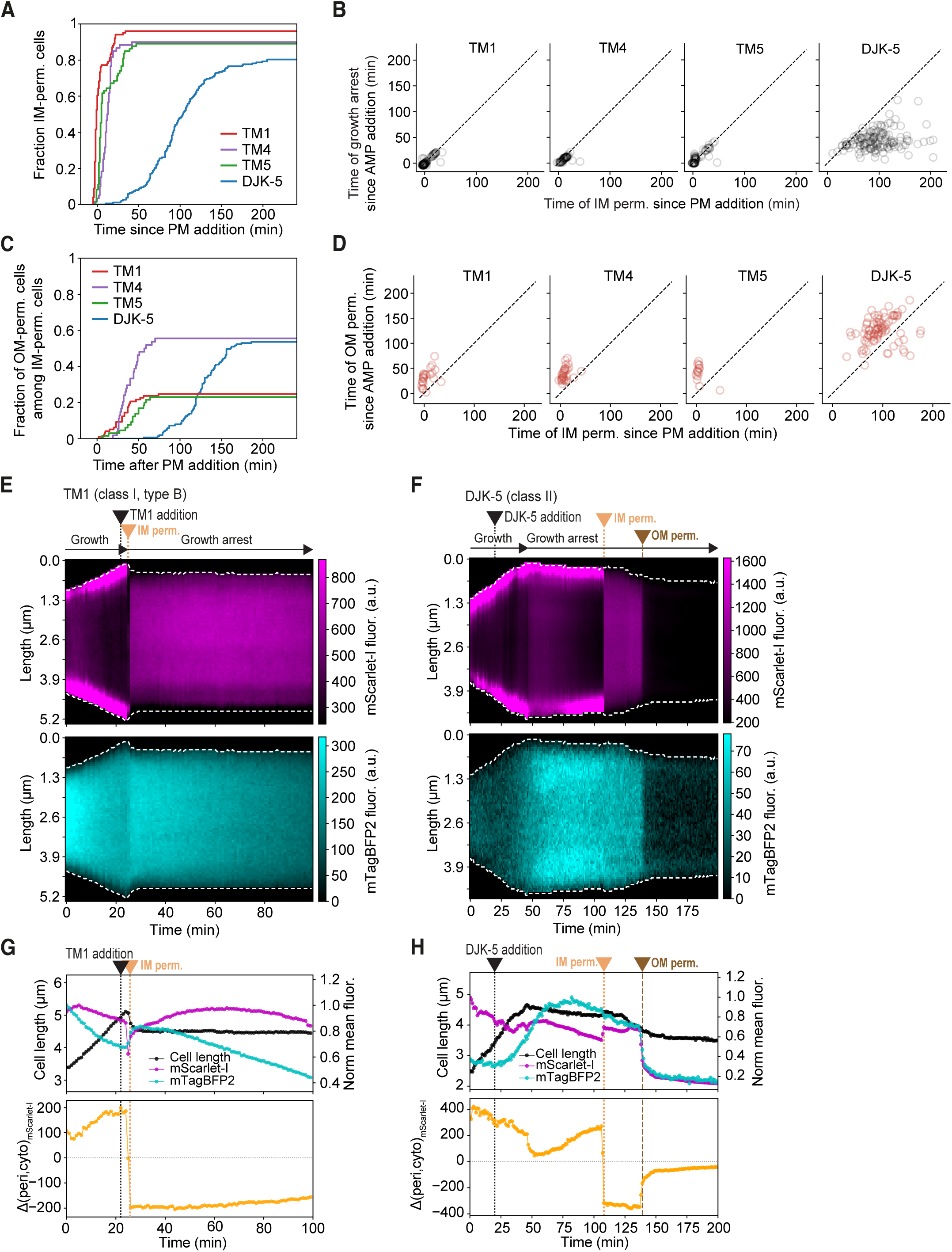
Phenotype-based classification applies to synthetic AMP mimics. For all panels, the concentration of peptidomimetics (PM) used was 2X MIC (see Figure 1A). **(A)** Plot showing the cumulative distribution of the fraction of IM-permeabilized cells treated with peptidomimetics (PMs) TM1, TM4, TM5, and DJK-5. The total number of IM-permeabilized cells is 97, 54, 65, and 151 for TM1, TM4, TM5, and DJK-5, respectively**. (B)** Scatterplots showing the time of growth arrest since PM addition *vs.* that of inner membrane (IM) permeabilization since PM addition for cells treated with TM1, TM4, TM5, and DJK-5. The total number of cells is the same as in (A). Opacity of each data point was set to 15% to aid visualization of the data density. The dashed line indicates *y = x***. (C)** Plot showing the cumulative distribution of the fraction of OM-permeabilized cells among the IM-permeabilized population during treatment with TM1, TM4, TM5, and DJK-5. The fraction of cells was 24/97 for TM1, 30/54 for TM4, 15/65 for TM5, and 81/151 for DJK-5. **(D)** Scatterplots showing the time of OM permeabilization since PM addition *vs.* that of IM permeabilization since PM addition for cells treated with TM1, TM4, TM5, and DJK-5, at 2X MIC. The total number of cells is the same as in (C). Opacity of each data point was set to 25% to aid visualization of the data density. The dashed line indicates *y = x***. (E)** Example kymographs showing the evolution of the mScarlet-I and mTagBFP2 signals for a representative CJW7845 cell exposed with TM1 at. **(F)** Same as (E) but for DJK-5). **(G)** Plots of the cell length, normalized mean fluorescence signals of mScarlet-I and mTagBFP2, and difference Δ(peri,cyto)_mScarlet-I_ between the mScarlet-I fluorescence signal in the cell periphery (periplasm) relative to the cell interior (cytoplasm) over time, for the cell represented in (E), treated with TM1. **(H)** Same as (G) but for the cell in (F) treated with DJK-5.

**Figure S6:**
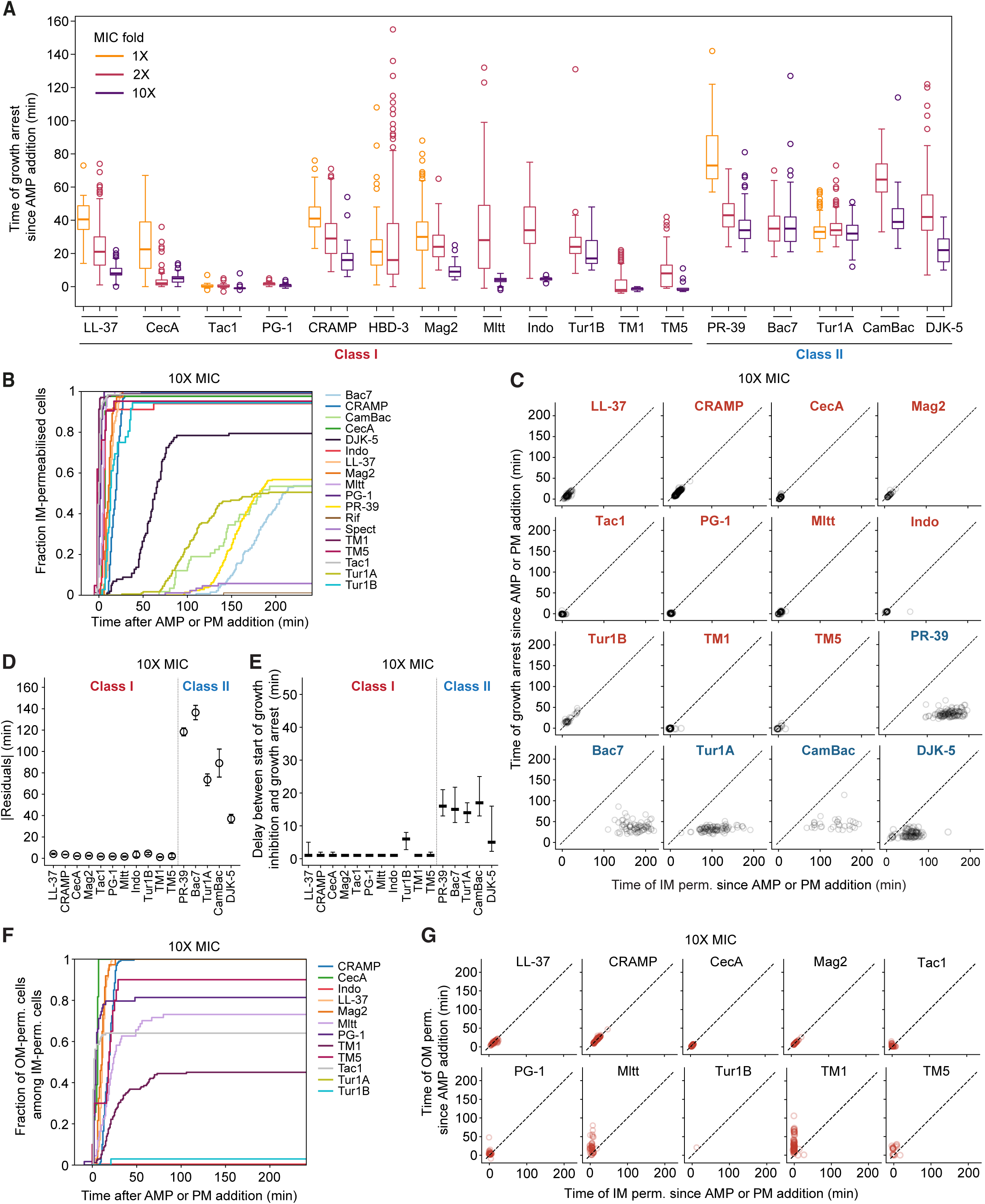
Quantification of the time delay between AMP addition and growth arrest for all tested AMPs across different peptide concentrations relative to their MICs. **(A)** Box plots showing the distribution of times of growth arrest since AMP or peptidomimetic (PM) addition for CJW7845 cells treated at 1X (orange), 2X (pink), and 10X (purple) MIC. AMPs or PMs are separated into class I (left) and class II (right). Boxes show the interquartile range; whiskers extend to 1.5 times the interquartile range; circles indicate outliers. The total number of cells per condition is as follows (1X/2X/10X MIC): CecA (52/100/84), LL-37 (14/167/90), CRAMP (58/194/201), Tac1 (73/274/130), Mag2 (300/181/37), HBD-3 (84/99/–), PG-1 (–/163/59), Mltt (–/156/67), Indo (–/119/34), Tur1B (–/54/36), TM1 (–/101/173), TM4 (–/60/–), TM5 (–/73/21), PR-39 (112/158/213), Bac7 (–/155/134), Tur1A (289/192/180), CamBac (–/98/58), and DJK-5 (–/188/102). **(B)** Cumulative distributions of the fraction of IM-permeabilized CJW7845 cells for all AMPs at 10X MIC, as a function of time since AMP addition. The total number of IM-permeabilized cells is 90, 82, 200, 128, 37, 67, 59, 32, 34, 171, 20, 121, 72, 91, 31, 81, 1, and 5 for LL-37, CecA, CRAMP, Tac1, Mag2, Mltt, PG-1, Indo, Tur1B, TM1, TM5, PR-39, Bac7, Tur1A, CamBac, DJK-5, Rif, and Spect, respectively. **(C)** Scatterplots showing the time of growth arrest since AMP or PM addition vs. that of inner membrane (IM) permeabilization since AMP or PM addition for all indicated AMPs and PMs at 10X MIC. The total number of cells is the same as in (B). Opacity of each data point was set to 15% to aid visualization of the data density. The dashed line indicates y = x. **(D)** Absolute residuals of times of growth arrest since AMP addition vs. times of IM permeabilization since AMP or PM addition from the diagonal (*y = x*), for the indicated AMPs and PMs at 10X MIC (same dataset used in (B)). **(E)** Delay between the start of growth inhibition and the growth arrest (points are medians; error bars represent interquartile range) for the indicated AMPs at 10X MIC (same dataset used in (B)). **(F)** Plot showing the cumulative distribution of the fraction of OM-permeabilized cells among the IM-permeabilized population during treatment with the indicated AMPs and PMs at 10X MIC. The total number of OM-permeabilized cells (out of all IM-permeabilized cells) was 90/90 for LL-37, 82/82 for CecA, 200/200 for CRAMP, 82/128 for Tac1, 37/37 for Mag2, 49/67 for Mltt, 48/59 for PG-1, 1/34 for Tur1B, 77/171 for TM1, and 18/20 for TM5. **(G)** Scatterplots showing the time of outer membrane (OM) permeabilization since AMP or PM addition vs. that of inner membrane (IM) permeabilization since AMP or PM addition for cells treated with the indicated AMPs or PMs, at 10X MIC. The total number of cells is the same as in (F). Opacity of each data point was set to 25% to aid visualization of the data density. The dashed line indicates *y = x*.

**Figure S7:**
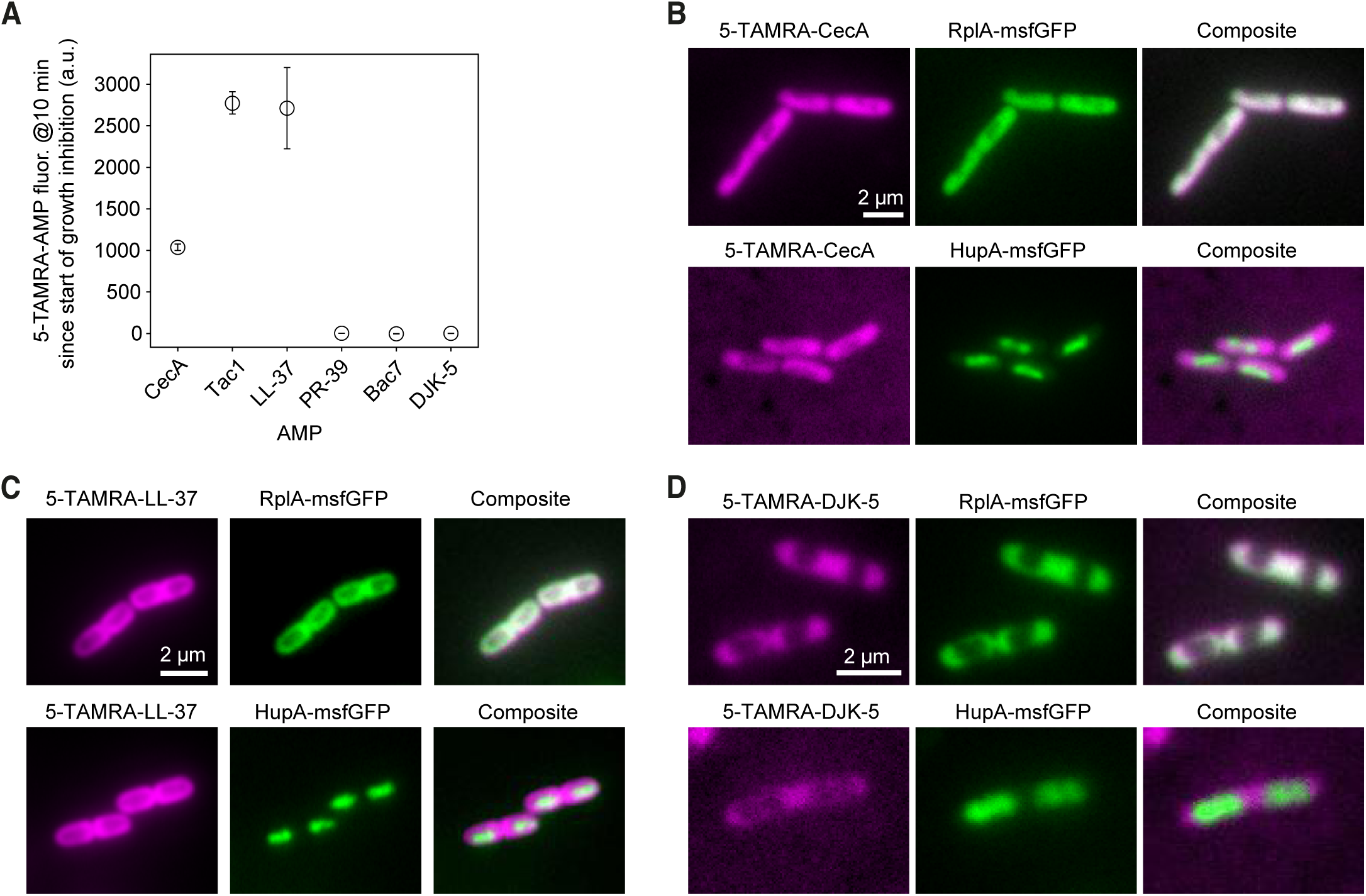
Intracellular enrichment and spatial localization of fluorescently labeled AMPs. **(A)** Intracellular fluorescence intensity (mean± 95 % CI) of 5-TAMRA conjugates to the indicated AMPs 10 min after their addition for MG1655 strain (same data as in Figure 4A) present at 50 nM together with the corresponding unlabeled AMPs (2X MIC) in MG1655 cells. Time is relative to the start time of growth inhibition. The total number of averaged cells is 138, 116, 101, 125, 254, 180, for CecA, Tac1, LL-37, PR-39, Bac7, and DJK-5, respectively. **(B)** Representative images of the indicated fluorescence signal in CJW7020 cells (expressing RplA-msfGFP, top) or CJW7859 (expressing HupA-msfGFP, bottom) exposed to a mixture of CecA (2X MIC) and 5-TAMRA-CecA (50 nM, 5% of the unlabeled version) for 10 min or 20 min, respectively. **(C)** Same as (B) except that cells were exposed to a mixture of LL-37 (2X MIC) and 100 nM (5% of the unlabeled version) of 5-TAMRA-LL-37 for 50 min. **(D)** Same as (B) except that cells were exposed to a mixture of DJK-5 (2X MIC) and 5-TAMRA-DJK-5 (250 nM, 5% of the unlabeled version) for 5 h and 40 min (for CJW7020 cell, top) or 3 h and 10 min (for CJW7859 cells, bottom).

**Figure S8:**
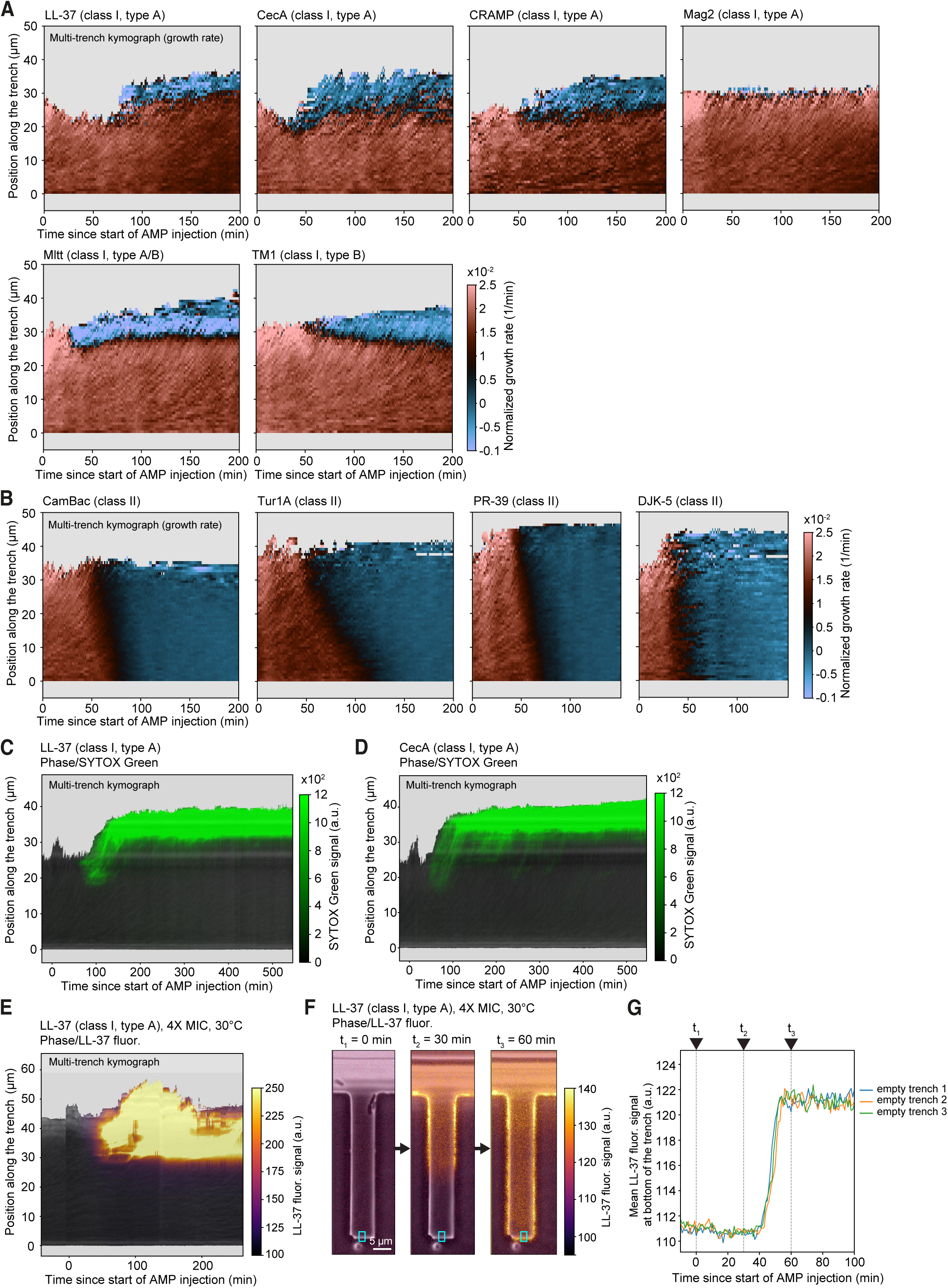
Ensemble kymographs of growth rate, SYTOX Green, and LL-37 fluorescence in wide trenches. **(A)** Ensemble kymographs of the normalized growth rate distribution along wide trenches over time, computed by averaging across multiple trenches for CJW7753 cells treated with the indicated class I AMPs or PMs at 2X MIC: The number of trenches is 18, 24, 12, 23, 18, and 17, and the number of cells is 5012, 5081, 2590, 6707, 6317, and 6266 for LL-37, CecA, CRAMP, Mag2, Mltt, and TM1, respectively. **(B)** Same as (A) but for CJW7753 cells treated with the indicated class II peptides at 2X MIC. The number of trenches is 32, 28, 27, and 28, and the number of cells is 4189, 4210, 4312, and 2989 for CamBac, Tur1A, PR-39, and DJK-5, respectively. **(C)** Multi-trench kymograph showing the ensemble average of phase-contrast (grey) and SYTOX Green fluorescence (green) along wide trenches over time for CJW7845 cells treated with LL-37 (class I type A) at 2X MIC. The kymograph was computed by averaging across 24 trenches. SYTOX Green signal represents cells with permeabilized membranes. **(D)** Same as (C) but for CJW7845 cells treated with CecA (class I type A), computed from 24 trenches. **(E)** Multi-trench kymograph showing phase-contrast and LL-37 fluorescence signal along wide trenches over time for CJW7859 cells (expressing HupA-msfGFP) treated with LL-37 (4X MIC) at 30°C, averaged across 18 trenches. The fluorescence reports on LL-37 absorption by cells. **(F)** Representative fluorescence images of a single empty trench at three selected timepoints (t₀, t₁, t₂) illustrating how LL-37 can diffuse all the way to the bottom of the empty trench within 60 min from the start of AMP injection. Cyan colored box represents the ROI where LL-37 signal intensity was computed over time and shown in (G). **(G)** Plot showing LL-37 fluorescence intensity as a function of time since AMP injection, measured at the bottom of three empty trenches by averaging pixel intensities within a defined ROI (as illustrated in (F)).

**Figure S9:**
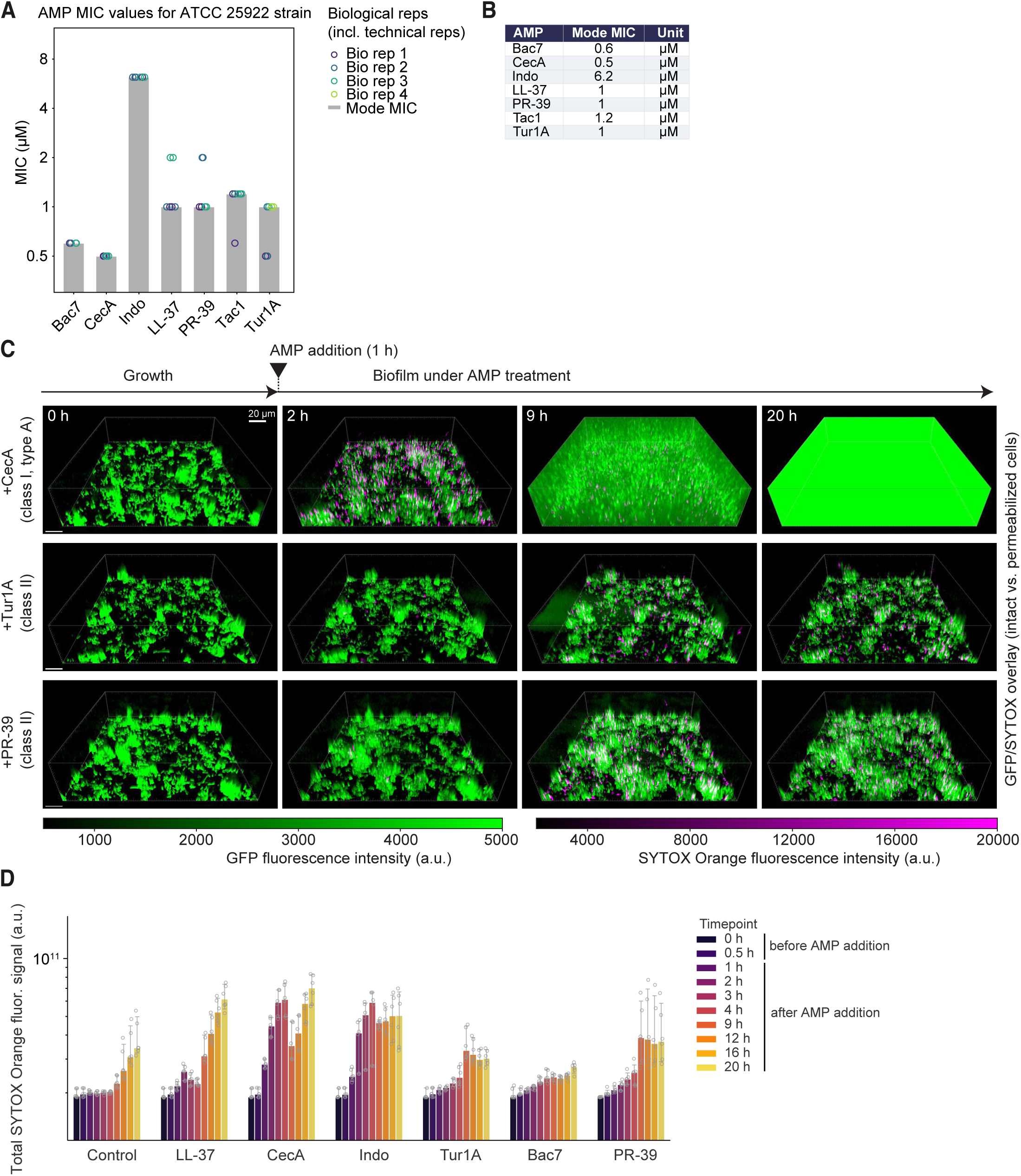
Biofilm growth under AMP treatment. **(A)** Bar plot showing the mode MIC across biological replicates for the indicated AMPs measured against *E. coli* ATCC 25922 in M9gluCAAT at 37°C for 20 h. Grey bars indicate the mode MIC across four biological replicates (circles). **(B)** Table summarizing the mode MIC values (in µM) for each AMP shown in (A). **(C)** Selected timepoints of biofilms treated with (or without) AMPs, showing overlaid GFP (cells) and SYTOX Orange (permeabilized cells) signals. AMPs were added at 2X MIC ∼1 h after the start of image acquisition. **(D)** Bar plot showing the quantification of the total SYTOX Orange signal within the imaged 3D volume over time for the indicated conditions. Fluorescence signals in (C) were rendered and exported from Imaris with gamma set to 2 for display only.

**Figure S10.**
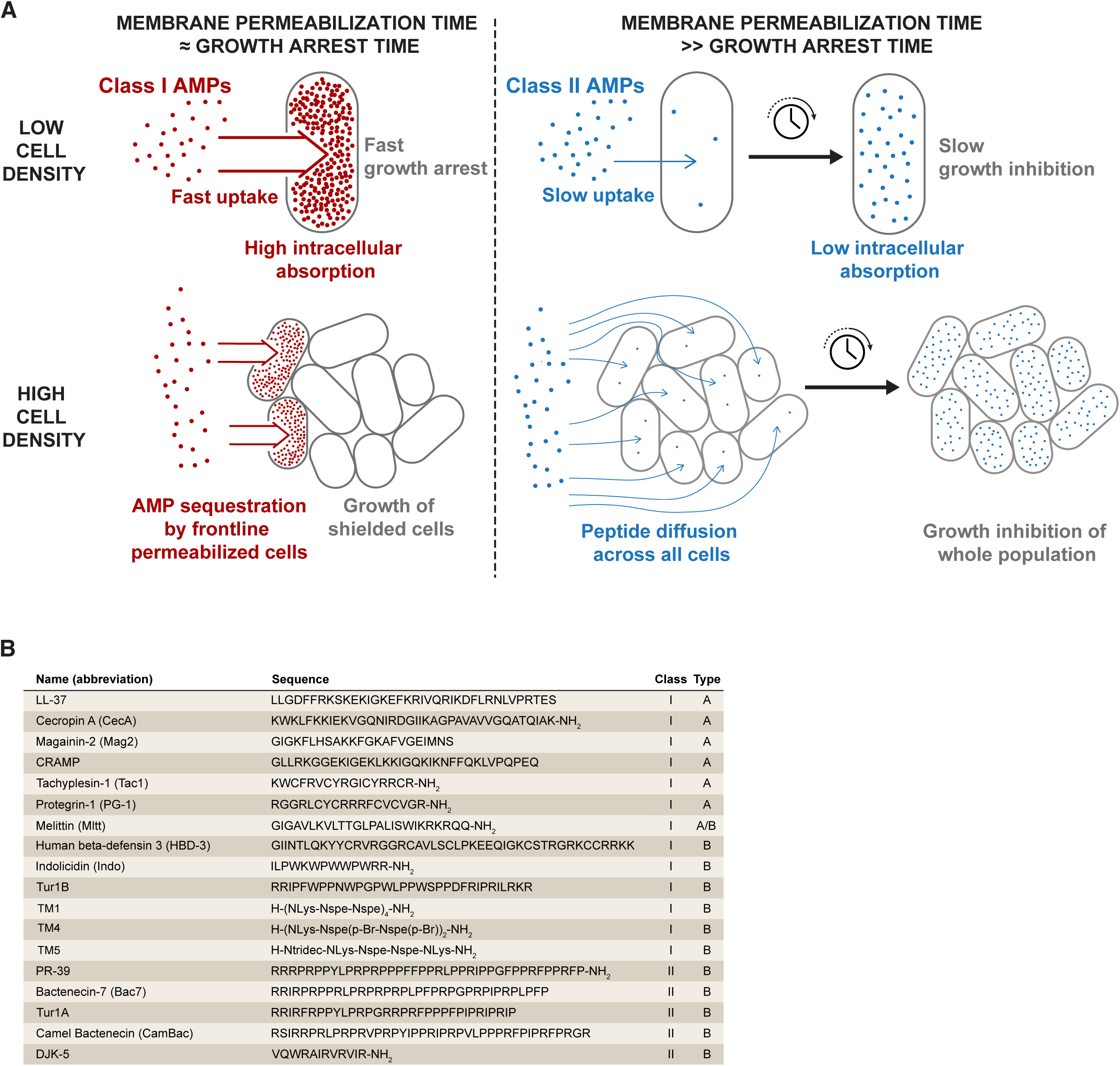
Summary of the functional trade-offs and AMP classification. **(A)** Class I AMPs (red) couple fast membrane permeabilization with rapid growth arrest. At low cell density, fast peptide uptake leads to high intracellular accumulation. At high cell density, frontline permeabilized cells rapidly sequester the bulk of available peptide, shielding the cells behind them from exposure. Class II AMPs (blue) permeabilize membranes more slowly than they inhibit growth, resulting in slow, gradual intracellular accumulation and low absorption per cell. At high cell density, this slow uptake allows peptides to diffuse across the population before any individual cell sequesters a significant fraction, ultimately inhibiting growth of the whole population. **(B)** Sequences and phenotypic classification of the AMPs and PMs used in this study. Class I AMPs and PMs cause growth arrest predominantly associated with inner membrane (IM) permeabilization, whereas class II compounds inhibit growth without substantial IM permeabilization. Within class I, type A AMPs predominantly permeabilize the outer membrane (OM) within 10 min of the IM, whereas type B AMPs and PMs permeabilize the IM well before (>10 min) the OM. All class II peptides are found to be type B. Melittin (Mltt) displays intermediate behavior and is classified as type A/B.

## VIDEO LEGENDS

**Video 1: Timelapse sequence showing inner and outer membrane permeabilization to proteins upon CecA treatment.** Montage video showing phase contrast, mScarlet-I fluorescence, mTagBFP2 fluorescence, and SYTOX Green fluorescence in the top row of a representative CJW7845 cell under treatment with 2X MIC CecA. SYTOX Green signal is plotted using a low and a high dynamic range (DR) of intensity values to enable visualization at both low- and high-intensity values, respectively. Time stamp shows h:min. The scale bar is 1 μm. The bottom row shows extracted cell parameters (cell length and mean intensities for each fluorescent probe) over time for the same cell. The red dashed line indicates the time of CecA addition. The black dotted line indicates the time at which the cell reaches the maximum length prior to cell shrinkage.

**Video 2: Timelapse sequence showing inner membrane permeabilization to proteins upon Indo treatment.** Montage video showing phase contrast, mScarlet-I fluorescence, mTagBFP2 fluorescence, and SYTOX Green fluorescence in the top row of a representative CJW7845 cell under treatment with 2X MIC Indo. SYTOX Green signal is displayed both with adjusted high brightness (high br.) and low brightness (low br.) to enable visualization at both low and high intensity values, respectively. The time stamp shows h:min. The scale bar is 1 μm. The bottom row shows extracted cell parameters (cell length and mean intensities for each fluorescent probe) over time for the same cell. The red dashed line indicates the time of Indo addition. The black dotted line indicates the time at which the cell reaches the maximum length prior to cell shrinkage.

**Video 3: Timelapse sequence showing intracellular DNA and ribosome reorganization upon CecA treatment.** Montage video showing phase contrast, RplA-GFP, HupA-mCherry, and mTagBFP2 fluorescence in the top row of a representative CJW7753 cell under treatment with 2X MIC CecA. The time stamp shows h:min. The scale bar is 1 μm. The bottom row shows plots of cell length, signal correlation factor (SCF), nucleocytoplasmic (NC) ratio, and mean mTagBFP2 intensity. The red dashed line indicates the time of CecA addition. The black dotted line indicates the time at which the cell reaches the maximum length prior to cell shrinkage.

**Video 4: Timelapse sequence showing DNA and ribosome reorganization upon PR-3G treatment.** Montage video showing phase contrast, RplA-GFP, HupA-mCherry, and mTagBFP2 fluorescence in the top row of a representative CJW7753 cell under treatment with 2X MIC PR-39. The time stamp shows h:min. The scale bar is 1 μm. The bottom row shows plots of cell length, signal correlation factor (SCF), nucleocytoplasmic (NC) ratio, and mean mTagBFP2 intensity. The red line indicates the time of PR-39 addition.

**Video 5: Timelapse sequence and single-cell tracking showing the poor killing activity of Tac1 against a dense population of *E. coli* cells in a wide trench.** Montage video showing phase contrast, RplA-GFP, and HupA-mCherry fluorescence of CJW7753 cells over time, together with their positions (circles), and their trajectories along the vertical axis in a representative wide trench under treatment with Tac1 at concentration that corresponds to 2X MIC at low cell density (see Figure 1A). The color scale represents the measured growth rate for each single cell (brown for growing, blue for growth inhibited). Time stamp shows h:min. The scale bar is 5 μm. The yellow label appearance indicates the time at which medium containing Tac1 began flowing into the microfluidic system.

Video 6: Timelapse sequence and single-cell tracking showing the ability of PR-39 to inhibit the growth of a dense population of *E. coli* cells in a wide trench. Montage video showing phase contrast, RplA-GFP, and HupA-mCherry fluorescence of CJW7753 cells over time, together with their positions (circles) and trajectories along the vertical axis (continuous lines) in a representative wide trench under treatment with PR-39 at a concentration corresponding to 2X MIC at low cell density (see Figure 1A). The color scale represents the measured growth rate for each single cell (brown for growing, blue for growth inhibited). Time stamp shows h:min. The scale bar is 5 μm. The yellow label appearance indicates the time at which medium containing PR-39 began flowing into the microfluidic system.

**Video 7: Timelapse sequence showing the localized permeabilization and overall poor killing activity of LL-37 against a dense E. coli cell population in wide trenches.** Montage video showing phase contrast overlaid with SYTOX Green fluorescence of CJW7845 cells in six representative wide trenches under treatment with LL-37 at a concentration corresponding to 2X MIC at low cell density (see Figure 1A). SYTOX Green signal marks cells with compromised membranes. Time stamp shows h:min. The scale bar is 5 µm. The yellow label appearance indicates the time at which medium containing LL-37 and SYTOX Green (5 nM) began flowing into the microfluidic system.

**Video 8: Timelapse sequence showing the localized permeabilization and overall poor killing activity of CecA against a dense E. coli cell population in wide trenches.** Montage video showing phase contrast overlaid with SYTOX Green fluorescence of CJW7845 cells in six representative wide trenches under treatment with CecA at a concentration corresponding to 2X MIC at low cell density (see Figure 1A). SYTOX Green signal marks cells with compromised membranes. Time stamp shows h:min. The scale bar is 5 µm. The yellow label appearance indicates the time at which medium containing CecA and SYTOX Green (5 nM) began flowing into the microfluidic system.

**Video G: Timelapse sequence showing the high cell absorption and poor killing activity of LL-37 at 4X MIC in a wide trench containing a dense *E. coli* cell population growing at 30°C.** Montage video showing phase contrast, LL-37 fluorescence, and their overlay for CJW7859 cells in a representative wide trench under treatment with LL-37 at a concentration corresponding to 4X MIC at low cell density (see Figure 1A) at 30°C. The empty trench on the left is shown to showcase the diffusion of LL-37 across the wide trench. Time stamp shows h:min. The scale bar is 5 µm. The yellow label appearance indicates the time at which medium containing LL-37 began flowing into the microfluidic system.

**Video 10: Timelapse sequence showing the growth, matrix production, and evolution of *E. coli* ATCC 25G22GFP biofilms in the absence of AMP (control).** Montage video showing 3D side views (left) with GFP fluorescence (green, cells) and EbbaBiolight 680 fluorescence (warm colors, extracellular matrix), alongside top views (right), for the same biofilm well. Biofilms were grown at 37°C in M9gluCAAT medium. Frames were acquired every 30 min. Time stamp shows hh:mm. For visualization, fluorescence channels were displayed and exported from Imaris with gamma set to 2; this adjustment was applied only for display.

**Video 11: Timelapse sequence showing the growth and evolution of *E. coli* ATCC 25G22GFP biofilms in the absence of AMP (control).** Montage video showing GFP fluorescence (green, cells) overlaid with SYTOX Orange fluorescence (magenta, permeabilized cells) over time at 37°C in M9gluCAAT medium. Frames were acquired every 30 min. Time stamp shows hh:mm. For visualization, fluorescence channels were displayed and exported from Imaris with gamma set to 2; this adjustment was applied only for display.

**Video 12: Timelapse sequence showing the growth and evolution of *E. coli* ATCC 25G22GFP biofilms treated with LL-37 at 2X MIC.** Montage video showing GFP fluorescence (green, cells) overlaid with SYTOX Orange fluorescence (magenta, permeabilized cells) over time at 37°C in M9gluCAAT medium. The appearance of the yellow label between the second and third frames indicates the time of LL-37 addition. Each frame was acquired every 30 min. Time stamp shows hh:mm. For visualization, fluorescence channels were displayed and exported from Imaris with gamma set to 2; this adjustment was applied only for display.

**Video 13: Timelapse sequence showing the growth and evolution of *E. coli* ATCC 25G22GFP biofilms treated with CecA at 2X MIC.** Montage video showing GFP fluorescence (green, cells) overlaid with SYTOX Orange fluorescence (magenta, permeabilized cells) over time at 37°C in M9gluCAAT medium. The appearance of the yellow label between the second and third frames indicates the time of CecA addition. Each frame was acquired every 30 min. Time stamp shows hh:mm. For visualization, fluorescence channels were displayed and exported from Imaris with gamma set to 2; this adjustment was applied only for display.

**Video 14: Timelapse sequence showing the growth and evolution of *E. coli* ATCC 25G22GFP biofilms treated with Indo at 2X MIC.** Montage video showing GFP fluorescence (green, cells) overlaid with SYTOX Orange fluorescence (magenta, permeabilized cells) over time at 37°C in M9gluCAAT medium. The appearance of the yellow label between the second and third frames indicates the time of Indo addition. Each frame was acquired every 30 min. Time stamp shows hh:mm. For visualization, fluorescence channels were displayed and exported from Imaris with gamma set to 2; this adjustment was applied only for display.

**Video 15: Timelapse sequence showing the growth inhibition of an *E. coli* ATCC 25G22GFP biofilms treated with Tur1A at 2X MIC.** Montage video showing GFP fluorescence (green, cells) overlaid with SYTOX Orange fluorescence (magenta, permeabilized cells) over time at 37°C in M9gluCAAT medium. The appearance of the yellow label between the second and third frames indicates the time of Tur1A addition. Each frame was acquired every 30 min. Time stamp shows hh:mm. For visualization, fluorescence channels were displayed and exported from Imaris with gamma set to 2; this adjustment was applied only for display.

**Video 16: Timelapse sequence showing the growth inhibition of *E. coli* ATCC 25G22GFP biofilms treated with Bac7 at 2X MIC.** Montage video showing GFP fluorescence (green, cells) overlaid with SYTOX Orange fluorescence (magenta, permeabilized cells) over time at 37°C in M9gluCAAT medium. The appearance of the yellow label between the second and third frames indicates the time of Bac7 addition. Each frame was acquired every 30 min. Time stamp shows hh:mm. For visualization, fluorescence channels were displayed and exported from Imaris with gamma set to 2; this adjustment was applied only for display.

**Video 17: Timelapse sequence showing the growth inhibition of *E. coli* ATCC 25G22GFP biofilms treated with PR-39 at 2X MIC.** Montage video showing GFP fluorescence (green, cells) overlaid with SYTOX Orange fluorescence (magenta, permeabilized cells) over time at 37°C in M9gluCAAT medium. The appearance of the yellow label between the second and third frames indicates the time of PR-39 addition. Each frame was acquired every 30 min. Time stamp shows hh:mm. For visualization, fluorescence channels were displayed and exported from Imaris with gamma set to 2; this adjustment was applied only for display.

## REFERENCES

Bakshi, S., Siryaporn, A., Goulian, M., & Weisshaar, J. C. (2012). Superresolution imaging of ribosomes and RNA polymerase in live *Escherichia coli* cells. Molecular Microbiology, 85(1), 21–38. 10.1111/j.1365-2958.2012.08081.x

Balleza, E., Kim, J. M., & Cluzel, P. (2018). Systematic characterization of maturation time of fluorescent proteins in living cells. Nature Methods, 15(1), 47–51. 10.1038/nmeth.4509

Benn, G., Pyne, A. L. B., Ryadnov, M. G., & Hoogenboom, B. W. (2019). Imaging live bacteria at the nanoscale: Comparison of immobilisation strategies. Analyst, 144(23), 6944–6952. 10.1039/C9AN01185D

Bevis, B. J., & Glick, B. S. (2002). Rapidly maturing variants of the Discosoma red fluorescent protein (DsRed). Nature Biotechnology, 20(1), 83–87. 10.1038/nbt0102-83

Boeynaems, S., Ma, X. R., Yeong, V., Ginell, G. M., Chen, J.-H., Blum, J. A., Nakayama, L., Sanyal, A., Briner, A., Van Haver, D., Pauwels, J., Ekman, A., Schmidt, H. B., Sundararajan, K., Porta, L., Lasker, K., Larabell, C., Hayashi, M. A. F., Kundaje, A.,…Gitler, A. D. (2023). *Aberrant phase separation is a common killing strategy of positively charged peptides in biology and human disease* [Preprint]. Cell Biology. 10.1101/2023.03.09.531820

Boman, H. G., Agerberth, B., & Boman, A. (1993). Mechanisms of action on Escherichia coli of cecropin P1 and PR-39, two antibacterial peptides from pig intestine. *Infection and Immunity*, 61(7), 2978–2984.

Cañas-Duarte, S. J. (2021). Understanding rare events in Escherichia coli. https://nrs.harvard.edu/URN-3:HUL.INSTREPOS:37368226

Cardoso, M. H., Meneguetti, B. T., Costa, B. O., Buccini, D. F., Oshiro, K. G. N., Preza, S. L. E., Carvalho, C. M. E., Migliolo, L., & Franco, O. L. (2019). Non-lytic antibacterial peptides that translocate through bacterial membranes to act on intracellular targets. International Journal of Molecular Sciences, 20(19). 10.3390/ijms20194877

Cherepanov, P. P., & Wackernagel, W. (1995). Gene disruption in *Escherichia coli*: TcR and KmR cassettes with the option of Flp-catalyzed excision of the antibiotic-resistance determinant. Gene, 158(1), 9–14. 10.1016/0378-1119(95)00193-A

Choi, H., Rangarajan, N., & Weisshaar, J. C. (2016). Lights, Camera, Action! Antimicrobial Peptide Mechanisms Imaged in Space and Time. Trends in Microbiology, 24(2), 111–122. 10.1016/j.tim.2015.11.004

Chongsiriwatana, N. P., Lin, J. S., Kapoor, R., Wetzler, M., Rea, J. A. C., Didwania, M. K., Contag, C. H., & Barron, A. E. (2017). Intracellular biomass flocculation as a key mechanism of rapid bacterial killing by cationic, amphipathic antimicrobial peptides and peptoids. Scientific Reports, 7(1), 1–15. 10.1038/s41598-017-16180-0

Chongsiriwatana, N. P., Patch, J. A., Czyzewski, A. M., Dohm, M. T., Ivankin, A., Gidalevitz, D., Zuckermann, R. N., & Barron, A. E. (2008). Peptoids that mimic the structure, function, and mechanism of helical antimicrobial peptides. Proceedings of the National Academy of Sciences of the United States of America, 105(8), 2794–2799. 10.1073/pnas.0708254105

Chung, C. T., Niemela, S. L., & Miller, R. H. (1989). One-step preparation of competent Escherichia coli: Transformation and storage of bacterial cells in the same solution. Proceedings of the National Academy of Sciences, 86(7), 2172–2175. 10.1073/pnas.86.7.2172

Costerton, J. W., Stewart, P. S., & Greenberg, E. P. (1999). Bacterial Biofilms: A Common Cause of Persistent Infections. Science, 284(5418), 1318–1322. 10.1126/science.284.5418.1318

Cutler, K. J., Stringer, C., Lo, T. W., Rappez, L., Stroustrup, N., Brook Peterson, S., Wiggins, P. A., & Mougous, J. D. (2022). Omnipose: A high-precision morphology-independent solution for bacterial cell segmentation. Nature Methods, 19(11), 1438–1448. 10.1038/s41592-022-01639-4

Datsenko, K. A., & Wanner, B. L. (2000). One-step inactivation of chromosomal genes in Escherichia coli K-12 using PCR products. *Proceedings of the National Academy of Sciences*, 97(12), 6640–6645. 10.1073/pnas.120163297

Ghosh, A., Kar, R. K., Jana, J., Saha, A., Jana, B., Krishnamoorthy, J., Kumar, D., Ghosh, S., Chatterjee, S., & Bhunia, A. (2014). Indolicidin Targets Duplex DNA: Structural and Mechanistic Insight through a Combination of Spectroscopy and Microscopy. ChemMedChem, 9(9), 2052–2058. 10.1002/cmdc.201402215

Gray, W. T., Govers, S. K., Xiang, Y., Parry, B. R., Campos, M., Kim, S., & Jacobs-Wagner, C. (2019). Nucleoid Size Scaling and Intracellular Organization of Translation across Bacteria. Cell, 177(6), 1632–1648.e20. 10.1016/j.cell.2019.05.017

Hale, J. D., & Hancock, R. E. (2007). Alternative mechanisms of action of cationic antimicrobial peptides on bacteria. Expert Review of Anti-Infective Therapy, 5(6), 951–959. 10.1586/14787210.5.6.951

Hancock, R. E. (1997). Peptide antibiotics. The Lancet, 349(9049), 418–422. 10.1016/S0140-6736(97)80051-7

Hancock, R. E. W., Haney, E. F., & Gill, E. E. (2016). The immunology of host defence peptides: Beyond antimicrobial activity. Nature Reviews Immunology, 16(5), 321–334. 10.1038/nri.2016.29

Hancock, R. E. W., & Sahl, H.-G. (2006). Antimicrobial and host-defense peptides as new anti-infective therapeutic strategies. Nature Biotechnology, 24(12), 1551–1557. 10.1038/nbt1267

Harris, C. R., Millman, K. J., van der Walt, S. J., Gommers, R., Virtanen, P., Cournapeau, D., Wieser, E., Taylor, J., Berg, S., Smith, N. J., Kern, R., Picus, M., Hoyer, S., van Kerkwijk, M. H., Brett, M., Haldane, A., del Río, J. F., Wiebe, M., Peterson, P.,…Oliphant, T. E. (2020). Array programming with NumPy. Nature, 585(7825), 357–362. 10.1038/s41586-020-2649-2

Hebisch, E., Knebel, J., Landsberg, J., Frey, E., & Leisner, M. (2013). High Variation of Fluorescence Protein Maturation Times in Closely Related Escherichia coli Strains. PLoS ONE, 8(10), e75991. 10.1371/journal.pone.0075991

Hsu, C.-H. (2005). Structural and DNA-binding studies on the bovine antimicrobial peptide, indolicidin: Evidence for multiple conformations involved in binding to membranes and DNA. Nucleic Acids Research, 33(13), 4053–4064. 10.1093/nar/gki725

Huang, W., Baliga, C., Aleksandrova, E. V., Atkinson, G., Polikanov, Y. S., Vázquez-Laslop, N., & Mankin, A. S. (2024). Activity, structure, and diversity of Type II proline-rich antimicrobial peptides from insects. EMBO Reports, 25(11), 5194–5211. 10.1038/s44319-024-00277-5

Hunter, J. D. (2007). Matplotlib: A 2D Graphics Environment. Computing in Science & Engineering, 9(3), 90–95. Computing in Science & Engineering. 10.1109/MCSE.2007.55

Kostakioti, M., Hadjifrangiskou, M., & Hultgren, S. J. (2013). Bacterial Biofilms: Development, Dispersal, and Therapeutic Strategies in the Dawn of the Postantibiotic Era. Cold Spring Harbor Perspectives in Medicine, 3(4), a010306–a010306. 10.1101/cshperspect.a010306

Krizsan, A., Knappe, D., & Hoffmann, R. (2015). Influence of the yjiL-mdtM gene cluster on the antibacterial activity of proline-rich antimicrobial peptides overcoming Escherichia coli resistance induced by the missing SbmA transporter system. Antimicrobial Agents and Chemotherapy, 59(10), 5992–5998. 10.1128/AAC.01307-15

Kyte, J., & Doolittle, R. F. (1982). A simple method for displaying the hydropathic character of a protein. Journal of Molecular Biology, 157(1), 105–132. 10.1016/0022-2836(82)90515-0

Lambert, G., & Kussell, E. (2014). Memory and Fitness Optimization of Bacteria under Fluctuating Environments. PLoS Genetics, 10(9), e1004556. 10.1371/journal.pgen.1004556

Le, C.-F., Fang, C.-M., & Sekaran, S. D. (2017). Intracellular Targeting Mechanisms by Antimicrobial Peptides. Antimicrobial Agents and Chemotherapy, 61(4), 1–16.

Lewis, K. (2007). Persister cells, dormancy and infectious disease. Nature Reviews Microbiology, 5(1), 48–56. 10.1038/nrmicro1557

Lewis, P. J., Thaker, S. D., & Errington, J. (2000). Compartmentalization of transcription and translation in Bacillus subtilis. The EMBO Journal, 19(4), 710–718. 10.1093/emboj/19.4.710

Lin, W.-H., & Jacobs-Wagner, C. (2022). Connecting single-cell ATP dynamics to overflow metabolism, cell growth, and the cell cycle in Escherichia coli. Current Biology, 32(18), 3911–3924.e4. 10.1016/j.cub.2022.07.035

Linnik, D., Maslov, I., Punter, C. M., & Poolman, B. (2024). Dynamic structure of E. coli cytoplasm: Supramolecular complexes and cell aging impact spatial distribution and mobility of proteins. Communications Biology, 7(1), 508. 10.1038/s42003-024-06216-3

Liu, H., Song, Z., Zhang, Y., Wu, B., Chen, D., Zhou, Z., Zhang, H., Li, S., Feng, X., Huang, J., & Wang, H. (2025). De novo design of self-assembling peptides with antimicrobial activity guided by deep learning. Nature Materials, 1–12. 10.1038/s41563-025-02164-3

Mardirossian, M., Pérébaskine, N., Benincasa, M., Gambato, S., Hofmann, S., Huter, P., Müller, C., Hilpert, K., Innis, C. A., Tossi, A., & Wilson, D. N. (2018). The Dolphin Proline-Rich Antimicrobial Peptide Tur1A Inhibits Protein Synthesis by Targeting the Bacterial Ribosome. Cell Chemical Biology, 25(5), 530–539.e7. 10.1016/j.chembiol.2018.02.004

Mattiuzzo, M., Bandiera, A., Gennaro, R., Benincasa, M., Pacor, S., Antcheva, N., & Scocchi, M. (2007). Role of the *Escherichia coli* SbmA in the antimicrobial activity of proline-rich peptides. Molecular Microbiology, 66(1), 151–163. 10.1111/j.1365-2958.2007.05903.x

McKinney, W. (2010). Data Structures for Statistical Computing in Python. Proceedings of the 9th Python in Science Conference, 56–61. 10.25080/Majora-92bf1922-00a

Megerle, J. A., Fritz, G., Gerland, U., Jung, K., & Rädler, J. O. (2008). Timing and Dynamics of Single Cell Gene Expression in the Arabinose Utilization System. Biophysical Journal, 95(4), 2103–2115. 10.1529/biophysj.107.127191

Melo, M. N., Ferre, R., & Castanho, M. A. R. B. (2009). Antimicrobial peptides: Linking partition, activity and high membrane-bound concentrations. Nature Reviews Microbiology, 7(3), 245–250. 10.1038/nrmicro2095

Mirdita, M., Schütze, K., Moriwaki, Y., Heo, L., Ovchinnikov, S., & Steinegger, M. (2022). ColabFold: making protein folding accessible to all. Nature Methods, 19(6), 679–682. 10.1038/s41592-022-01488-1

Mookherjee, N., Anderson, M. A., Haagsman, H. P., & Davidson, D. J. (2020). Antimicrobial host defence peptides: Functions and clinical potential. Nature Reviews Drug Discovery, 19(5), 311–332. 10.1038/s41573-019-0058-8

Naves, P., Del Prado, G., Huelves, L., Gracia, M., Ruiz, V., Blanco, J., Dahbi, G., Blanco, M., Del Carmen Ponte, M., & Soriano, F. (2008). Correlation between virulence factors and in vitro biofilm formation by Escherichia coli strains. Microbial Pathogenesis, 45(2), 86–91. 10.1016/j.micpath.2008.03.003

Okuta, R., Unno, Y., Nishino, D., Hido, S., & Loomis, C. (2017). CuPy: A NumPy-Compatible Library for NVIDIA GPU Calculations. 31st Conference on Neural Information Processing Systems (NIPS 2017).

Papagiannakis, A., Yu, Q., Govers, S. K., Lin, W.-H., Wingreen, N. S., & Jacobs-Wagner, C. (2025). Nonequilibrium polysome dynamics promote chromosome segregation and its coupling to cell growth in Escherichia coli. eLife, 14, RP104276. 10.7554/eLife.104276

Paszke, A., Gross, S., Massa, F., Lerer, A., Bradbury, J., Chanan, G., Killeen, T., Lin, Z., Gimelshein, N., Antiga, L., Desmaison, A., Köpf, A., Yang, E., DeVito, Z., Raison, M., Tejani, A., Chilamkurthy, S., Steiner, B., Fang, L.,…Chintala, S. (2019). PyTorch: An imperative style, high-performance deep learning library. In Proceedings of the 33rd International Conference on Neural Information Processing Systems (pp. 8026–8037). Curran Associates Inc.

Paulsen, V. S., Blencke, H.-M., Benincasa, M., Haug, T., Eksteen, J. J., Styrvold, O. B., Scocchi, M., & Stensvåg, K. (2013). Structure-Activity Relationships of the Antimicrobial Peptide Arasin 1— And Mode of Action Studies of the N-Terminal, Proline-Rich Region. PLOS ONE, 8(1), e53326. 10.1371/journal.pone.0053326

Pedregosa, F., Varoquaux, G., Gramfort, A., Michel, V., Thirion, B., Grisel, O., Blondel, M., Prettenhofer, P., Weiss, R., Dubourg, V., Vanderplas, J., Passos, A., Cournapeau, D., Brucher, M., Perrot, M., & Duchesnay, É. (2011). Scikit-learn: Machine Learning in Python. J. Mach. Learn. Res., 12(null), 2825–2830.

Podda, E., Benincasa, M., Pacor, S., Micali, F., Mattiuzzo, M., Gennaro, R., & Scocchi, M. (2006). Dual mode of action of Bac7, a proline-rich antibacterial peptide. Biochimica Et Biophysica Acta, 1760(11), 1732–1740. 10.1016/j.bbagen.2006.09.006

Pränting, M., Negrea, A., Rhen, M., & Andersson, D. I. (2008). Mechanism and Fitness Costs of PR-39 Resistance in *Salmonella enterica* Serovar Typhimurium LT2. Antimicrobial Agents and Chemotherapy, 52(8), 2734–2741. 10.1128/AAC.00205-08

Roth, B. L., Poot, M., Yue, S. T., & Millard, P. J. (1997). Bacterial viability and antibiotic susceptibility testing with SYTOX green nucleic acid stain. *Applied and Environmental Microbiology*, 63(6), 2421–2431. 10.1128/aem.63.6.2421-2431.1997

Sanamrad, A., Persson, F., Lundius, E. G., Fange, D., Gynnå, A. H., & Elf, J. (2014). Single-particle tracking reveals that free ribosomal subunits are not excluded from the *Escherichia coli* nucleoid. Proceedings of the National Academy of Sciences, 111(31), 11413–11418. 10.1073/pnas.1411558111

Savitzky, Abraham., & Golay, M. J. E. (1964). Smoothing and Differentiation of Data by Simplified Least Squares Procedures. Analytical Chemistry, 36(8), 1627–1639. 10.1021/ac60214a047

Scocchi, M., Lüthy, C., Decarli, P., Mignogna, G., Christen, P., & Gennaro, R. (2009). The Proline-rich Antibacterial Peptide Bac7 Binds to and Inhibits in vitro the Molecular Chaperone DnaK. International Journal of Peptide Research and Therapeutics, 15(2), 147–155. 10.1007/s10989-009-9182-3

Scocchi, M., Tossi, A., & Gennaro, R. (2011). Proline-rich antimicrobial peptides: Converging to a non-lytic mechanism of action. Cellular and Molecular Life Sciences, 68(13), 2317–2330. 10.1007/s00018-011-0721-7

Schindelin, J., Arganda-Carreras, I., Frise, E., Kaynig, V., Longair, M., Pietzsch, T., Preibisch, S., Rueden, C., Saalfeld, S., Schmid, B., Tinevez, J.-Y., White, D. J., Hartenstein, V., Eliceiri, K., Tomancak, P., & Cardona, A. (2012). Fiji: an open-source platform for biological-image analysis. Nature Methods, 9(7), 676–682. 10.1038/nmeth.2019

Sharan, S. K., Thomason, L. C., Kuznetsov, S. G., & Court, D. L. (2009). Recombineering: A homologous recombination-based method of genetic engineering. Nature Protocols, 4(2), 206–223. 10.1038/nprot.2008.227

Sneideris, T., Erkamp, N. A., Ausserwöger, H., Saar, K. L., Welsh, T. J., Qian, D., Katsuya-Gaviria, K., Johncock, M. L. L. Y., Krainer, G., Borodavka, A., & Knowles, T. P. J. (2023). Targeting nucleic acid phase transitions as a mechanism of action for antimicrobial peptides. Nature Communications, 14(1), 7170. 10.1038/s41467-023-42374-4

Snoussi, M., Talledo, J. P., Del Rosario, N. A., Mohammadi, S., Ha, B. Y., Košmrlj, A., & Taheri-Araghi, S. (2018). Heterogeneous absorption of antimicrobial peptide LL37 in escherichia coli cells enhances population survivability. eLife, 7, 1–21. 10.7554/eLife.38174

Stylianidou, S., Brennan, C., Nissen, S. B., Kuwada, N. J., & Wiggins, P. A. (2016). SuperSegger: Robust image segmentation, analysis and lineage tracking of bacterial cells. Molecular Microbiology, 102(4), 690–700. 10.1111/mmi.13486

Subach, O. M., Cranfill, P. J., Davidson, M. W., & Verkhusha, V. V. (2011). An Enhanced Monomeric Blue Fluorescent Protein with the High Chemical Stability of the Chromophore. PLoS ONE, 6(12), e28674. 10.1371/journal.pone.0028674

Subbalakshmi, C., & Sitaram, N. (1998). Mechanism of antimicrobial action of indolicidin. FEMS Microbiology Letters, 160(1), 91–96. 10.1111/j.1574-6968.1998.tb12896.x

Taheri-Araghi, S. (2024). Synergistic action of antimicrobial peptides and antibiotics: Current understanding and future directions. Frontiers in Microbiology, 15, 1390765. 10.3389/fmicb.2024.1390765

Thappeta, Y., Cañas-Duarte, S. J., Wang, H., Kallem, T., Fragasso, A., Xiang, Y., Gray, W., Lee, C., Hardo, G., Cegelski, L., & Jacobs-Wagner, C. (2025). Glycogen phase-separation drives macromolecular rearrangement and asymmetric division in E. coli. The EMBO Journal, 44(24), 7434–7476. 10.1038/s44318-025-00621-y

Thomason, L. C., Costantino, N., & Court, D. L. (2007). E. coli genome manipulation by P1 transduction. Current Protocols in Molecular Biology, Chapter 1, 1.17.1–1.17.8. 10.1002/0471142727.mb0117s79

Virtanen, P., Gommers, R., Oliphant, T. E., Haberland, M., Reddy, T., Cournapeau, D., Burovski, E., Peterson, P., Weckesser, W., Bright, J., van der Walt, S. J., Brett, M., Wilson, J., Millman, K. J., Mayorov, N., Nelson, A. R. J., Jones, E., Kern, R., Larson, E.,…van Mulbregt, P. (2020). SciPy 1.0: Fundamental algorithms for scientific computing in Python. Nature Methods, 17(3), 261–272. 10.1038/s41592-019-0686-2

Walt, S. van der, Schönberger, J. L., Nunez-Iglesias, J., Boulogne, F., Warner, J. D., Yager, N., Gouillart, E., & Yu, T. (2014). scikit-image: Image processing in Python. PeerJ, 2, e453. 10.7717/peerj.453

Wan, F., Wong, F., Collins, J. J., & De La Fuente-Nunez, C. (2024). Machine learning for antimicrobial peptide identification and design. Nature Reviews Bioengineering, 2(5), 392–407. 10.1038/s44222-024-00152-x

Wang, G., Li, X., & Wang, Z. (2016). APD3: The antimicrobial peptide database as a tool for research and education. Nucleic Acids Research, 44(D1), D1087–D1093. 10.1093/nar/gkv1278

Wang, G., Zietz, C. M., Mudgapalli, A., Wang, S., & Wang, Z. (2022). The evolution of the antimicrobial peptide database over 18 years: Milestones and new features. Protein Science, 31(1), 92–106. 10.1002/pro.4185

Waskom, M. L. (2021). seaborn: Statistical data visualization. Journal of Open Source Software, 6(60), 3021. 10.21105/joss.03021

Wernersson, E., Gelali, E., Girelli, G., Wang, S., Castillo, D., Mattsson Langseth, C., Verron, Q., Nguyen, H. Q., Chattoraj, S., Martinez Casals, A., Blom, H., Lundberg, E., Nilsson, M., Marti-Renom, M. A., Wu, C.-t., Crosetto, N., & Bienko, M. (2024). Deconwolf enables high-performance deconvolution of widefield fluorescence microscopy images. Nature Methods, 21(7), 1245–1256. 10.1038/s41592-024-02294-7

Wimley, W. C., & Hristova, K. (2011). Antimicrobial peptides: Successes, challenges and unanswered questions. Journal of Membrane Biology, 239(1–2), 27–34. 10.1007/s00232-011-9343-0

Xiang, Y., Surovtsev, I. V., Chang, Y., Govers, S. K., Parry, B. R., Liu, J., & Jacobs-Wagner, C. (2021). Interconnecting solvent quality, transcription, and chromosome folding in Escherichia coli. Cell, 184(14), 3626--3642.e14. 10.1016/j.cell.2021.05.037

Yonezawa, A., Kuwahara, J., Fujii, N., & Sugiura, Y. (1992). Binding of tachyplesin I to DNA revealed by footprinting analysis: Significant contribution of secondary structure to DNA binding and implication for biological action. Biochemistry, 31(11), 2998–3004. 10.1021/bi00126a022

Zhu, Y., Mohapatra, S., & Weisshaar, J. C. (2019). Rigidification of the Escherichia coli cytoplasm by the human antimicrobial peptide LL-37 revealed by superresolution fluorescence microscopy. Proceedings of the National Academy of Sciences of the United States of America, 116(3), 1017–1026. 10.1073/pnas.1814924116

